# Redox Conduction Through Cytochrome ‘Nanowires’ Can Sustain Cellular Respiration

**DOI:** 10.1101/2024.04.03.587941

**Authors:** Matthew J. Guberman-Pfeffer

## Abstract

Micron-scale electron transfer through polymeric cytochrome ‘nanowires’ powers prokaryotic life from hydrothermal vents to terrestrial soils in ways not fully understood. Herein, six reduction potentials from recently reported spectroelectrochemistry are each assigned with <0.04 eV to the cryogenic electron microscopy structure of the hexa-heme homopolymeric outer-membrane cytochrome type S (OmcS) from *Geobacter sulfurreducens* using hybrid quantum/classical computations. The unambiguous assignments define a reversible free energy ‘roller-coaster’ that is dynamically modulated by <0.1 V under the flow of electrons due to redox cooperativities between adjacent hemes. A physiologically relevant tens to hundreds of filaments are predicted to suffice for cellular respiration by pairing, in the context of non-adiabatic Marcus theory, the free energy landscape with reorganization energies that account for active site or protein-water electronic polarizability, and electronic couplings characteristic of the highly conserved heme packing motifs. General considerations on protein electron transfer and comparison to all known cytochrome ‘nanowires’ suggest the mechanistic insights are broadly applicable to multi-heme cytochromes in all kingdoms of life.

## 1. Introduction

Prokaryotes from hydrothermal vents to terrestrial soils can exhale ∼10^6^ electrons/s/cell (∼100 fA/cell)^1-7^ through filamentous cytochrome ‘nanowires’^8^ that electrify microbial communities^9^ and biotic-abiotic interfaces.^10^ This ‘rock breathing’ strategy for anaerobic life, known as extracellular electron transfer, is ancient,^11^ ubiquitous,^12, 13^ environmentally significant,^14-17^ and holds promise for sustainable technologies,^18-26^ but only if its mechanistic underpinnings are elucidated. A fundamental question beyond the limits of current experimental techniques^8, 27-29^ that is addressed herein by hybrid quantum/classical computations is: What is the electron transfer mechanism that connects the atomic structures of filamentous cytochromes to their physiological role as mesoscopic electrical conductors?

All known cytochrome filaments^8, 30-33^ have the basic anatomy summarized in Figure 1. A single type of cofactor—a bis-histidine ligated *c*-type heme—is repeated hundreds of times and stacked in highly conserved geometries^8^ to form a micron-long (linear or branched) spiraling chain, which is encased by a mostly unstructured (≥50% turns and loops) protein sheath. Each filament is constructed of a different Lego-like multi-heme cytochrome that polymerizes either exclusively through non-covalent interactions, or one or more coordination bonds between protomers. These filaments attest to the great lengths, biosynthetically and literally, life will go to breathe.

**Figure 1.**
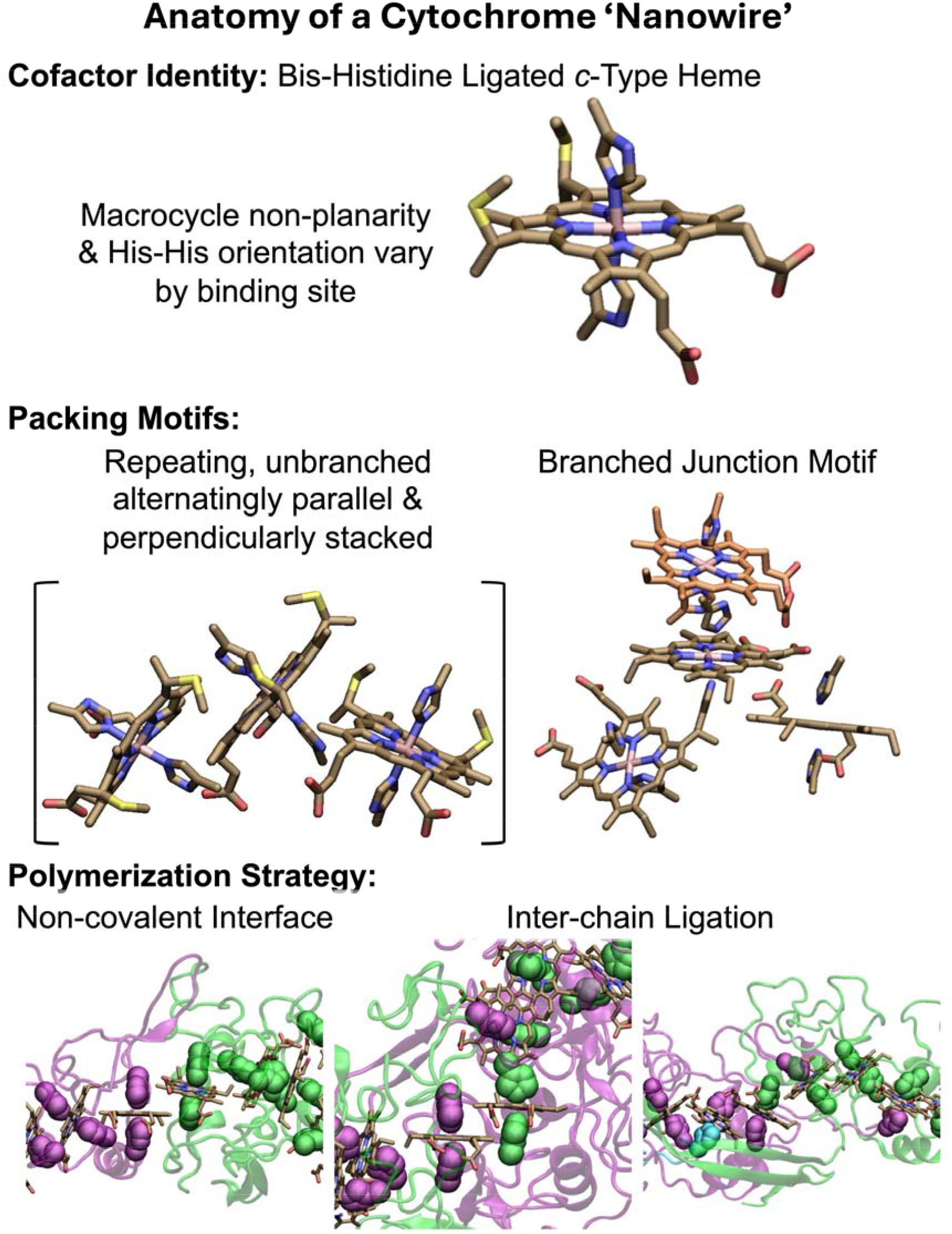
Anatomy of a cytochrome ‘nanowire.’ The heme cofactor, its packing motifs, and filament polymerization strategies are shown. Purple, green, and cyan indicate different protein chains. His ligands are shown in vdW representation. A heme that is coordinated by two differently colored His ligands receives one of them from the adjacent protein chain. The subunit interfaces shown at the bottom from left to right come from OmcZ (PDB 7LQ5),^33^ OmcS (PDB 6EF8),^31^ and F2KMU8 (PDB 8E5G).^8^ The figure was designed with VMD version 1.9.4a57.^34^

The first-discovered and most extensively characterized representative of the structural class to date is the outer-membrane cytochrome type S (OmcS) from *Geobacter sulfurreducens*.^30, 31, 35^ OmcS is a hexa-heme cytochrome that has only ∼19% regular 2° structure and polymerizes by donating a coordinating His ligand to the penultimate heme of another OmcS molecule.^31^

Theoretical models spanning the entire continuum of electron transfer theory^35-41^ have been proposed to describe the function of OmcS since its structure was resolved in 2019.^30, 31^ None of these models, however, have successfully linked the atomistic structure to a functional characterization or physiological role. Regardless of whether micron-scale electron transfer is pictured as a relay of reduction-oxidation (redox) reactions between individual (purely incoherent transfer) or blocks (coherence-assisted transfer) of hemes,^35-39^ or quantum transport through electronic bands,^40-42^ the models have not explained the high electronic conductivity (30 mS/cm) measured by atomic force microscopy (AFM);^35^ a discrepancy all the more striking since OmcS is 10^3^-fold less conductive than another cytochrome filament by the same technique.^43^

The discrepancy is customarily interpreted as a failure of theory.^36, 40^ Concerns recently raised about structural and electronic artifacts of the experimental conditions,^27^ however, may suggest the comparison itself is inappropriate. By instead comparing to the metabolic flux of electrons from a *G. sulfurreducens* cell, it is herein demonstrated that a relay of redox reactions is physiologically sufficient if, as observed^44^ or anticipated by bioenergetic calculations,^45^ tends to hundreds of filament/cell (once thought to be pili but now argued to be OmcS)^31^ are expressed.

The redox conductivity model is predicated on quantum mechanical/molecular mechanical computations at molecular dynamics generated configurations (QM/MM@MD)^38^ that are shown to have predicted within 0.04 eV each of the six redox potentials of the hemes in OmcS 16-months before they were reported by spectroelectrochemistry. The unparalleled theory-experiment comparison enables an integrated view of structural, spectroelectrochemical, and microbiological observations.

The analysis is distinguished from prior modeling efforts^35-37, 41^ by: (1) Comparing solution-phase computations of the redox potentials based on the CryoEM structure to solution-phase spectroelectrochemical measurements that are thought to be physiologically relevant^29^ (Sections 2.2 and 2.3); (2) Quantifying heme-heme redox cooperativities with the recently reported, validated, and one of the only known computational methods for this purpose^46, 47^ (Section 2.4); (3) Predicting an assignment of the spectroelectrochemical potentials to the hemes in the CryoEM structure (Section 2.5); (4) Systematically evaluating the influence of active site and environment electronic polarizability on filament conductivity (Sections 2.6 and 2.9); (5) Comparing the computed rates to experimental estimates for other multi-heme proteins with the same heme-packing geometries (Section 2.8); and (6) Comparing the predicted conductivity to the cellular respiratory rate instead of abiological solid-state electrical measurements^27^ (Section 2.9).

Along the way, the fundamental conclusion that a bioenergetically manageable and experimentally observed tens-to-hundreds of cytochrome filaments/cell can more than adequately discharge the cellular respiratory current is shown to be insensitive to either the proposed assignment of redox potentials to the CryoEM structure or the degree to which electronic polarization lowers electron transfer activation barriers. General considerations on biological electron transfer suggest that the method, model, and mechanistic insights are generally applicable to multi-heme chains in all kingdoms of life.

## 2. Results and Discussion

### 2.1. Overview

The analysis begins with a comparison of reported and computed redox titration curves, and a justification of the approximation needed to propose a molecular-level interpretation of the data (Sections 2.2 to 2.3), including the quantification of heme-heme interaction energies (Section 2.4). An assignment of spectroelectrochemical potentials to the CryoEM structure is proposed in Section 2.5 based on the prior experimental conclusion of single-heme-dominated transitions^29^ and the <0.04 V agreement with computations for each of six such modeled redox transitions. The mapping of redox potentials to the hemes of the CryoEM structure enables the definition of a free energy landscape, although results are also presented throughout the article for the 718 other possible landscapes given different assignments of the potentials to the hemes in the structure. The other ingredient of the electron transfer activation barrier needed to model filament conductivity, namely, reorganization energy, is considered in Section 2.6 with specific attention to the debated role of active site versus environmental electronic polarization.^48-52^ Section 2.7 and 2.8 discuss the resulting activation barriers and predicted electron transfer rates relative to experimental estimates. Section 2.9 finally presents the predicted range of charge diffusion constants based on the computed rates and compares the value to both the demands of cellular physiology and prior electrical measurements.

Overall, the analysis shows that if the filaments expressed *by G. sulfurreducens* have the CryoEM-resolved structures under physiological conditions and function as redox conductors in the limit of non-adiabatic Marcus theory, a bioenergetically manageable^45^ and experimentally estimated^44^ tens-to-hundreds of cytochrome filaments are more than sufficient to discharge the cellular metabolic current.

### 2.2. Computations Accurately Predict the Measured Spectroelectrochemical Titration Curve

To assess whether the picture of consecutive redox reactions is mechanistically viable for micron-scale electron transfer before resorting to competing alternatives like coherence-assisted charge hopping^36^ or decoherent quantum transport,^40, 41^ a faithful reproduction of the redox properties of OmcS is mandatory. Table 1 therefore compares, for the first time, all previously published (computed^35, 38^ and measured^29^) redox potentials of OmcS, and Figure 2 shows the corresponding redox titration profiles. The subsequent interpretation of this data in terms of the molecular structure shown alongside it in the figure is one of the unique contributions of the present article.

**Table 1.** Comparison of spectroelectrochemically-derived and computed redox potentials. All values are in V vs. Standard Hydrogen Electrode (SHE). The parenthetical numbers indicate the heme to which the computed potential belongs in the simulations. The spectroelectrochemical data is reproduced from Ref. 29. The computed potentials from Guberman-Pfeffer are reproduced from Ref. 38; those from Dahl *et al*. are reproduced from Ref. 35. 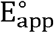 is the apparent macroscopic midpoint potential for OmcS.

| Index | Experiment | Cathodic Direction |  | Anodic Direction |
| --- | --- | --- | --- | --- |
|  |  | Guberman-Pfeffer<br>QM/MM@MD | Dahl *et al.*<br>QM/MM@MD | Dahl *et al.*<br>QM/MM@MD |
| 1 | -0.027 | $-0.063 \pm 0.034$ (#1) | $-0.080 \pm 0.06$ (#6) | $-0.421 \pm 0.05$ (#2) |
| 2 | -0.082 | $-0.071 \pm 0.027$ (#3) | $-0.108 \pm 0.04$ (#2) | $-0.431 \pm 0.06$ (#6) |
| 3 | -0.108 | $-0.095 \pm 0.031$ (#6) | $-0.126 \pm 0.05$ (#3) | $-0.491 \pm 0.05$ (#1) |
| 4 | -0.158 | $-0.111 \pm 0.027$ (#2) | $-0.191 \pm 0.05$ (#1) | $-0.647 \pm 0.05$ (#4) |
| 5 | -0.172 | $-0.177 \pm 0.024$ (#5) | $-0.334 \pm 0.04$ (#4) | $-0.727 \pm 0.05$ (#3) |
| 6 | -0.230 | $-0.248 \pm 0.036$ (#4) | $-0.521 \pm 0.05$ (#5) | $-0.826 \pm 0.05$ (#5) |
| $E_{\text{app}}^{\circ}$ | -0.132 | -0.116 | -0.168 | -0.572 |

**Figure 2.**
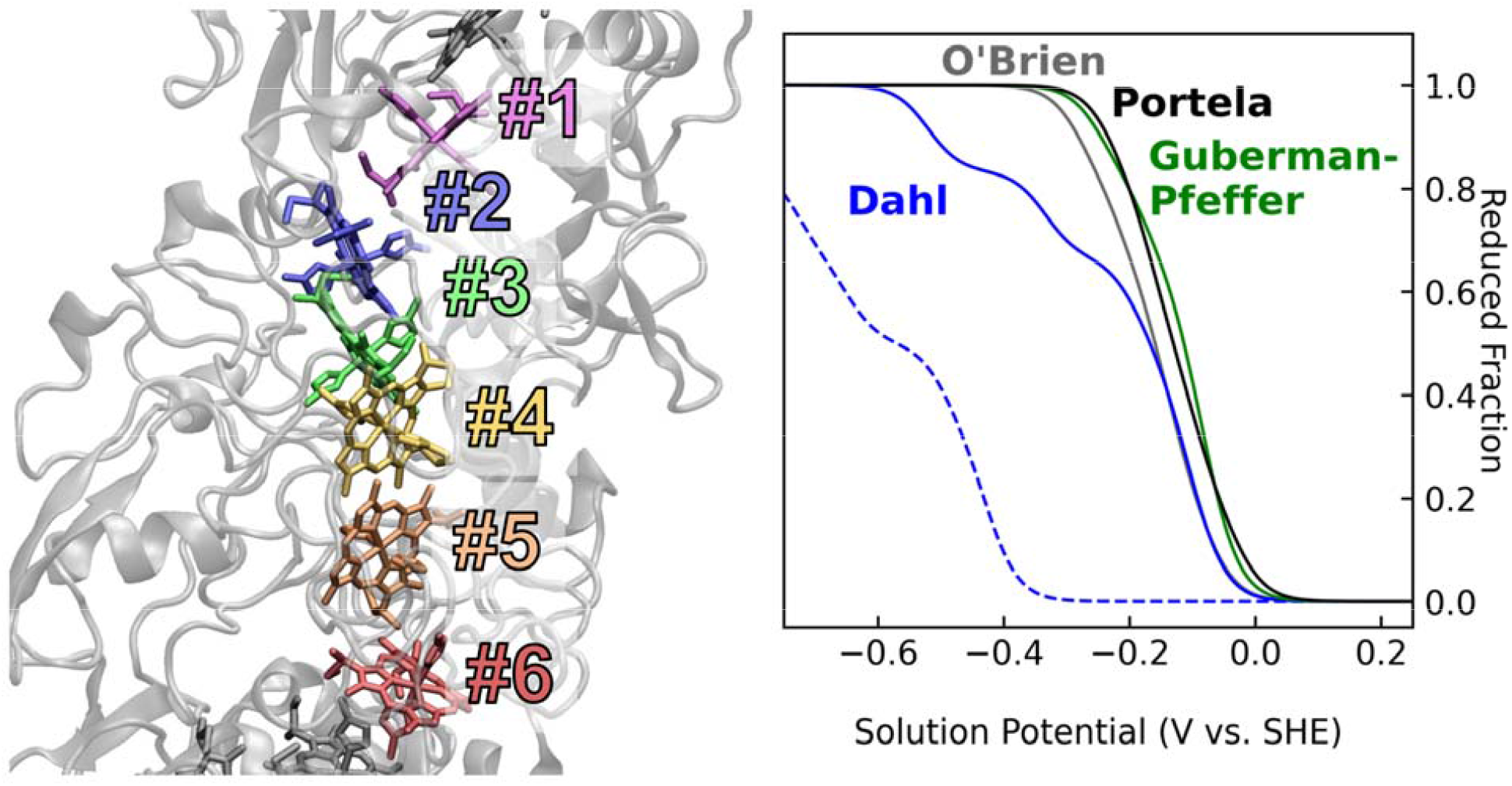
Starting from only the structure of OmcS, quantum mechanical/molecular mechanical computations at molecular dynamics-generated configurations (QM/MM@MD) accurately reproduce the experimental spectroelectrochemical titration curve. The structure is from PDB 6EF8.^31^ The six hemes in the N-to C-terminal direction are numbered sequentially and colored purple, blue, yellow, orange, green and red. The experimental titration curves in gray and black are respectively reproduced from O’Brien^53^ and Portela *et al*.^29^ The computed titration curves are simulated from redox potentials reported by Dahl *et al*.^35^ and Guberman-Pfeffer.^38^ Note that the experimental finding of O’Brien was published >496 days *before* the computational prediction by Dahl *et al*. from the same laboratory.

The figure shows the fraction of reduced hemes in the OmcS filament as a function of the solution potential. The gray, blue, green and black solid-line titration curves are respectively and chronologically the redox profiles spectroelectrochemically measured by O’Brien,^53^ computationally predicted by Dahl *et al*.,^35^ Guberman-Pfeffer,^38^ and measured using spectroelectrochemistry by Portela *et al*.^29^ The computed solid-line titration curves represent an approximation to a cathodic electrochemical sweep in which each heme was separately reduced while all other hemes were held in the oxidized state. The blue dashed-line titration curve represents an anodic electrochemical sweep simulated by Dahl *et al*.^35^ in which each heme was separately oxidized while all other hemes were fixed in the reduced state.

Portela *et al*.^29^ reported that the experimental (black) titration curve is well fit by a sum of Nernst equations for six independent or non-interacting heme cofactors. Consistent with this conclusion, a sum of six Nernst equations using the redox potentials computed with QM/MM@MD techniques by Guberman-Pfefer gives a titration curve that virtually superimposes on the experimental profile. The experimental and computed midpoint potentials are -0.132 and -0.114 V vs. Standard Hydrogen Electrode (SHE), respectively.

A sum of Nernst equations using the six redox potentials computed by Dahl *et al*. gives significantly worse agreement. The simulated cathodic trace has a reasonable midpoint potential (−0.168 V vs. SHE), but distorted shape at strongly reducing voltages because of predicted redox potentials of -0.334 and -0.521 V vs. SHE (Table 1), which are up to 0.291 V more negative than the potential for any redox transition reported by Portela *et al*.^29^

The titration curve predicted using the redox potentials of Dahl *et al*. for an anodic electrochemical sweep is in even worse agreement than the cathodic trace. The predicted midpoint potential of -0.572 V vs. SHE is shifted by -0.404 V relative to the simulated cathodic trace. This enormous and erroneously predicted hysteresis by Dahl *et al*. (Table 1; Figure 2) was previously used to explain why OmcS is six-fold more conductive in the fully reduced versus fully oxidized state, an observation that itself is inconsistent with the nature of a cytochrome. Redox conduction through a cytochrome is maximal when there is an equal proportion of reduced (electron donating) and oxidized (electron accepting) hemes, and rapidly falls to zero when all hemes are either reduced or oxidized.^54^

The conclusion is that the redox potentials predicted by Dahl *et al*. (including the present author), and the entire edifice of the model built upon them to rationalize the experimentally observed conductivity of OmcS must now be unequivocally rejected given the newly published spectroelectrochemical data. Notably, four authors, including a senior investigator on both the Dahl *et al*.^35^ and Portela *et al*.^29^ works chose *not* to comment on the inconsistency with experiment of the predicted redox potentials well in excess of any known for bis-histidine ligated *c*-type hemes^55^ as in OmcS, or the predicted enormous electrochemical hysteresis.

Worse, pages 86 to 90 of a dissertation^53^ published in 2020 by Patrick O’Brien and supervised by a senior investigator on the two-years-later-published Dahl *et al*. work already showed no redox potential in OmcS in excess of -0.280 V and no substantial electrochemical hysteresis. The findings in the dissertation were reproduced by Catharine Shipps under the supervision of the same investigator and published on pages 39 to 42 of a dissertation in 2022.^56^ The *timeline of the published record* clearly indicates that the senior investigator published computational predictions that were already invalidated by two sets of independently generated data available within their lab but were not disclosed publicly (O’Brien’s dissertation was embargoed at the time). Noting such conduct is of value to the community.

### 2.3. Spectroelectrochemical Potentials are Unambiguously Assigned to the CryoEM structure Using a Computational ‘Rosette Stone’

Further insight is possible by approximating the spectroelectrochemical titration curve in Figure 2 by a sum of Nernst equations to obtain the potentials for discrete redox transitions that were too finely spaced to be resolved experimentally.^57^ There must be as many one-electron Nernst equations in the sum as there are one-electron redox centers in the protein. A subunit of OmcS has six one-electron donating/accepting heme groups. If the redox profile of a subunit does not depend on its position in the helical pitch of the filament (∼4 subunits/pitch), a minimum of six Nernst equations is needed to approximate the titration curve in Figure 2.

Each of these Nernst equations describes a transition between macroscopic redox states (macrostates) in which OmcS gains/losses a single reducing equivalent. As shown in Figure 3, each macrostate (S0 – S6) comprises an ensemble of microstates that describes all the permutations for distributing the reducing equivalents among the six hemes at that macroscopic reduction stage. Sequential addition of six electrons takes OmcS from the fully oxidized (S0) to fully reduced (S6) macrostate, with the number of microstates at each reduction stage following Pascale’s triangle (Figure 3). The redox potential of each microstate, weighted by its fractional occupancy among the other microstates for that reduction stage, determines the macroscopic redox potentials obtained by fitting six Nernst equations to the titration curve in Figure 2.

**Figure 3.**
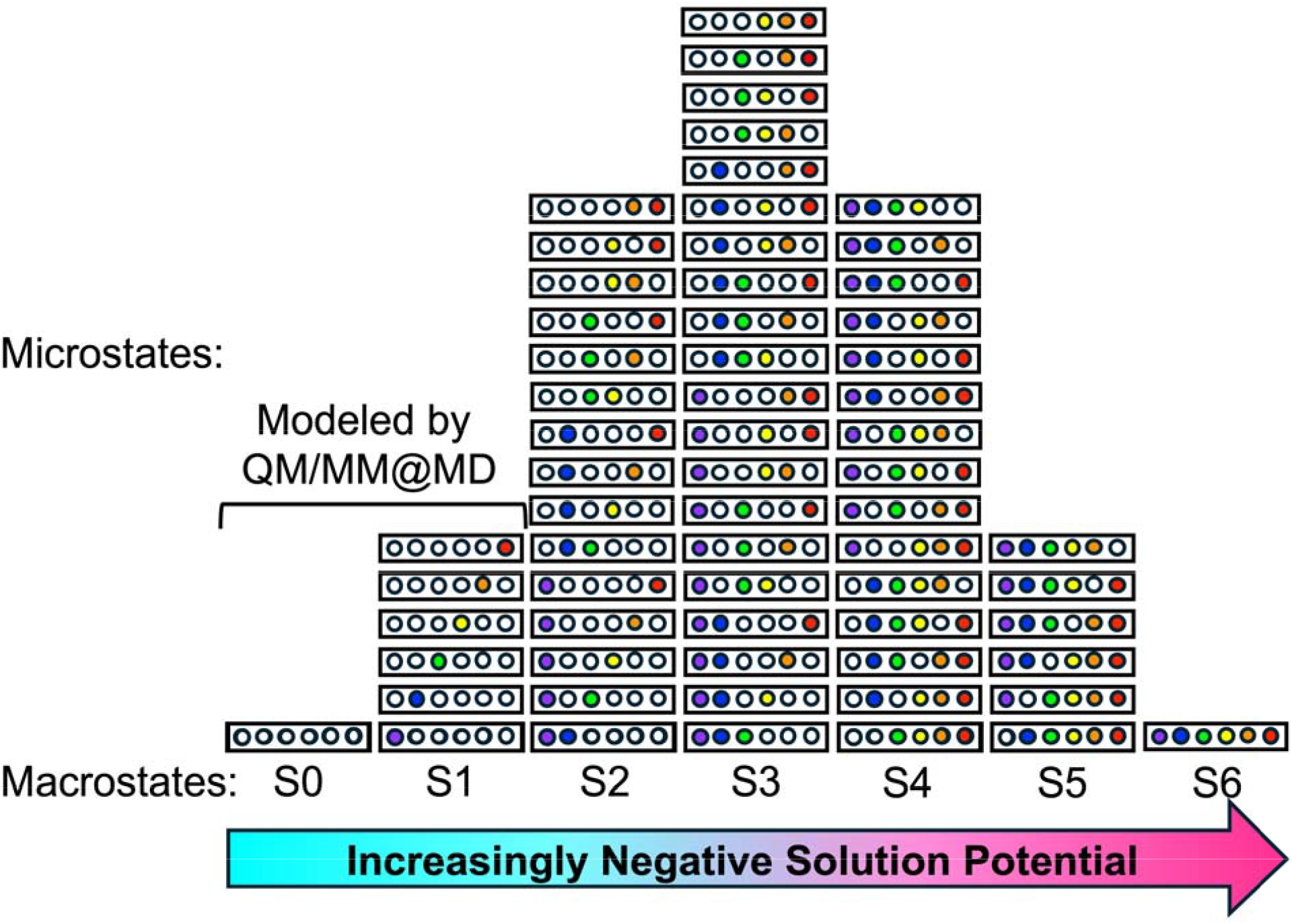
Under increasingly negative solution potentials, OmcS passes from the fully oxidized to fully reduced redox macrostate (S0 S6 columns), with the sequentially added reducing equivalents distributed among the hemes in various microstates (rectangles). Each circle represents a heme. Purple, blue, green, yellow, orange, and red indicate Hemes #1 through #6 as labeled in Figure 2. Note that if the protonation of acid/base sites are coupled to the redox transitions, then the number of microstates shown for a given macrostate would increase by (*i*.*e*., there would be two stacks of microstates for each acid/base site corresponding to when that site is protonated or deprotonated). The case shown is the simplest possible model.

It is computationally intractable to model by QM/MM@MD techniques the 64 microstates shown in Figure 3. For perspective, the computations by Guberman-Pfeffer^38^ analyzed in Figures 2 and 4 involved 1.9 µs of production-stage molecular dynamics followed by >3×10^3^ QM/MM energy evaluations *just* to model the 7 microstates comprising the S0 and S1 macrostates (Figure 3). Fortunately, Portela *et al*.^29^ found little difference between fitting six sequentially-coupled or six independent Nernst equations to the experimental titration curve.^57^ This finding suggests that the potential at which each heme is reduced does not significantly depend on the redox state of any other heme. Transitions between macrostates are then equivalent to transitions between microstates in which only one heme is reduced (*i*.*e*., only the six unique transitions from S0 to S1 in Figure 3 need to be considered). As Portela *et al*. conclude, “The close agreement between the experimental macro- and microscopic potentials suggested that a single heme contributed to each redox transition between the different oxidation stages in OmcS.”^29^

**Figure 4.**
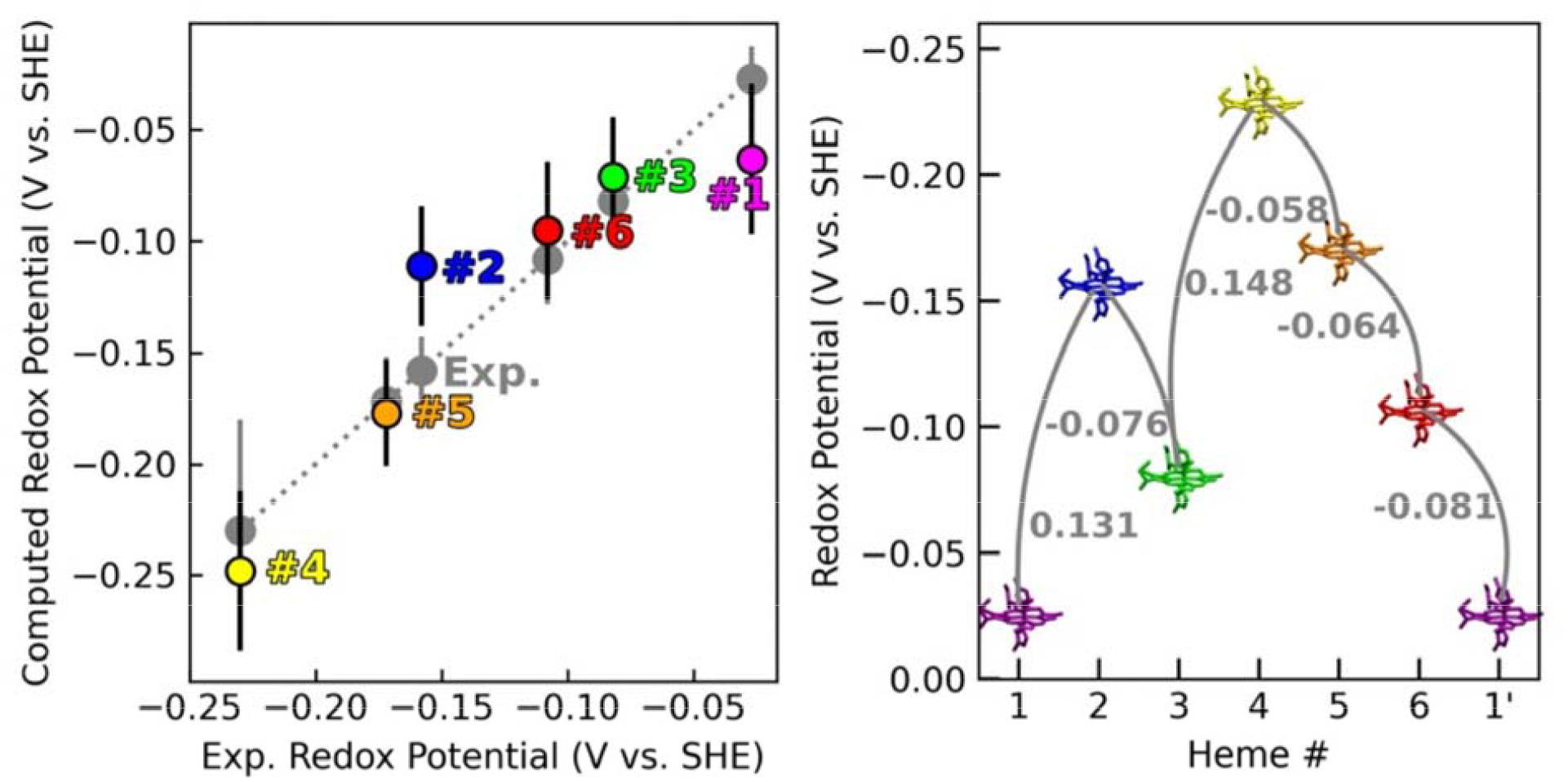
Redox potentials computed by Guberman-Pfeffer agree within 0.036 eV of the spectroelectrochemically measured potentials and define a potential ‘roller-coaster’ along the linear heme sequence in OmcS. Deviations from the diagonal in the computation (Comp.) versus experiment (Exp.) plot at left indicate differences from the experimental value. For the comparison to be non-arbitrary, the computed and measured redox potentials were both sorted in the same way before making a 1:1 mapping. The computed and experimental redox potentials are reproduced, respectively, from Portela *et al*.^*29*^ and Guberman-Pfeffer.^38^ The redox potential landscape at right is plotted from more positive potentials at bottom to more negative potentials on top so ‘uphill’ and ‘downhill’ electron hops (represented by arrows) correspond to the free energy changes indicated in gray. The first heme of the next subunit (labeled “1’”) is included to show the repeating pattern for the homopolymer.

If a single heme dominantly contributes to each redox transition, then the potential of that transition can reasonably be assigned to a specific heme. This independent-heme approximation has also been used in all modeling of redox conductivity in OmcS to date, but critically, it is assessed below for the first time.^37-39^ Thus, direct comparison between the six redox potentials from experiment and theory is possible.

A *non-arbitrary* way to compare the experimentally-derived and computationally-predicted redox potentials is to sort *both sets of potentials by the same rule* (*e*.*g*., most negative to most positive potential). Doing so reveals (Table 1 and Figure 4) that the six independently predicted redox potentials by Guberman-Pfeffer,^38^ starting from *only* the CryoEM structure, each come within ≤0.036 eV (0.8 kcal/mol) of an experimental redox potential, which is only slightly more than thermal energy at 298 K (0.026 eV). The level of agreement is better than can be expected and probably benefits from error cancellation (as do approximate hybrid density functionals by construction for quantum chemical calculations). Of note, however, is the *consistently close* agreement and the fact that the errors are *not systematic*. Three redox potentials are underestimated by 0.013 to 0.047 eV whereas the other three potentials are overestimated by 0.005 to 0.036 eV. These deviations between theory and experiment are no greater than the standard deviation for each technique.

It should be noted that the six-parameter fit used by Portela *et al*. may suffer from the problem of von Neumann’s elephant^58^ (*i*.*e*., there are too many adjustable parameters to give a unique solution). Portela *et al*. do not discuss performing a sensitivity analysis to see how the fit solution depends on the initial guess, or how many different sets of six redox potentials can equivalently describe the titration curve in Figure 2.

Trusting at face-value the analysis of Portela *et al*., however, the consistent agreement between experiment and theory empowers the modeling to serve as a ‘rosette stone’ for mapping the spectroelectrochemical potentials to specific hemes in the CryoEM structure (Figure 4).^59, 60^ The sequence of hemes in order of increasingly positive (measured, computed) potential in volts versus SHE is: Heme #4 (−0.230±0.051, -0.248±0.036) < Heme #5 (−0.172±0.019, -0.177±0.024) < Heme #2 (−0.158±0.010, -0.111±0.027) < Heme #6 (−0.108±0.033, -0.095±0.031) < Heme #3 (−0.082±0.023, -0.071±0.027) < Heme #1 (−0.027±0.017, -0.063±0.034).

It is clear from Figure 4 that the linear sequence of hemes in the OmcS structure does not conform to the thermodynamic sequence of redox potentials. Electrons passing along the heme chain are geometrically constrained to changes in potential of 0.131, -0.076, 0.148, -0.058, -0.064, -0.081 V from Heme #1 of one subunit to Heme #1 in the next subunit. The implications of this potential ‘roller-coaster’ for electron transfer are discussed below. Structural determinants of the redox potentials, which are overwhelmingly electrostatic in origin, have already been delineated by Guberman-Pfeffer.^39^

By contrast, the same experiment-versus-theory comparison (Table 1) reveals that two of the redox potentials predicted by Dahl *et al*. are 0.162 and 0.291 V too far negative. Because the discrepancy is not consistent for all the redox potentials predicted by Dahl *et al*., the computed redox potential differences governing electron transfer between adjacent hemes were adversely affected in that work. Indeed, the unphysically negative potentials caused a severe underestimation of the redox conductivity at high temperature, and *apparent* agreement with the experimental observation of *anti*-Arrhenius kinetics.^38^ This conclusion was reached by Guberman-Pfeffer from a computational analysis^38^ *16-months before* the spectroelectrochemical data was published. Now that the experimental data of Portela *et al*. confirms that analysis, the model of Dahl *et al*. is shown to be physically baseless. There is left in its place no model to rationalize the *anti*-Arrhenius conductivity reported for OmcS, although Guberman-Pfeffer offered several hypotheses that have not yet been investigated.^38^ It is of note, however, that the specific temperature dependence observed by Dahl *et al*. and the senior investigator on that work a decade earlier has not been reproduced by other laboratories.^61^

### 2.4. Redox Cooperativities Dynamically fluctuate Potentials by ≤0.1 V under Electron Flux

As already noted, the computational predictions of Figures 2 and 4 approximated each redox transition as an independent event dominated by a single heme. In reality, however, each macroscopically observed redox transition by spectroelectrochemistry corresponds to a transition between ensembles of microstates in which different hemes are sequentially reduced (Figure 3). What is missing from the prior computations—and even experiments—for OmcS is a quantification of the redox cooperativities, or a measure of how much the redox potential of one heme is shifted by the oxidation of another heme. Portela *et al*. suggested redox cooperativities are comparable to the ≤0.05 V experimental uncertainties, which is consistent with the level of agreement shown in Figure 4. But does an analysis of redox cooperativities using the CryoEM structure come to the same conclusion?

Answering this question by either experimental (*e*.*g*., advanced nuclear magnetic resonance)^57^ or computational techniques requires methodological innovations. The latter were recently reported by Guberman-Pfeffer and validated in a collaboration with the Anderson laboratory.^46^ In that work, redox cooperativities were predicted with quantitative accuracy for *de novo* designed solution-phase and membrane-embedded di-heme maquette proteins using the BioDC program.^46^ BioDC^47^ is an interactive Python workflow that automates the computation of redox potentials, cooperativities, and conductivities in multi-heme cytochromes. As an additional benchmark, BioDC predicts interaction energies of 0.026 (Exp. 0.027±0.002)^62^ eV between Hemes I and III and 0.026 (Exp. 0.041±0.003)^62^ eV between Hemes III and IV (there is no Heme II) in the periplasmic *c*-type cytochrome isoform A (PpcA) from *G. sulfurreducens*. Given that the interaction energies are of the correct magnitude and comparable to thermal energy at 300 K (0.03 eV), the theory-experiment agreement is reasonable.

Interestingly, application of the BioDC methodology to OmcS (Table 2, Figure 5) indicates that *only* the oxidation of the first nearest neighbors along the spiraling staircase of hemes shifts the redox potential of the heme on the adjacent rung by more than thermal energy at 300 K (∼0.1 vs. 0.03 eV). As electrons ‘hop’ from heme-to-heme, the redox potential of the adjacent heme is predicted to shift by ∼0.1 V; in this sense, the redox potentials dynamically fluctuate in response to the flow of electrons. This effect is well-known for multi-heme proteins,^63^ but hitherto not considered for cytochrome ‘nanowires.’

**Table 2.** Heme-heme interaction energy matrix. The diagonal elements (in V vs. SHE) are the redox potentials computed from quantum mechanics/molecular mechanics at molecular dynamics-generated configurations (QM/MM@MD). The green upper-right and blue lower-left off-diagonal elements (in eV) are interaction energies from Poisson-Boltzmann Solvation Area calculations performed using the BioDC workflow on the two cryogenic electron microscopy structures of OmcS, PDB 6EF8 and 6NEF respectively. The prime mark (‘) indicates a heme that belongs to the next subunit in the filament.

| Heme # | 1 | 2 | 3 | 4 | 5 | 6 | 1' |
| --- | --- | --- | --- | --- | --- | --- | --- |
| 1 | **-0.063** | 0.085 | 0.025 | 0.015 | 0.007 | 0.005 | 0.004 |
| 2 | 0.080 | **-0.111** | 0.121 | 0.038 | 0.013 | 0.008 | 0.005 |
| 3 | 0.023 | 0.110 | **-0.071** | 0.095 | 0.025 | 0.015 | 0.007 |
| 4 | 0.013 | 0.034 | 0.087 | **-0.248** | 0.121 | 0.039 | 0.012 |
| 5 | 0.006 | 0.011 | 0.022 | 0.104 | **-0.177** | 0.068 | 0.018 |
| 6 | 0.005 | 0.007 | 0.013 | 0.030 | 0.056 | **-0.095** | 0.102 |
| 1' | 0.004 | 0.005 | 0.007 | 0.011 | 0.015 | 0.090 | **-0.063** |

**Figure 5.**
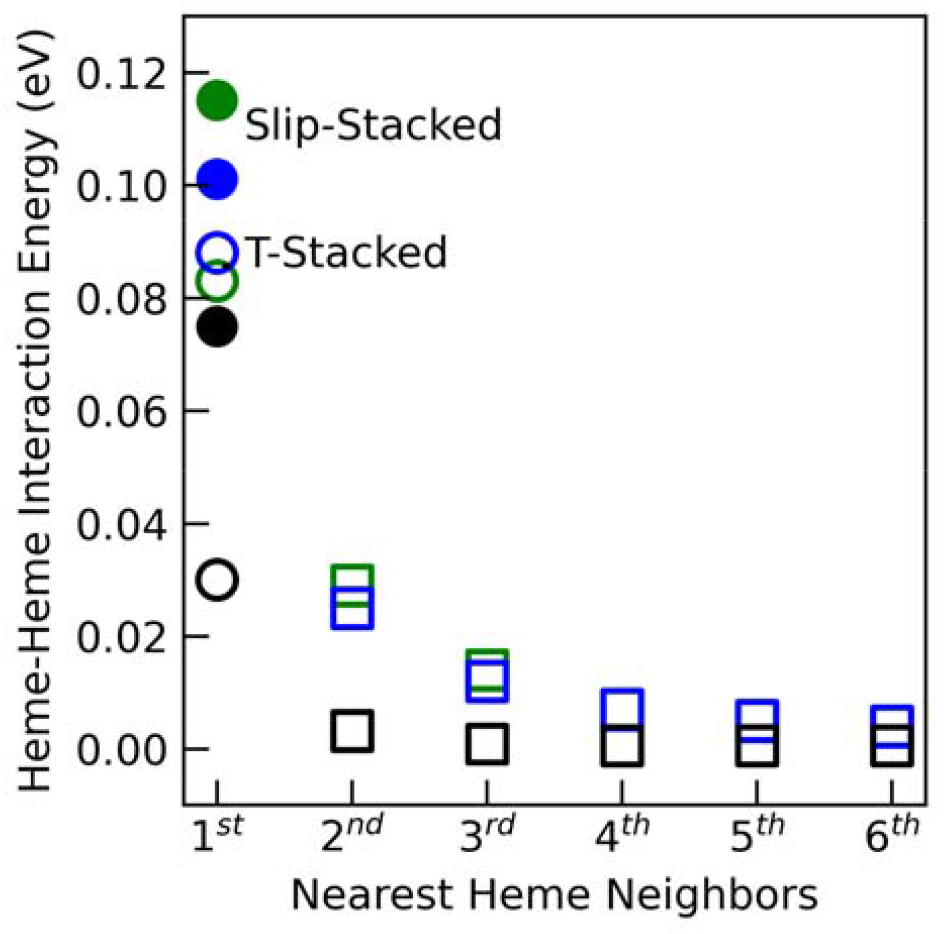
Computed heme-heme electrostatic interaction energies are similar for the two cryogenic electron microscopy structures of OmcS (green = PDB 6EF8; blue = 6NEF) and in close agreement with the predictions of a Debye-Hückel shielded electrostatics (DHSE) model parameterized on experimental data for 17 multi-heme proteins.^64^

Redox cooperativities of a similar (∼0.08 eV) magnitude were previously reported from constant pH and redox molecular dynamics (C(E,pH)MD) simulations^38^ by noting the difference in potentials when each heme was titrated with either all other hemes held in the fully oxidized state or allowed to be titrated at the same time. The interaction energies in Figure 5 are also consistent with, but overestimated compared to the predictions of a Debye-Hückel shielded electrostatics (DHSE) model^64^ for heme groups based on the Fe-to-Fe separation in either the CryoEM structures^30, 31^ or hundreds of nanoseconds of classical molecular dynamics on OmcS (Table S1 in the Supporting Information).

The DHSE model is based on the analysis of redox cooperativities in 17 multi-heme cytochromes from 5 different microbial genera^62, 64^ and assumes an effective dielectric constant (ϵ_eff_) of 8.6. That the heme-heme interaction energy is larger than expected for the Fe-to-Fe separation found in OmcS suggests that the interactions are less effectively screened, or ϵeff is lower than assumed by the DHSE model. Indeed, the same modeling that accurately predicts the spectroelectrochemical potentials ascribed ϵeffs between 3 and 7 (instead of 8.6) to the heme active sites in OmcS. Thus, the analysis of redox cooperativities has provided insight on the ϵeff*s* of the heme binding sites. However, the overestimation of heme-heme electrostatic interactions relative to the DHSE model may also reflect, to some degree, the use of fixed (unpolarized) atomic partial charges in the calculations.

if 1^st^ nearest neighbor heme-heme interaction energies are ∼0.1 eV, why do redox potentials computed assuming no heme-heme interactions come within ≤0.04 eV of the experimentally measured values? There are a few possible explanations: Each heme in the electronically embedded QM/MM redox potential calculations was polarized by the electrostatics of the environment, including adjacent hemes, whereas each heme in the interaction energy calculations was assigned fixed atomic partial charges for the given oxidation state. Because of the presence and absence, respectively, of environment-induced polarization, some heme-heme interaction was already implicitly included in the redox potentials, whereas the interaction energies were overestimated. The latter appears to be the case, for example, relative to the DHSE model.

With the uncertainties on the redox potentials from experiment and computation being ≤0.05 eV and the heme-heme electrostatic interaction energies probably being less than a factor of 2 larger, the foregoing analysis is generally consistent with the conclusion of Portela *et al*. that independent and sequentially-coupled models for the heme redox potentials give comparable fits to the spectroelectrochemical titration curve. There is simply not enough resolution to clearly discern the effect of heme-heme interactions from the error bars on the redox potentials. The implications of this conclusion are that (1) heme-heme interactions can *effectively* be neglected, (2) macroscopic redox potentials are dominated by microstates in which a single heme primarily undergoing the redox transition, and (3) the one-to-one mapping of spectroelectrochemical potentials to the CryoEM structure by their close agreement with structure-based calculations is warranted.

The analysis of Figure 4 should be considered a *prediction* to be validated by advanced nuclear magnetic resonance experiments.^57^ Given that the computed (green) redox titration curve predicted the experimental (black) curve in Figure 2 by ∼1.5 years, the herein described model has already demonstrated predictive ability and its further assessment is therefore eagerly awaited.

### 2.5. Structurally Assigned Spectroelectrochemical Potentials Define a Reversible Free Energy ‘Rollercoaster’ for Electron Transfer

Figure 4 shows the free energy landscape for electron transfer through a unit cell of the OmcS filament based on the mapping in Figure 2 of spectroelectrochemical redox potentials to the CryoEM structure. Transport of an electron through a unit cell of the filament is a reversable process (ΔG° - 0 overall), even though ΔG° for each heme-to-heme electron ‘hop’ can be as exergonic as -0.081 eV (Heme #6 to #1’) or as endergonic as 0.148 eV (#3 to #4). A reversible free energy landscape was previously proposed for the deca-heme metal-reducing cytochrome type F (MtrF) from *Shewanella oneidensis*,^65^ although unlike OmcS, that protein is not polymeric and the electronic couplings were greatest for endergonic instead of exergonic electron transfer steps (Figure S1).

The reversible nature of the free energy landscape in OmcS is a consequence of the filament being homopolymeric and may explain why all known cytochrome ‘nanowires’ so far discovered are in fact homopolymers: The free energy difference for moving an electron through any integer number of unit cells (*e*.*g*., from Heme #1 in one subunit to Heme #1 in any number of subsequent subunits) must be zero. The only free energy differences that matter are those for transferring electrons into and out of the filament.

The pattern of ‘rising’ and ‘falling’ free energy within a subunit that depends on the particular assignment of redox potentials to the geometrical sequence of hemes in the CryoEM structure is irrelevant in this sense and may not be subjected to natural selection. More than three decades ago Page, Moser, and Dutton demonstrated that free energy barriers can be hundreds of millivolts energetically uphill and still not be rate-limiting for electron transfers relative to turnover at catalytic centers, as long as cofactors are within 14 Å; the minimum edge-to-edge spacing between hemes in all known cytochrome filaments is 3 – 6 Å.^8, 66^ The consecutive alternatingly exo- and endergonic |0.10 – 0.35| V potential differences for electron transfer from cytochrome c_2_, through a tetra-heme protein, and into the photo-oxidized special pair of the *Rhodopseudomonas viridis* reaction center is a classic example of how evolution does not seem to optimize free energy landscapes for the *sole* consideration of maximal rate.^66, 67^

Even so, it is still interesting to observe that the free energy of an electron generally increases, reaches a maximum, and then decreases as it moves through a unit cell of the filament. A similar pattern was also computed^39^ for the OmcZ ‘nanowire,’ which is a homopolymer of an octa-heme protein. The redox potentials in OmcZ were proposed to increasingly become more negative towards the middle of a subunit to decrease the energy barrier for ‘leaking’ electrons from the main heme chain onto a nearly fully solvent exposed heme that branches off from it. The ‘leaked’ electrons can more readily reduce soluble species or be transferred to other filaments. But no such physiological purpose can be ascribed to the similarly peaked free energy landscape in OmcS since the hemes are more shielded from the solvent by the protein and there is no branching heme.

It is of utmost importance to note that the free energy landscape in Figure 4 is a *prediction* that corresponds to one of 720 possible ways of assigning the six macroscopic spectroelectrochemical potentials to the six microscopic hemes of the CryoEM structure under the approximation (justified above) of independent heme redox transitions. The particular landscape, which is shown to be physiologically relevant in subsequent sections, is based on the close (<0.04 eV) concurrence between six independently computed redox potentials for the hemes of the solution-phase CryoEM structure and the solution-phase spectroelectrochemically measured potentials. Because Portela *et al*. reported that the redox properties of OmcS were similar in solution and on solid surfaces, the CryoEM structure and the herein described calculations based on it that reproduce those properties are relevant to either experimental condition, as well as the physiological situation of *G. sulfurreducens* growing on a grain of sand.^45^

However unlikely it is for the close theory-experiment agreement on the redox potentials to be due to chance, the following analysis of filament electrical conductivity considers *all possible free energy landscapes*. To this end, the 720 permutations for assigning the six redox potentials to Hemes #1 through #6 were tabulated and ΔG° was computed for every adjacent pair of hemes. Figure 6 shows the resulting distribution of possible ΔG°s for each heme-to-heme electron transfer step; six ΔG° s from the distribution from one of the 719 possible free energy landscapes considered in the below conductivity analysis. Note that the number of landscapes is one less than the number of possible ways of assigning the redox potentials because ΔG° is computed from the difference in potentials between adjacent hemes.

**Figure 6.**
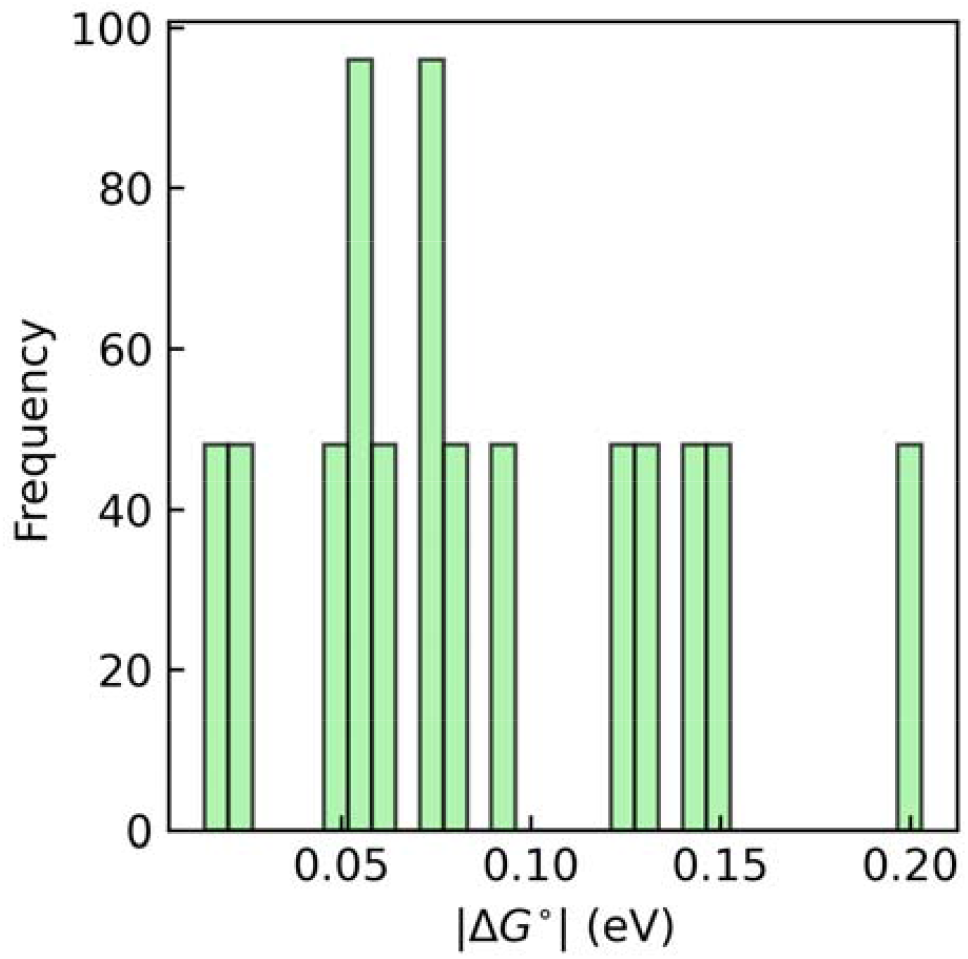
Distribution of all possible s for an electron transfer step in OmcS given the six spectroelectrochemically reported redox potentials. There are 6! or 720 ways of assigning those potentials to the six hemes of an OmcS filament subunit, and therefore 719 possible redox potential differences or s, but the histogram shows a considerable amount of degeneracy.

### 2.6. Electronic polarization substantially lowers Reorganization Energies in OmcS

The reaction free energy just discussed is only a part of the activation barrier (E_a_) for electron transfer in the limit of non-adiabatic Marcus theory (NAMT).^68^ Specifically,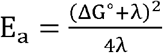, where λ is the reorganization energy. NAMT is the appropriate theoretical framework for the solution-phase physiological redox processes of interest here because heme-heme electronic couplings in monomeric and filamentous multi-heme cytochromes under these conditions are known to be orders of magnitude smaller (0.001 to 0.010 eV versus 0.1 to 1.0 eV) than the reorganization energies.^69^

λ for electron transfers in OmcS has consistently been found to be 0.5 – 1.0 eV from classical molecular dynamics simulations that neglect electronic polarizability,^35, 38, 39^ as well as an approach based on the solvent accessibility of the heme group that was parameterized to reproduce λ from electronically polarizable molecular dynamics.^37^ But the activation-barrier-lowering effects of electronic polarization of the active site and environment on λ have not yet been carefully examined for any cytochrome filament. For context, a brief discussion of the theory is given in Section 2.6.1, followed by the results in Section 2.6.2 to 2.6.3, and the take-away is summarized in Section 2.6.4.

#### 2.6.1. Oxidation-State Dependent Electronic Polarization is a Reorganization Energy-Lowering Mechanism

Electron transfer reorganization energy is generally dissected into inner-(λ_in_) and outer-(λ_out_) sphere components. λ_in_ reflects oxidation-state dependent changes in internal coordinates that, for a heme group, contribute 0.05 – 0.08 eV to the activation barrier.^70, 71^ Small geometrical changes upon reduction/oxidation of the heme group (Figure S2) suggest there is no dramatic change in electronic structure.

λ_out_ reflects the energetic penalty for rearranging the environment to accommodate altered charge distributions on the electron donor and acceptor. λ_out_ is determined from the distributions of electrostatic vertical energy gaps (VEGs, ΔU) in the reactant (R) and product (P) states. If the VEGs between the R and P states are Gaussian distributed and ergodically sampled according to Boltzmann statistics on the timescale of the electron transfer, λ_out_ is equivalently defined in terms of the average ((ΔU)_X_,x = R or P; Eq.1) or variance (σ_X_; Eq.2) of the VEG distributions in the two states. These definitions are respectively known as the Stokes-shift (λ^st^) and variance (λ^var^) reorganization energies; if they are not equivalent, the reaction reorganization energy (λ^rxn^) that should be used in Marcus theory is defined in terms of their ratio (Eq. 3).^51^ In Eqs. 1 to 3, k_b_ and T are respectively the Boltzmann constant and absolute temperature.

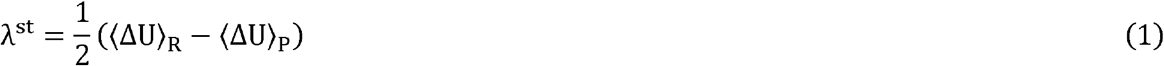

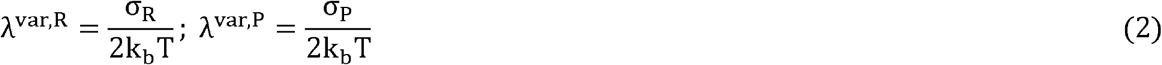

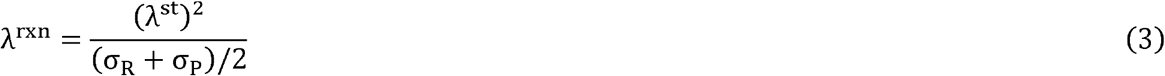

The VEGs of which the statistical moments are taken to compute λ_out_ can be approximated as a sum of terms (Eq. 4):^51^ (1) The intrinsic energy change upon electron transfer (ΔU_0_), which is expected to be nearly zero for self-exchange between different conformers of the same chemical group (here adjacent heme cofactors); (2) The change in Coulombic energy (ΔU_coul_) associated with electrostatic interactions of the environment with the altered charge distributions on the donor (D) and acceptor (A) upon oxidation and reduction, respectively, and (3) the energy associated with the differential polarization (ΔU_pol_) of the donor and the acceptor upon electron transfer under the electric field of the protein-water environment.

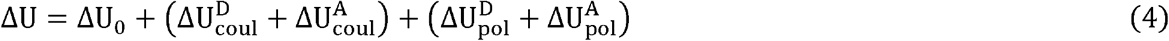

Each ΔU_coul_ is given by the product of the change in partial atomic charge for each atom of the donor or acceptor (Δq) with the electrostatic potential (ϕ) exerted by the environment on that atom, summed over all the atoms of the group (Eq. 5). Each ΔU_pol_ is related to the difference in the second-rank polarizability tensor between the oxidation states for that group (Δα_jk_; j, k =x, y, z) and the electric field vector **(E)** acting on it (Eq. 6), which is approximated according to literature precedent^48, 72^ (although over simplistically) as the field acting on the Fe center of each heme. Note that because Δ always means final – initial, Δα - n(α^red^ − α^ox^), where n = 1 for the acceptor and -1 for the donor.

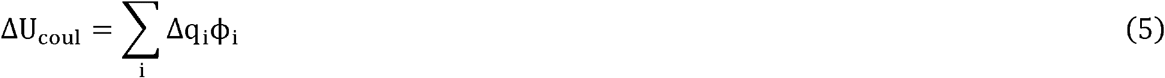

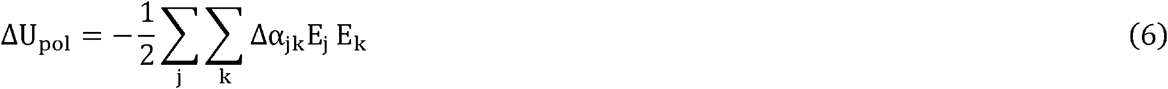

Standard NAMT assumes the ΔU is linearly coupled to the thermal fluctuations of the environment,^51^ which are Gaussian distributed because of the central-limit theorem applied to the large number of particles. In this limit, 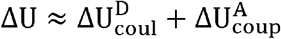 for each (reactant and product) state. Assuming ergodic sampling, all the definitions of λ_out_ (Eqs. 1 to 3) are equivalent in this limit. However, if ΔUpol is non-negligible, then ΔU quadratically depends on the electric field, the energies are no longer Gaussian distributed, λ^var^ > λ^st^, and the effective λrXn that enters into the NAMT rate expression is less than it otherwise would be in the absence of electronic polarization.^49^

Given that the maximum ΔGO possible with the spectroelectrochemical redox potentials (Figure 6) is at least 2.5-fold smaller than λ_out_ computed without electronic polarization,^39^ it is clear that the λ-dominated activation energies can be significantly lowered if ΔU_pol_ is sizable. It is important to know whether this effect is needed to account for the cellular flux of ∼10^6^ electrons/s through >20^44^ to ∼100^45^ filaments/cell.

#### 2.6.2. Electronic Polarization of the Environment Lowers Reorganization Energy by ∼0.15 eV in OmcS

λ_out_ is routinely computed from distributions of ΔU obtained during classical molecular dynamics simulations that lack both electronic polarization of the active site and the environment. To account for electronic polarization of the environment in response to oxidation-state cycling at the active site, a scaling factor (designated *f*_*λ*_) of 0.56 – 0.80 has been recommended.^73, 74^ Application of this scaling factor to the set of λ^rxn^ in Table 3 that entirely neglects polarization results in decreases of as much as 0.14 when *f*_*λ*_ = 0.80 and 0.17 eV when *f*_*λ*_ - 0.56. The ranges for λ^rxn^ respectively become 0.344 – 0.562 eV and 0.241 – 0.393 eV. Both scaled sets of λ^rxn^ are considered in the conductivity analysis below to reach conclusions that are independent of the particular choice of *f*_*λ*_. Dynamical simulations performed with polarizable forcefields are needed—but remain computationally intractable for the necessary timescales—to refine more precisely how much electronic polarizability of the environment reduces λ_out_ for OmcS.

**Table 3.** Electronic polarization of each heme in a unit cell of the OmcS filament significantly lowers the reorganization energy, though the effect is probably overestimated by the exaggerated^75^ magnitude and variance of the electric field generated by a fixed-point charge classical forcefield for molecular dynamics.

| | $\lambda^{\text{st}}$ (Eq. 1) | $\lambda^{\text{var,f}}$ (Eq. 2) | $\lambda^{\text{var,b}}$ (Eq. 2) | $\lambda^{\text{rxn}}$ (Eq. 3) | $\kappa_g^{\text{a}}$ |
| --- | --- | --- | --- | --- | --- |
| Unpolarized Active Site |  |  |  |  |  |
| 1↔2 | 0.813 | 0.910 | 0.972 | 0.702 | 1.158 |
| 2↔3 | 0.470 | 0.511 | 0.516 | 0.430 | 1.092 |
| 3↔4 | 0.566 | 0.561 | 0.524 | 0.590 | 0.959 |
| 4↔5 | 0.662 | 0.649 | 0.606 | 0.697 | 0.949 |
| 5↔6 | 0.646 | 0.775 | 0.738 | 0.551 | 1.172 |
| 6↔1' | 0.570 | 0.689 | 0.636 | 0.490 | 1.163 |
| Polarized Active Site |  |  |  |  |  |
| 1↔2 | 0.664 | 3.808 | 3.212 | 0.125 | 5.289 |
| 2↔3 | 0.434 | 1.693 | 1.174 | 0.131 | 3.302 |
| 3↔4 | 0.946 | 2.126 | 3.536 | 0.316 | 2.994 |
| 4↔5 | 0.595 | 0.845 | 2.960 | 0.060 | 9.924 |
| 5↔6 | 0.299 | 2.645 | 3.425 | 0.029 | 10.162 |
| 6↔1' | 0.593 | 2.519 | 4.609 | 0.099 | 6.012 |

<sup>a</sup> $\kappa_g = \frac{\lambda^{\text{var}}}{\lambda^{\text{st}}}$ . Deviations from unity quantify departure from the assumptions of standard non-adiabatic Marcus theory.

#### 2.6.3. Electronic Polarization of Heme by a Sizable Electric Field Lowers Reorganization Energy by 0.2 – 0.6 eV in OmcS

Active site polarizability has been considered (here for OmcS and earlier for other proteins^48-52^) by evaluating Eq. 6 for the donor and acceptor in both the reactant and product states using Δα obtained from quantum mechanical calculations and the electric field exerted by the fixed-charge (non-polarizable) environment of classical molecular dynamics simulations. Using this approach, it was proposed^48, 49, 51^ that the change in electronic polarizability upon reduction of the heme cofactor significantly lowers λ_out_ in cytochrome *c*. The claim was challenged and the observation was instead attributed to a 0.1 V/Å difference in the sampled protein-water electric field.^50, 52^ It is conceivable that either mechanism also applies to the differently (His-His instead of His-Met) ligated hemes of polymeric multi-heme filaments like OmcS.

To test this possibility, Δα was examined for His-Met and His-His ligated hemes, both in vacuum and in the protein-water environments of cytochrome *c* and OmcS, respectively. As described in detail in the *Assessment of Heme Electronic Polarizability* section of the Supporting Information, the ligand set (His-His versus His-Met) and the presence or absence of the protein-water environment did not have a large effect on the computed polarizability tensor. The discussion here therefore focuses on the results for the His-His ligated heme of OmcS in vacuum and makes comparisons with the literature on the His-Met ligated heme of cytochrome *c* in vacuum.

Density functional theory (DFT) techniques have been found to accurately reproduce the (iso/aniso)-tropic polarizabilities of rigid to semi-rigid molecules,^76-78^ including a heme memetic.^79^ Using these methods and ensuring basis set convergence on the results, Figure 7 (Table S6) shows that the isotropic polarizability (α_iso_) is 86 to 104 Å^3^ in either oxidation state regardless of the approximate density functional and basis set combination examined. This range falls within the experimental uncertainty on α_iso_ for the heme memetic *meso*-tetraphenyl-iron(III) porphyrin,^79^ and exceeds the prior estimates of 54 and 27 Å^3^ in the reduced and oxidized states, respectively, from the Zerner’s Intermediate Neglect of Differential Overlap with Singles (ZINDO/S)^80, 81^ method applied to a His-Met ligated *c*-type heme. The change in isotropic polarizability between oxidation states (Δα) is found by DFT methods to be at most 6.5 Å^3^, which is 4-fold smaller than the previous prediction by ZINDO/s, but in good agreement with the lower-estimate obtained from an analysis of the UV-visible absorption spectrum of cytochrome *c*.^82^

**Figure 7.**
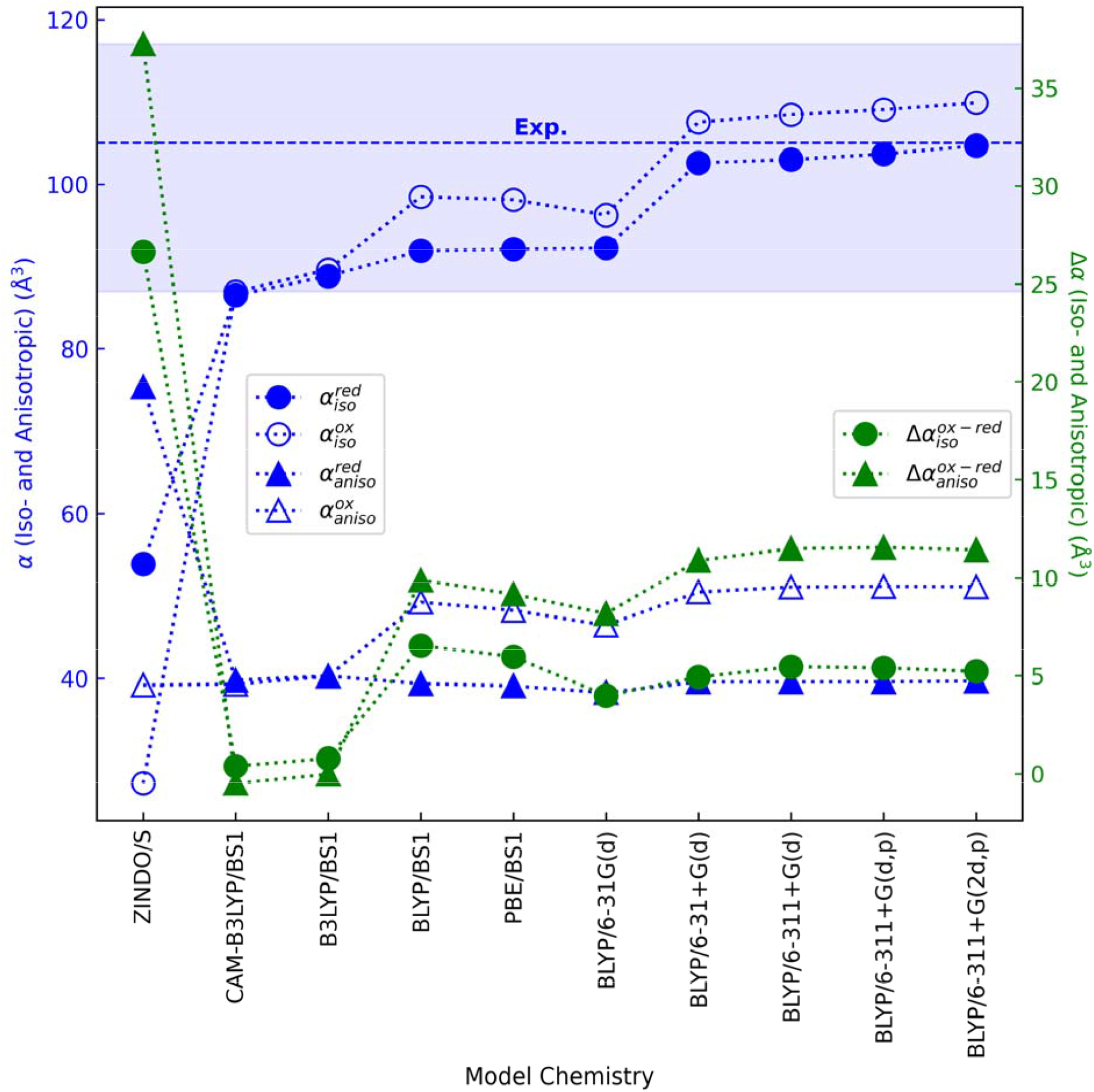
Predicted iso- and anisotropic polarizabilities for a His-His ligated *c*-type heme in the reduced and oxidized states, as well as the difference polarizabilities between oxidation states are similar regardless of the approximate density functional and basis set combination tested, especially in contrast to results previously published with ZINDO/s.^51, 52^ The average (dashed line) and standard deviation (shaded region) of the experimental isotropic polarizability for the heme memetic tetraphenylporphyrin-iron(III) chloride are shown.^79^ Neither the isotropic polarizability in the reduced state nor the anisotropic polarizability in either oxidation state were reported.

A similar story applies to the anisotropic polarizability (α_aniso_). DFT methods predict α_aniso_ to be ∼40 and ∼50 Å^3^ in the reduced and oxidized states, respectively, giving Δα_anisw_ of ∼10 Å^3^. Prior ZINDO/s calculations on a His-Met ligated heme, by contrast, predicted 75 and 39 Å^3^ in the reduced and oxidized states, respectively, with a difference, again, four-fold larger than found by DFT methods.

The discrepancy between ZINDO/s and DFT most likely reflects the inadequacy and unreliability of the former method for predicting polarizabilities. The use of ZINDO/s for this purpose requires summing contributions to the polarizability from vertical excitations computed up to 10 eV of the ground state.^80, 81^ As the sum over excited states must grow to large (*e*.*g*., 100)^48^ numbers to converge the polarizability, increasingly larger errors accumulate when using a method parameterized to reproduce only the lowest-lying excited states. Furthermore, ZINDO/s, unlike DFT, has not been parameterized or benchmarked against experiment for computing polarizabilities.

The overall conclusion is that: Contrary to prior work at the ZINDO/s level of theory, oxidation-state dependent changes in iso- and anisotropic polarizabilities for the heme cofactor are <15 Å^3^ regardless of the different axial ligands and molecular environments. This magnitude, which is visualized in Figure 8 for the type of heme found in OmcS, is considered much too small^82^ to substantially change λ_out_. However, as shown next, sizable electric field magnitudes and variances modify this conclusion because λ^var^ is proportional to the squared fluctuations in ΔU_pol_ (Eq. 2), which in turn depends on the squared field magnitude (Eq. 6).^52^

**Figure 8.**
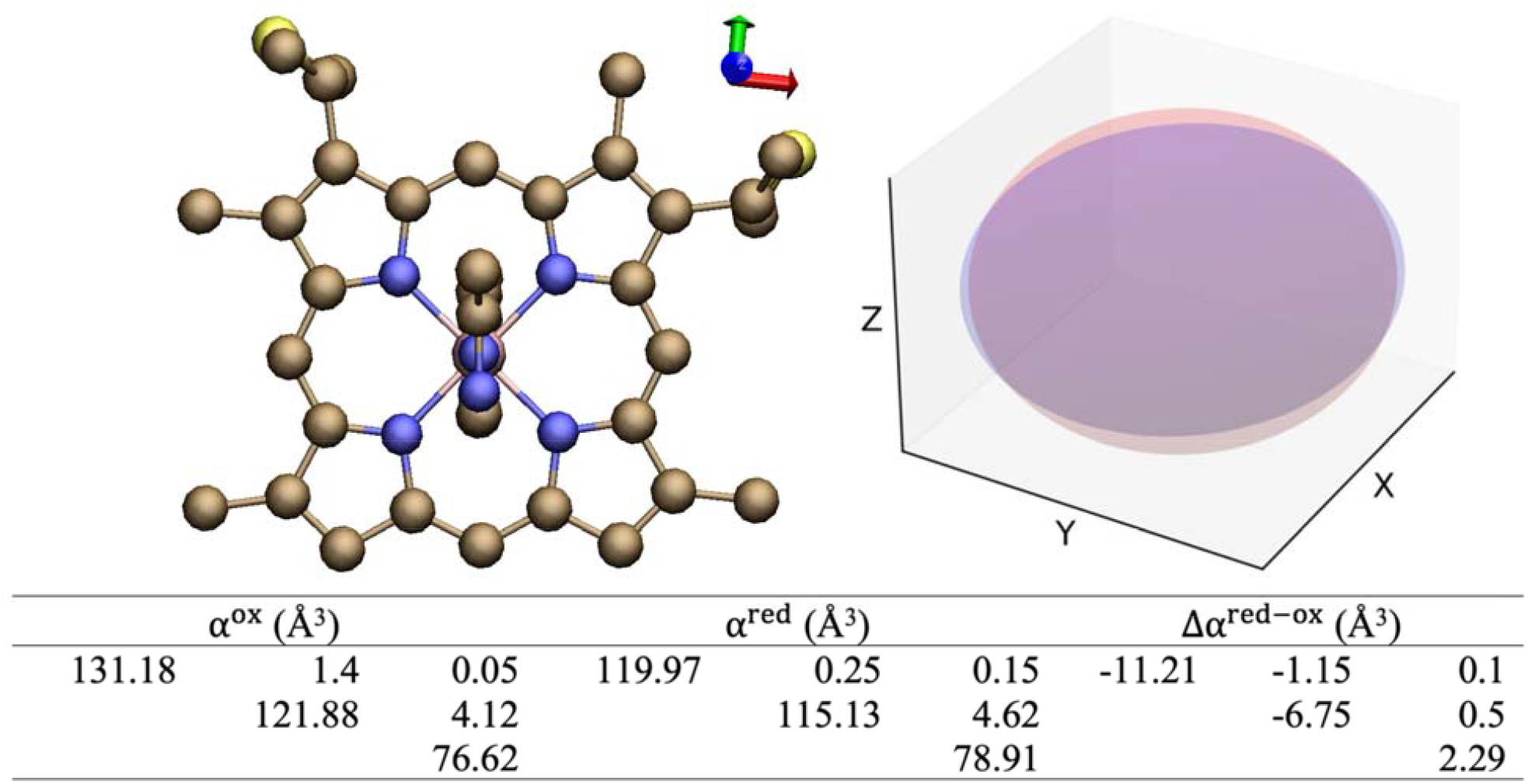
The dipolar polarizability tensor for the heme cofactor minimally changes upon reduction/oxidation. The symmetrical tensor in the oxidized and reduced states is shown in upper-right triangle form and visualized as blue and red ellipsoids, respectively. The difference tensor is also given. The xx, yy, anc zz components of the tensors have the largest magnitudes and are directed along the red, green, and blue directions indicated next to the heme structure. The tensors were computed at the BLYP/6-311+G(2d,p) level of theory for the shown structure (hydrogens omitted for clarity), which was optimized using the BLYP/6-31G(d) model chemistry.

Figure 9a shows the distribution of electric field magnitudes exerted on the Fe centers of the reduced and oxidized hemes in the reactant and product states for the Heme #1 → #2 electron transfer. Similar histograms for the other electron transfer steps in the filament, as well as tabulated statistical descriptors are presented in Table S10 and Figures S4 to S9 of the Supporting Information. The *Influence of Electronic polarization on Reorganization energy* section of the Supporting Information describes in depth the computational protocol for the analysis in Figure 9b.

**Figure 9.**
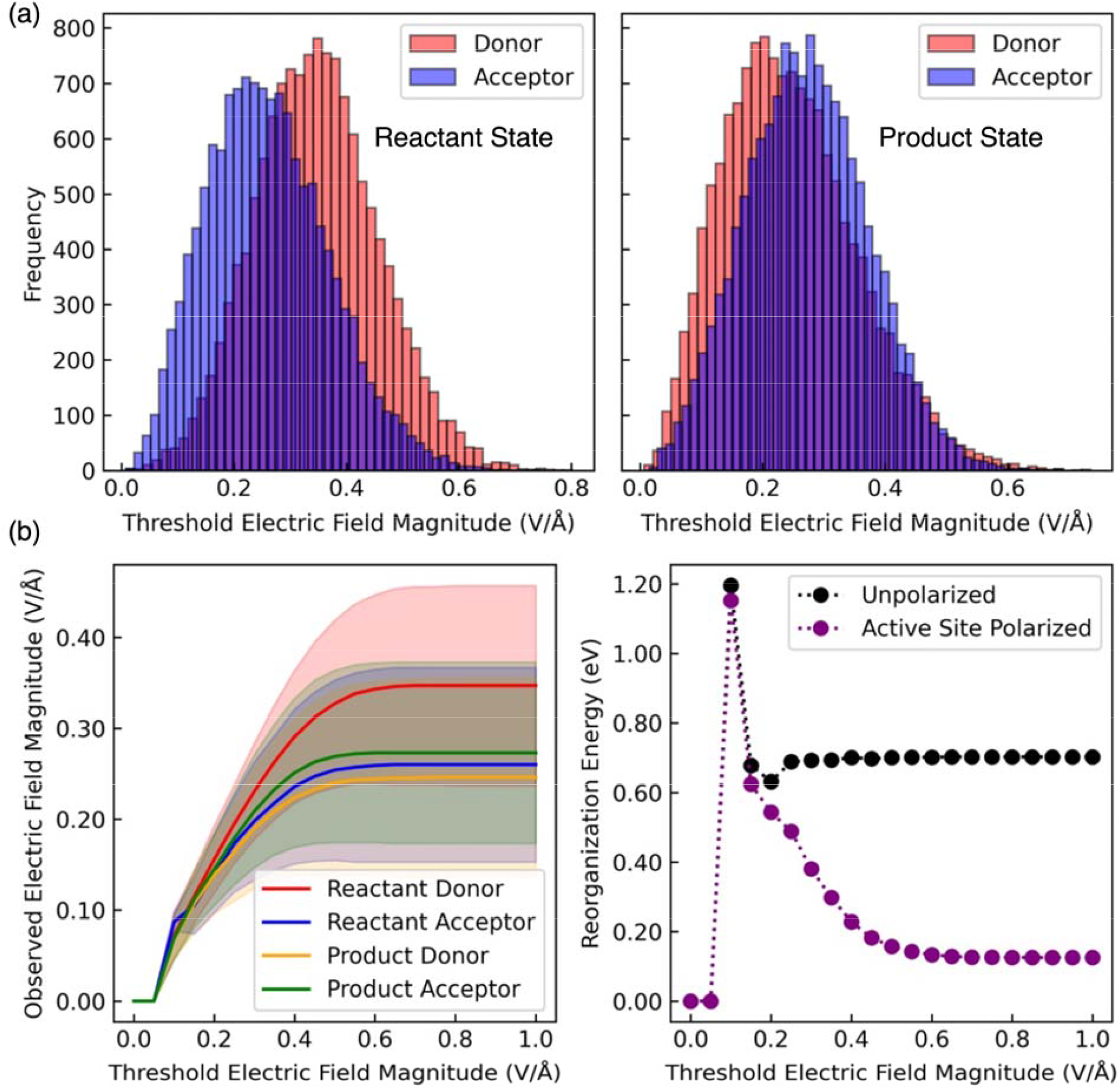
A wide distribution of electric field magnitudes exerted on the Fe centers of the heme groups in OmcS induces a significant decrease in the outer-sphere reorganization energy. In (a), the distribution of field magnitudes on the reduced donor (red) and oxidized acceptor (blue) in the reactant (left) and product (right) states are shown. In part (b), the average magnitude and standard deviation of the field on the donor and acceptor in both states (left) and the reorganization energy (right) are plotted as a function of the threshold field magnitude used to filter the configurations from molecular dynamics. The data is shown for electron transfer between Hemes #1 and #2. Analogous data for all heme pairs in OmcS is presented in Figure S23 to S27 of the Supporting Information.

Figure 9b shows how the reorganization energy depends on the magnitude of the electric field and its variance by filtering configurations from molecular dynamics in which the field magnitude is within a given threshold value.

As shown in Figure 9a for one heme pair, the distributions of electric field magnitudes in OmcS are generally centered at 0.2 to 0.5 V/Å and have standard deviations of ∼0.1 V/Å. Table S10 shows that the standard deviation in the magnitude of each Cartesian-coordinate component of the electric field vector is also ∼0.1 V/Å. These distributions couple to Δα to significantly lower the λ^rxn^ by 0.234 to 0.637 eV (Table 3). However, these results are most likely overestimates. The electric field magnitude and variance in classical molecular dynamics simulations have been shown to be overestimated by the non-polarizable forcefields used here and in other studies.^75^ Electrical conductivities are predicted in a subsequent section neglecting and including these estimates of λ-lowering due to active site polarizability.

#### 2.6.4. Summary of the Effect of Electronic Polarization on Reorganization Energy

The take-away from Section 2.6.1 to 2.6.3 is accounting for electronic polarization of the environment or the active site lowers λ^rxn^ by hundreds of meV. To have conclusions insensitive to the various approximations, four sets of reorganization energies are considered in the conductivity analysis (Figure 10): (1) The unpolarized set directly computed from ΔU = ∑_D,A_ ΔU_coul_ obtained from non-polarizable classical molecular dynamics; (2) The active-site polarized set in which ΔU = ∑_D,A_ ΔU_coul_ +L ∑_D,A_ ΔU_pol_; (3) The environment-polarized set with *f* _*λ*_ = 0.8; and (4) The environment-polarized set with *f* _*λ*_ = 0.56. Prior work has argued that either the correction for active site or environment polarization taken separately is sufficient to bring computed reorganization energies into accord with experiment, and therefore the combination of both of them is not considered here.^50, 51^

**Figure 10.**
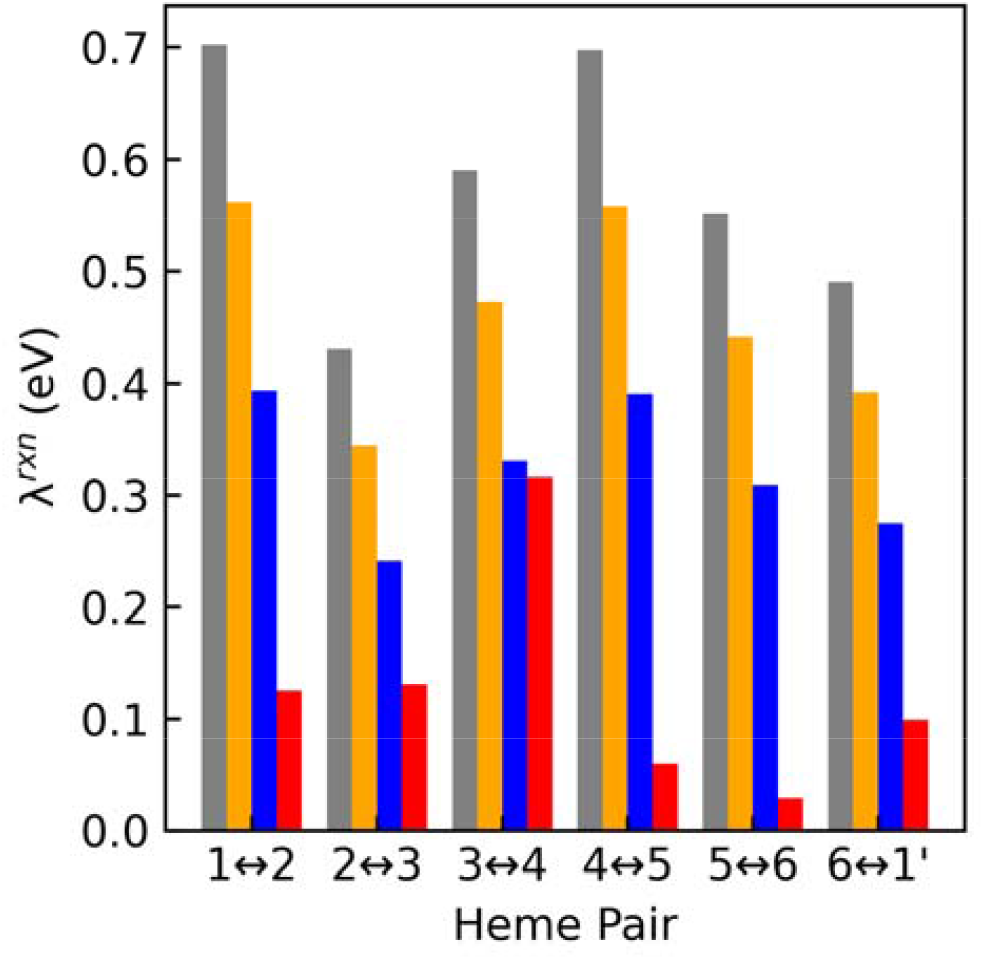
Four different sets of reorganization energies for electron transfer through a unit cell of the OmcS filament are considered for the conductivity analysis that differ by the inclusion of electronic polarization. The different sets reflect no electronic polarization (gray), polarization of the environment computed by scaling the unpolarized values by either 0.80 (orange) or 0.56 (blue), or polarization of the active site computed via Eq. 7 (red). Prior work has argued for including the electronic polarization of the active site or environment, but not both together.^50, 51^

### 2.7. Heme-to-Heme Electron Transfer Energetics are Conserved Among Cytochrome ‘Nanowires’

Figure 11 shows the succession of activation energies for electron transfers through a unit cell of the OmcS filament in the “forward” (N → C-terminal) and “Backward” (C → N-terminal) directions assuming the computationally-assigned ΔG° landscape of Figure 4 and the four sets of λ^rxn^ accounting for different amounts of electronic polarization. Figure 12 shows the distribution of activation energies for each electron transfer step obtained by pairing 719 possible ΔG° landscapes for different ways of assigning the spectroelectrochemical potentials to the hemes with the four sets of λ^rxn^. Note that selection of a particular activation energy for one electron transfer step contracts the distribution of possible values for the remaining steps because the selected activation energy removes possible combinations of the six redox potentials for the other hemes.

**Figure 11.**
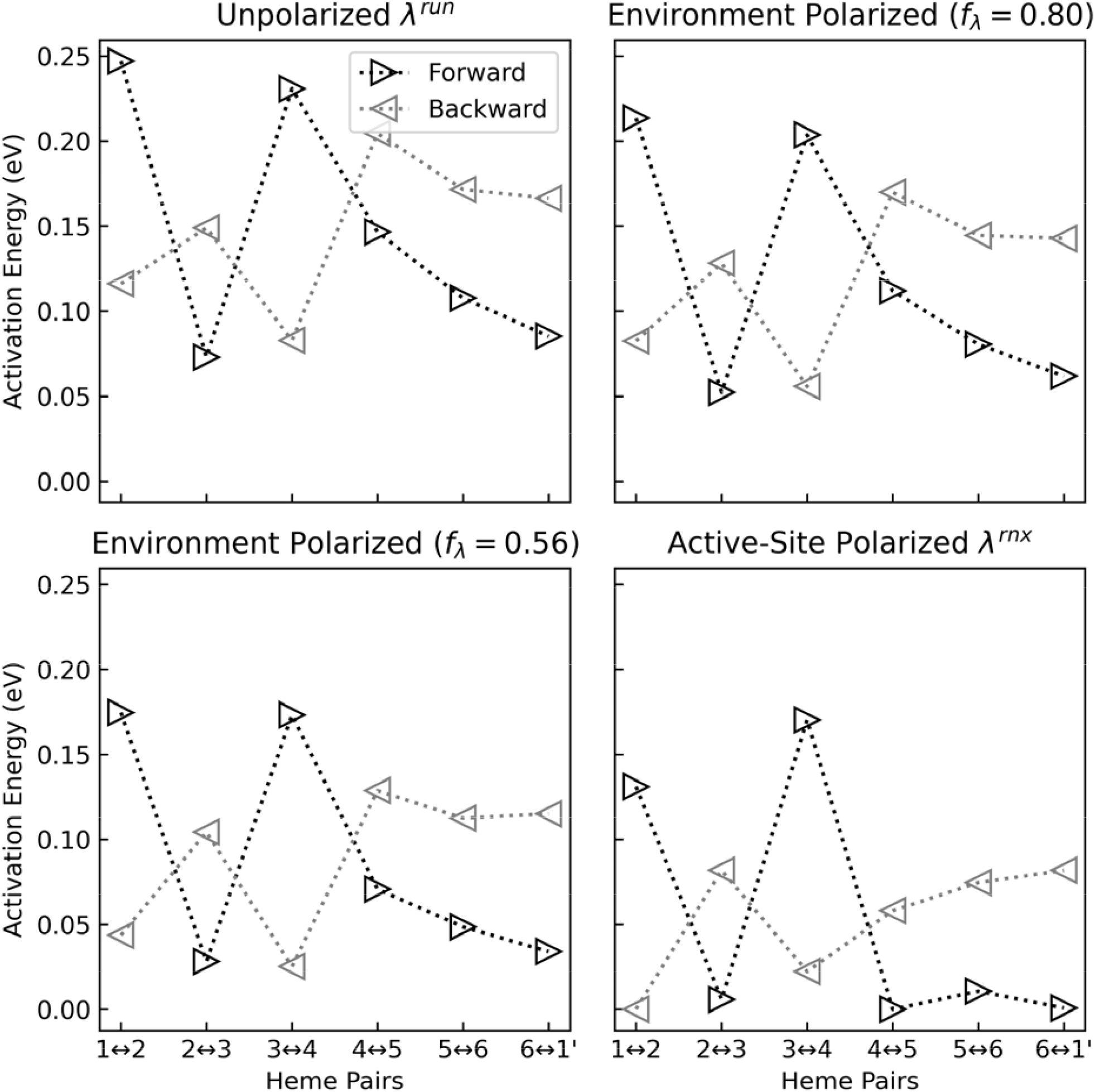
The activation energies for successive electron transfers through a unit cell of the OmcS filament assuming the computationally assigned free energy landscape of Figure 4 and four different sets of reaction reorganization energy that differ in the amount of electronic polarization that is included.

**Figure 12.**
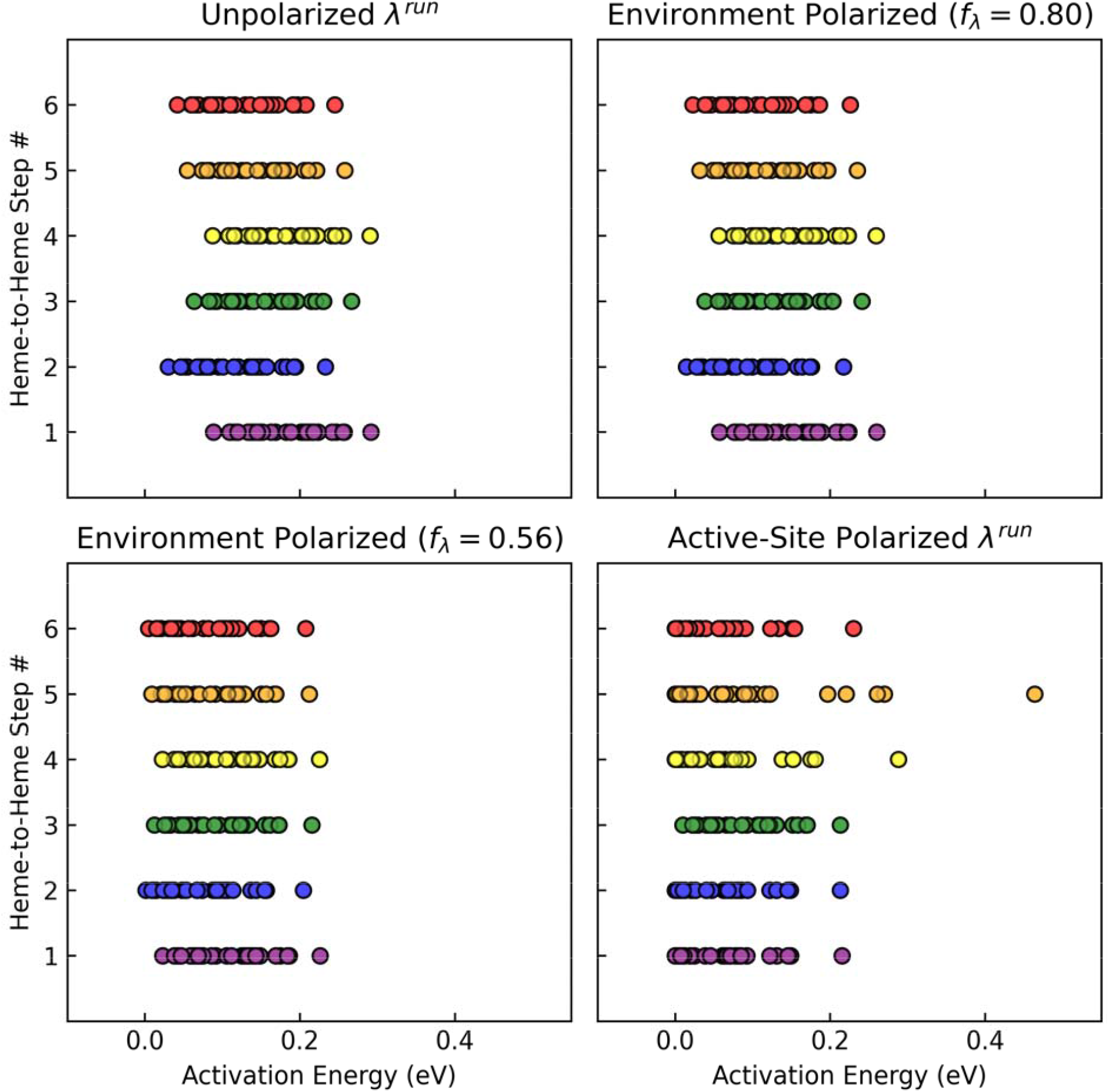
Distributions of activation energies for each electron transfer step through a unit cell of the OmcS filament for all 720 possible ways of assigning the previously reported spectroelectrochemical potentials to the hemes in the CryoEM structure of OmcS.

A key observation is that E_a_s are generally ≤0.28 eV regardless of whether and how electronic polarization is considered in λ^rxn^ (Figure 11) and regardless of which of the 719 possible ΔG ° landscapes are selected (Figure 12). The maximum E_a_ in either direction through the filament is typically 0.1 – 0.2 eV. Similar E_a_s are predicted for all structurally characterized cytochrome ‘nanowires’ to date (Table S12 in the Supporting Information). This finding can be understood given the considerations in the following two paragraphs.

The minimum edge-to-edge van der Waals distance between adjacent hemes in the highly conserved packing motifs ensures both hemes experience a similar electrostatic environment. Because the electrostatic environment seems to overwhelmingly tune heme redox potentials in these proteins (as opposed to conformational distortions of the semi-rigid cofactor),^39^ adjacent hemes that are chemically identical (same peripheral substituents and axial ligands) will have similar redox potentials, or a small redox potential difference (*i*.*e*., a small ΔG° for electron transfer). For several multi-heme cytochromes studied to date, ΔG° is ≤|0.3| eV.^37, 39, 47, 63, 83^

The λ^rxn^ component of E_a_ is also likely to be similar for the structural class. The λ_in_ contribution will be similar because the same redox active heme cofactor is involved. The λ_out_ contribution should be similar because the reorganization of the protein-water matrix is a collective property of the environment with as much as 50% coming from the solvent.^70^ Binding sites with similar dielectric properties may be expected to bind the same cofactor, and thus, to have similar reorganization energies.

Returning to Figure 11, it is notable that the sum of activation energies through a unit cell of the OmcS filament in is 0.3 to 0.9 eV. The range is the same regardless of which of the 719 possible ΔG° landscapes is computed with any of the four sets of λ^rxn^ that account for different amounts of electronic polarization (Figure 13). The accumulated energy barrier for moving an electron through a 300-nm ‘short’ OmcS filament composed of ∼64 subunits would be 19 to 58 eV. This energetic penalty is far larger than the energy provided by the 0.070 and 0.170 V differences between the intra- and extra-cellular redox reactions for the minimal and maximal cellular respiratory rate of *Geobacter sulfurreducens*.^54^ The conclusion follows that the electrochemical gradient is *insufficient* to drive electrons through a micron-long cytochrome ‘nanowire.’ If the filaments are physiologically relevant, the driving force must instead be supplied by the concentration gradient of ∼10^6^ electrons/s that need to be discharged by a single cell.

**Figure 13.**
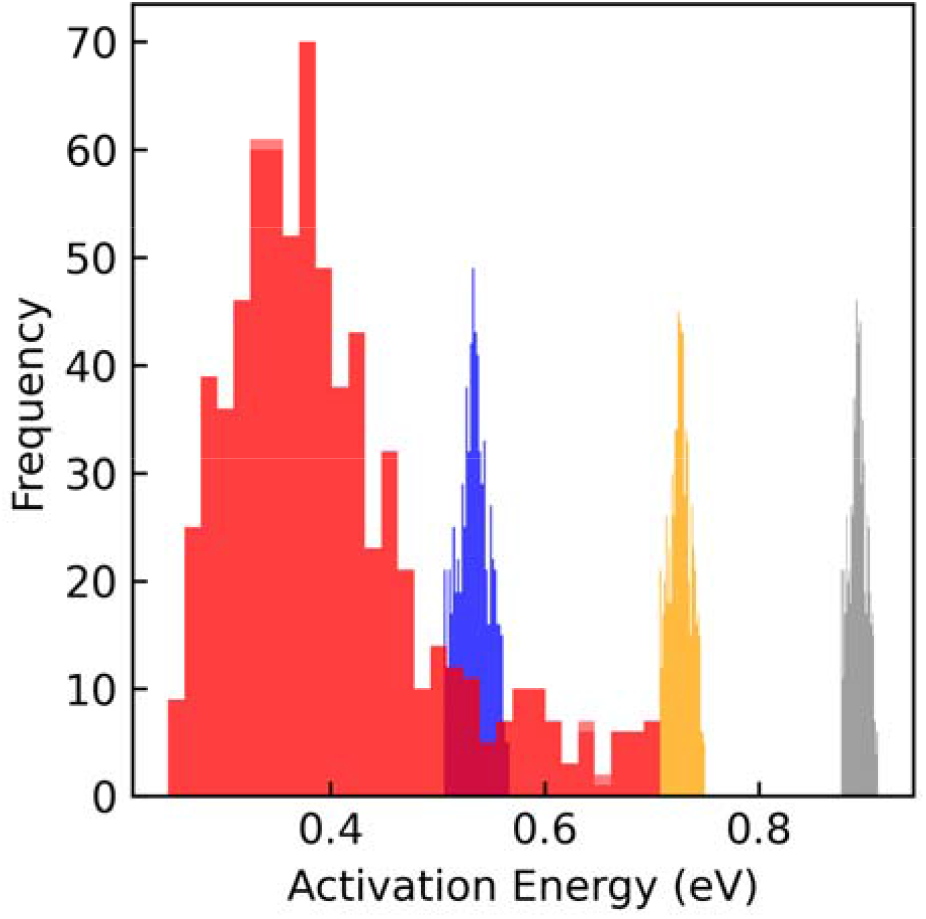
The distribution of accumulated activation energy through a unit cell of the OmcS filament falls in the 0.3 to 0.9 eV range for all 719 possible landscapes associated with different assignments of the spectroelectrochemical redox potentials to the hemes of the CryoEM structure. The gray, orange, blue, and red histograms correspond to pairing these landscapes with reaction reorganization energies that entirely neglect electronic polarization, include polarization of the environment by either scaling the unpolarized values by factors of 0.80 or 0.56, respectively, or include polarization of the active site via Eq. 6. The latter distribution is much more sensitive to the details of the free energy landscape because the reorganization energy is comparable in magnitude.

As with E_a_, computations on OmcS^35, 37-39^ and other multi-heme proteins with the same heme packing motifs^63, 65^ establish that the root-mean-squared electronic coupling 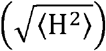 between hemes is ≤ 0.016 eV. This result is a consequence of 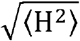 principally being determined by the highly conserved^8^ distance and orientation of the charge donating and accepting heme groups.^84,85^

Thus, the two energetic quantities controlling the charge transfer rate (k_et_) in the framework of non-adiabatic Marcus theory,

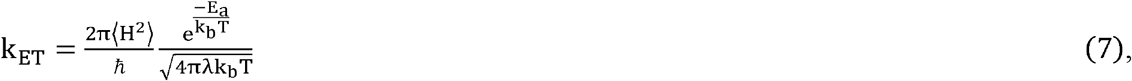

are highly conserved for the structural class of polymeric multi-heme cytochrome filaments. In Eq 7, *ħ*, k_b_, and T are respectively the Plank constant, Boltzmann constant, and absolute temperature (300 K). Conserved energetic ranges imply, by Eq. 7, conserved kinetic ranges, which means that the overall conductivity of a cytochrome ‘nanowire’ should fall in a limited range. The present work therefore has general implications beyond the specific OmcS filament.

### 2.8. Heme Chains Encode few ps to hundreds of ns Electron Transfer Rates

Using the energetic quantities specific to OmcS (Figure 4 for ΔG°, Table 3 for λ^rxn^, and a previously report^38^ for 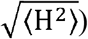, Eq. 7 returns (Figure 14) electron transfer time constants 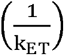 of 18 ps – 556 ns (unpolarized λ^rxn^), 7 ps – 135 ns environment polarized λ^rxn^, *f* _λ_ = 0.8), 2 ps – 25 ns (environment polarized λ^rxn^, *f* _λ_ = 0.56), and 0.2 ps – 4 ns (active site polarized λ^rxn^). Slower rates of 3 to 591 μs are realized when all 719 possible ΔG° landscapes are considered (Figure 15). The implication is that any assignment of the spectroelectrochemical redox potentials to select a particular ΔG° landscape, paired with a set of λ^rxn^s that does or does not include the effect of electronic polarization gives rates that meet or exceed the typical millisecond timescale for enzymatic turnover.^86^

**Figure 14.**
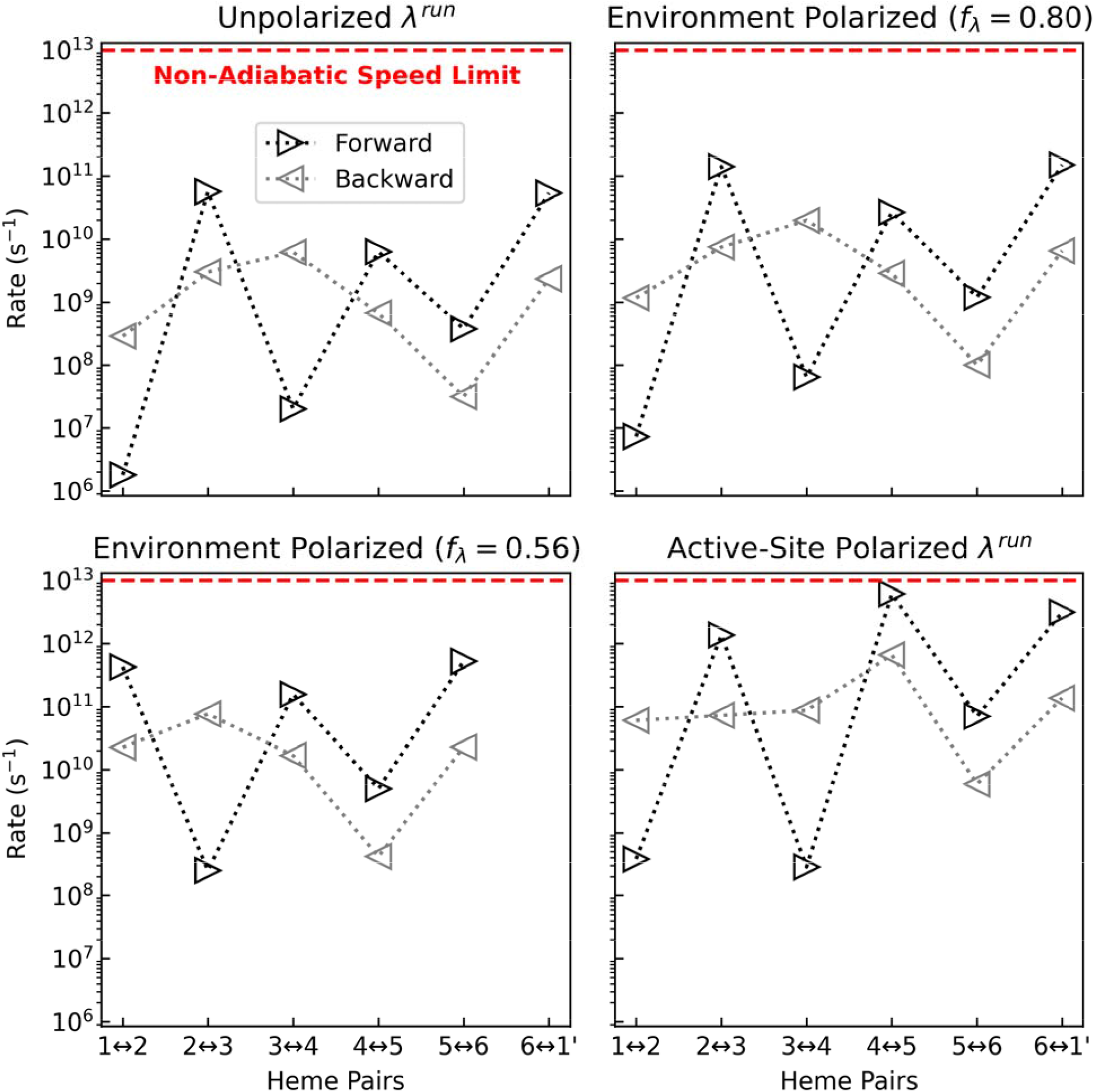
Rates of 10^6^ to 10^13^ s^-1^ (hundreds of ns to sub-ps) are predicted based on the computationally assigned landscape, the four sets of accounting for different amounts of electronic polarization between oxidation states, and previously computed of a generally accepted magnitude for adjacent hemes in multi-heme cytochromes. The rates of ≤1 ps are physically unrealistic, as explained in the main text.

**Figure 15.**
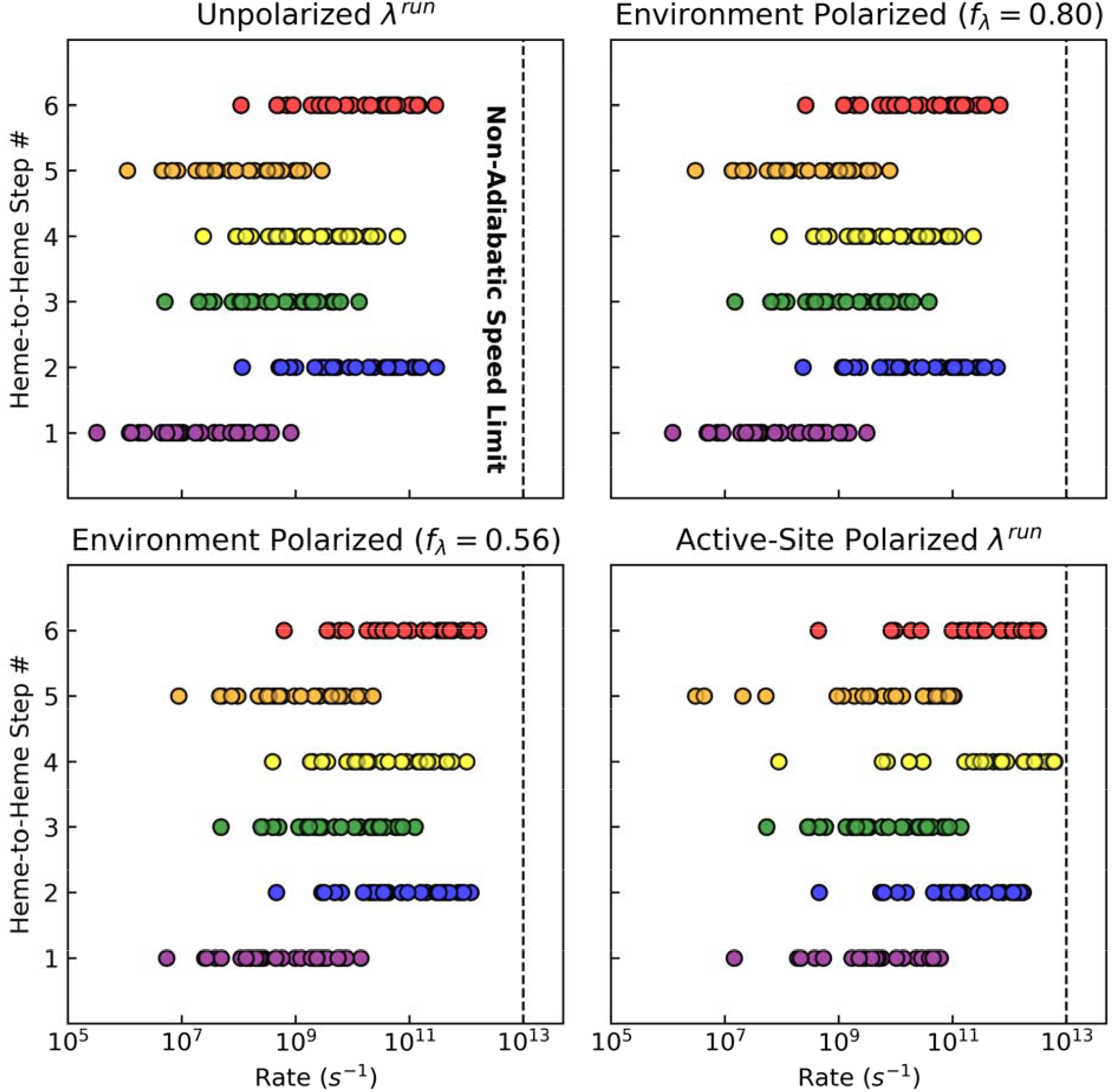
Distributions of reaction rates for each electron transfer step through a unit cell of the OmcS filament for all 720 possible ways of assigning the previously reported spectroelectrochemical potentials to the hemes in the structure paired with four sets of that differ in the account of electronic polarization.

For context on the high-end of the predicted rates, the time constant associated with the electronic coupling-maximized non-adiabatic speed limit for electron transfer between metal ions in van der Waals contact is ∼0.1 ps.^87^ Since the Fe centers of adjacent hemes in any cytochrome filament are separated by a distance at least three-fold larger (9 – 12 vs. 3 Å), the physical plausibility of sub-ps rates should therefore be questioned. In the active-site polarized λ^rxn^ case (Figures 14 and 15), the exaggerated electric field magnitude and variance from non-polarizable classical simulations likely produced an over-polarization of the hemes, which in turn lowered the reorganization energies and activation barriers too much.

Sub-picosecond electron transfer rates in OmcS were proposed to explain transient absorption spectral features,^88^ but these rates pertain to photo-excited charge transfer, *not* the ground-state, physiologically relevant electron transfer process of interest here for the non-photosynthetic bacterium *Geobacter sulfurreducens*. The photo-excited electron transfer was proposed to be a disproportionation reaction in which an oxidized heme transfers an electron to an adjacent oxidized heme to give a radical cation ‘doubly’ oxidized heme and a reduced heme. That process, if it occurs, is very different from the physiologically-relevant reaction of a reduced heme transferring an electron to an adjacent oxidized heme to give an oxidized heme and a reduced heme, respectively. Furthermore, the quantum dynamical calculations performed (by and to the regret of the present author) to rationalize sub-ps rates in that study inappropriately neglected nuclear motions, electron-electron interactions, and the presence of the protein environment, on top of using an inadequate description (semi-empirical Extended Hückel theory) for the molecular orbitals.^40^

More insight is available by categorizing the predicted rates as between parallel- or perpendicularly-stacked adjacent hemes, and by comparing to experimentally measured heme-to-heme electron transfer rates in other multi-heme cytochromes with these highly conserved packing geometries (Figure 16).^89, 90^ Two key observations are apparent from the figure: (1) Rates for perpendicularly-stacked hemes are slower and therefore rate limiting compared to rates for parallel-stacked hemes; and (2) Predicted rates meet or exceed experimental expectations for heme-to-heme electron transfers. The prediction of rates in excess of the experimental estimates departs from an earlier finding that neither considered the reported spectroelectrochemical potentials to define the ΔG° landscape nor electronic polarization effects on λ^rxn^.^27^

**Figure 16.**
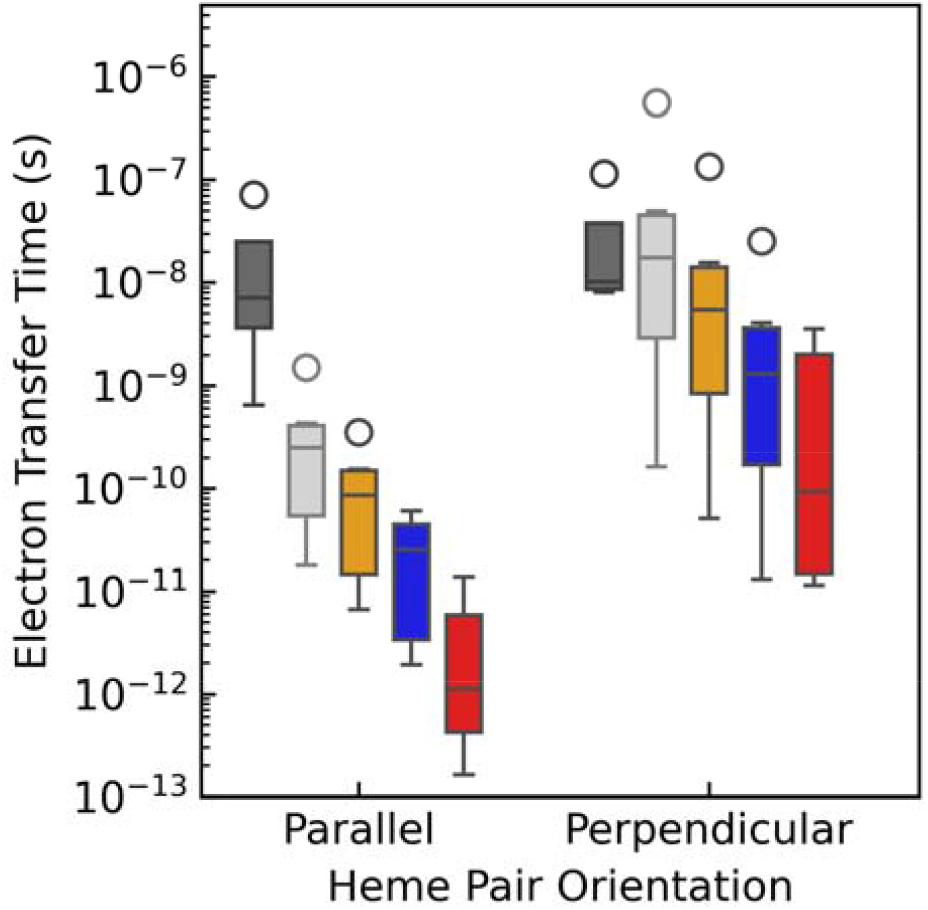
Predicted heme-to-heme electron transfer rates meet or exceed experimental expectations from other multi-heme cytochromes with the same parallel and perpendicular heme-stacking motifs when assuming the computationally assigned landscape (Figure 4) and any of the four sets of s that differ in the inclusion of electronic polarization. The color scheme is the same as in Figure 10, except dark gray indicates the previously reported experimental rates^89, 90^ whereas light gray corresponds to the rates obtained using s that entirely neglected electronic polarization.

The first of these observations deserves additional comment. Because all known cytochrome filaments feature an almost equal proportion of perpendicular to parallel stacked hemes,^8, 30-33^ it is clear that the heme chains in these proteins are not optimized for maximal electrical conductivity. The typical arrangement of alternating parallel and perpendicularly-stacked hemes may reflect a geometrical constraint imposed by the thioether linkages of each heme to the polypeptide backbone and the conformations the backbone can adopt. The thioether linkages, in turn, are perhaps needed, *inter alia*,^91^ to oppose the entropic penalty for organizing densely packed heme groups in a protein interior;^92^ multi-heme proteins with more than three hemes exclusively have *c*-instead of *b*-type hemes, where the difference is the presence of the thioether linkages.^93^

In a nutshell, electron transfer requires the hemes to be placed within tunneling distance (≤14 Å), and the closer the hemes are packed, the more readily endergonic steps can be tolerated and traversed.^67^ But to oppose the entropic penalty of closely packed hemes, each heme may be covalently linked through thioether bonds with Cys residues to the polypeptide backbone. These linkages, in addition to axial ligands donated to each heme Fe center from other residues, place geometrical constraints on the packing geometries. Most notably, the parallel-stacked geometry that is most favorable for heme-to-heme electron transfer cannot be geometrically accommodated for more than two consecutive heme pairs.^8^ The geometrically acceptable possibilities include a continuous chain of perpendicularly stacked hemes,^8^ which is not observed, or the typical alternating pattern of parallel and perpendicularly-stacked hemes. It is not clear why natural selection has favored the latter and if or when the former design is ever used. But either way, it is clear that electron transfer through multi-heme cytochromes is physically constrained and limited by the geometry of their construction.

### 2.9. Redox conduction through the CryoEM Structure at the Spectroelectrochemically Measured Potentials Can Sustain Cellular Respiration Rates

The set of k_ET_s for moving an electron through a unit cell of the OmcS filament can be used to evaluate the analytical Derrida formula^94^ for diffusive charge hopping along the periodic (pseudo)-one-dimensional heme chain of the cytochrome homopolymer. To set the stage for comparing the obtained charge diffusion constant to the demands of cellular respiration, it is useful to first discuss the cellular efflux of electrons that must be transported by cytochrome filaments *if* they are physiologically relevant.

#### 2.9.1. *G. sulfurreducens* Discharges ∼10^6^ electrons/s at -0.1 V

Measurements using micro- and nanoelectrodes, as well as a continuous flow microbial fuel cell have found respiration rates of 6.2 × 10^5^ – 1.2 × 10^6^ e^-^/s/cell (∼100 to 200 fA/cell) for both G*eobacter sulfurreducens* strain DL-1^1^ and *Shewanella oneidensis* strain MR-1.^4, 5^ An electron flux of 5(±2) × 10^5^ to 3 × 10^6^ e^-^/s/cell was measured for salt-respiring *G. sulfurreducens* in the lag and exponential growth phases, respectively, at 30°C and pH 7.4.^2, 6^ A rate of ∼9 × 10^6^ e^-^ /s/cell was measured for *G. sulfurreducens* growing on poorly crystalline Fe oxide.^3^ The yield coefficient (number of cells per mol electron) for *G. sulfurreducens* measured at voltages from - 0.003 to +0.397 V vs. SHE was ∼3 × 10^12^ to ∼3 × 10^11^ cells/(mol e^-^), respectively.^7^ The cellular efflux rate obtained by dividing these values of the yield coefficient into the specific growth rate of the bacterium (0.1 hours^-1^)^95^ and multiplying by Avogadro’s number gives (after unit conversion) 6 × 10^6^ to 6 × 10^7^ e^-^/s/cell. Taken together, the cellular respiratory rate is on the order of ∼10^6^ e^-^/s/cell regardless of whether *G. sulfurreducens* respires using electrodes, salts, or minerals as the terminal extracellular electron acceptor.

In the salt reduction experiments, the bacterium reduced complexes containing Ag^+^, Fe^3+^, Co^3+^, V^5+^, Cr^6+^, and Mn^7+^ ions with zero-order kinetics and the same (±2 × 10^5^) efflux rate. When cytochrome expression was downregulated or certain cytochromes were deleted by mutagenesis, the electron efflux rate was maintained by increasing the ratio of Fe^2+^/Fe^3+^ hemes in the remaining cytochromes. In all experiments, the observation of a nearly constant Fe^2+^-heme concentration during the salt reduction reactions demonstrated that the rate of Fe^2+^-heme oxidation by the metal salts equaled the rate of Fe^3+^-heme reduction by the respiratory machinery (specifically, an inner-membrane menaquinol dehydrogenase). The measurement therefore reflected the intrinsic cellular respiratory rate that the cell must maintain for ATP synthesis and homeostasis.

This finding is independent of whether cytochrome ‘nanowires’ were specifically expressed in the salt reduction experiments. How electrons exit the cell is a different question from how many electrons *need* to exit the cell. There is no indication, let alone evidence, to suggest that cellular respiration is markedly different under conditions that induce ‘nanowire’ expression; this point is discussed further below.

Another notable finding was that the respiration rate was the same (within experimental uncertainty) when reducing AgCl and AgNO_3_, even though the redox potentials of these salts range from ∼0.21 to 0.8 V versus normal hydrogen electrode (NHE).^6^ Indeed, much earlier voltammetry experiments indicated that *G. sulfurreducens* reaches its maximum respiratory rate on anodes posed at -0.1 V versus SHE and does not take advantage of the additional potential energy available from anodes at higher potentials.^96^

The experimental data summarized above suggests that *G. sulfurreducens* discharges ∼10^6^ e^-^ /s/cell at a potential of -0.1 V versus SHE.^96^ Assuming for concreteness that the efflux is ∼9 × 10^6^ e^-^/s/cell at this voltage, the needed conductance (G in Siemen (S)) for the electrical conduit is given by Ohm’s law (Eq. 8),

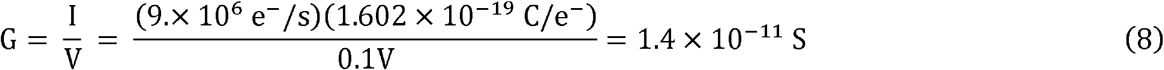

where I and V are respectively the current and voltage. If a cytochrome filament is hypothesized (and as recently proposed^29^) to be the only conductive channel out of the cell, the conductance can be related to a charge diffusion constant by Eq. 9, and thereby directly compared to the diffusion constants computed from NAMT and numerical simulations on the CryoEM structure.

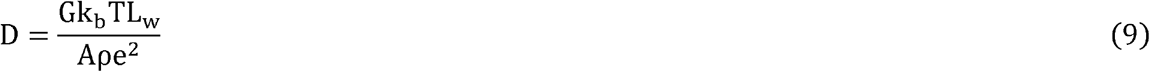

In Eq. 9, k_b_, e, and T are respectively the Boltzmann constant, the elementary charge, and absolute temperature, respectively; A and L_w_ are the cross-sectional area and length of the conduction channel; ρ is the charge density; and G is the conductance.

Assuming the conduction channel through the helical filament is cylindrical and that the charges are only transported through the heme chain, A = πr^2^. where r = 0.75 nm for the heme group. L_w_ is taken to be 300 nm, which is a typical electrode spacing bridged by cytochrome filaments in experiments. The ρ that is associated with maximal charge flux^36^ is 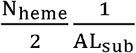, where N_heme_ is the number of hemes in a subunit of the homopolymer and L_sub_ is the length of a subunit. From the CryoEM-resolved structures of OmcS,^30, 31^ L_sub_ is 4.67 nm.

Evaluating Eq. 9 with the indicated values gives a diffusion constant of ∼1.0 nm^2^/ns. This value is the diffusion constant a *single* nanowire would need if it carried the *entire* cellular efflux. There are no firm estimates for the number of filaments/cell, but *G. sulfurreducens* is known to express >20 filaments/cell that were originally thought to be pili^44^ but now argued to be OmcS.^31^ Prior calculations indicate that the bacterium can synthesize 100 filaments/cell at a manageable bioenergetic cost.^45^ Regardless of the specific value, the G for a single ‘nanowire’ should be replaced by 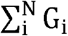, where N is the number of filaments/cell because the conductance of N parallel wires is additive. Thus, if the estimate of 100 filaments/cell is adopted, the required diffusion constant for cellular respiration on a per-filament basis is 0.01 nm^2^/ns/filament.

This value is still likely an overestimate because *G. sulfurreducens* has other pathways for discharging electrons (*e*.*g*., porin cytochrome complexes).^97^ If the cytochrome filaments are physiologically relevant, they would only need to carry some fraction of the total cellular current. The conclusion in the next sub-section is only strengthened by assuming an overestimated value.

#### 2.9.2. Tens to Hundreds of Filaments/Cell are Predicted to Suffice for Cellular Respiration

Figure 17a compares the computed diffusion constants for all 719 possible ΔG°landscapes obtained by differently assigning the six spectroelectrochemical potentials to the hemes of the CryoEM structure paired with the four sets of λ^rxn^ that account for different amounts of electronic polarization upon oxidation-state cycling. Figure 17b translates the results into the number of filaments/cell that would be needed to carry the entire cellular efflux given the predicted single-filament diffusion constant.

**Figure 17.**
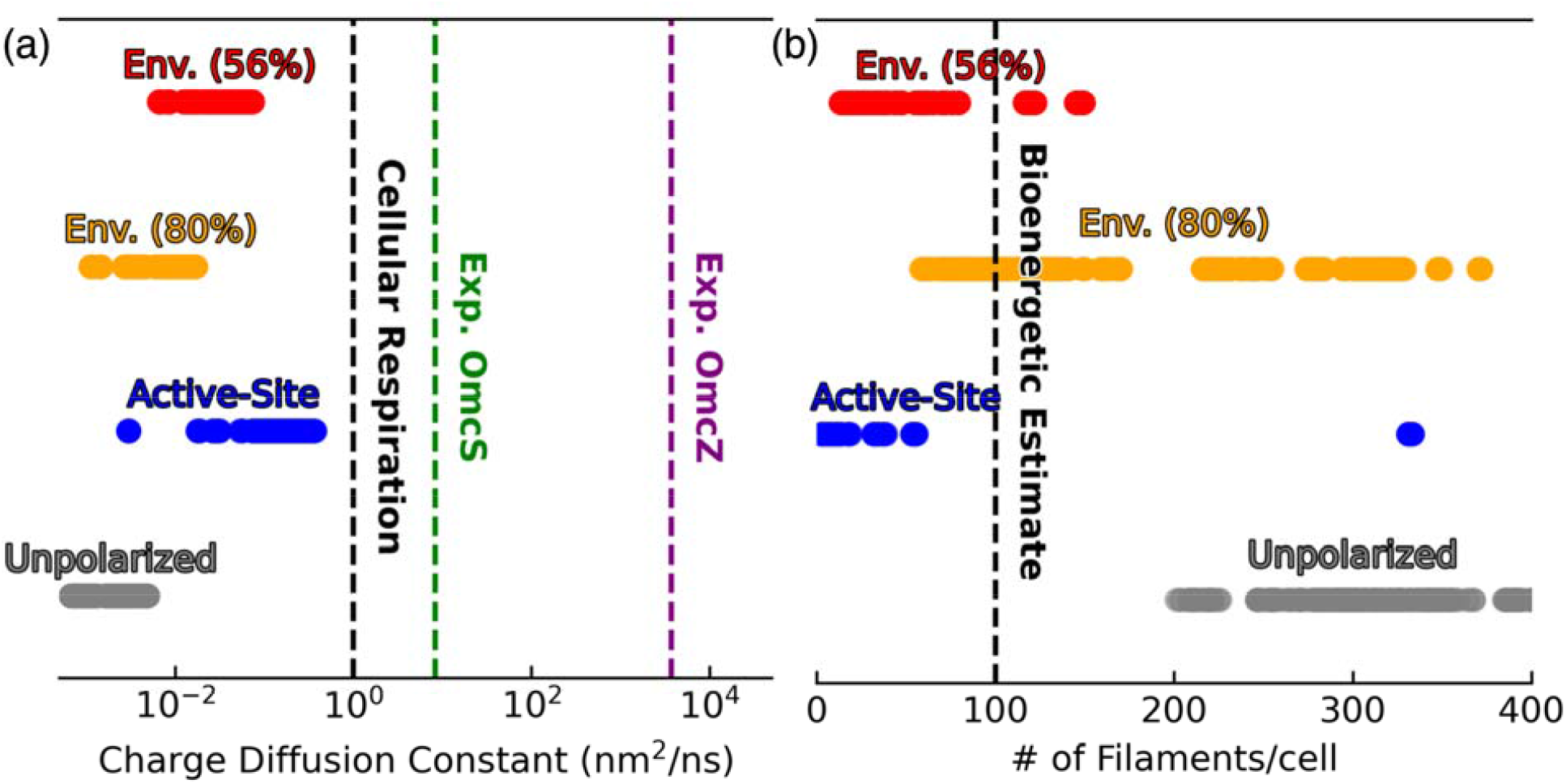
Distributions of predicted (a) charge diffusion constants and (b) the number of filaments/cell needed to discharge the cellular respiratory current. The distributions reflect 720 possible ways of assigning the previously reported spectroelectrochemical potentials to the hemes of the structure to define all possible free energy landscapes. These landscapes are also paired with four sets of reorganization energies that differ in whether and how they account for electronic polarization. Gray is for no account of electronic polarization; orange and blue respectively signify use of scaling factors of 0.80 and 0.56 to account for polarization of the environment, and red reflects polarization of the active site heme via Eq. 7. Also indicated are the diffusion constants shown in the main text to be needed for cellular respiration through an effective single filament, as well as diffusion constants computed with Eq. 10 for the conductivities reported for Omc-S and Z.

The figure indicates that a *minimum* of 201, 58, 13 or 3 OmcS filaments would be needed to carry the entire cellular efflux given the largest diffusion constant predicted with any assignment of the spectroelectrochemical potentials to specific hemes and λ^rxn^s that respectively neglect electronic polarization, include polarization of the environment by scaling λ^rxn^ by 0.80 or 0.56, or include active site polarization by Eq. 7. The predictions specifically using the computationally assigned ΔG° landscape of Figure 4 are 254, 70, 16, and 4 filaments/cell.

All of these estimates are in reasonable agreement with the observation of >20^44^ or calculated estimate of ∼100^45^ filaments/cell. The results are also in reasonable accord with a previous estimate of ≥7 filaments/cell obtained for a *generic* heme chain using the average of experimental time constants for electron transfer within parallel and perpendicularly-stacked hemes.^27^ Furthermore, a *G. sulfurreducens* cell is likely to express many more than the minimum number of filaments for reasons not related to intrinsic conductivity, but rather, for example, to maximize the likelihood of productive contacts between the filaments and a suitable electron acceptor.

Since a physiologically reasonable tens to hundreds of cytochrome filaments are predicted to be sufficient to sustain the entire cellular electrical efflux, the filaments are even more suited to this task when it is considered that *G. sulfurreducens* has additional mechanisms for discharging electrons,^97^ and the filaments would only have to carry a fraction of the total cellular current.

#### 2.9.3. Reported Conductivities from Electrical Measurements are Physiologically Irrelevant

Figure 17a also shows the diffusion constant obtained by evaluating Eq. 10 with the experimentally reported single-filament conductance of OmcS (G = 1.1 × 10^−10^ S)^35^ and OmcZ (G = 4.91 × 10^−8^ S)^43^ filaments and adjusting the geometrical parameters to be consistent with the CryoEM structure of either filament.

The diffusion constants obtained from electrical measurements on single OmcS and OmcZ filaments are 8.3 and 3700 nm^2^/ns, which are 10^1^ to 10^3^-fold larger than that required for the respiration of the entire cell. What use can a maximally respiring ∼10^2^ fA-generating cell at -0.1 V bias have for a filament capable of carrying 3 × 10^4^ fA as reported for OmcS? And why would the cell ever need to evolve an even more conductive filament capable of discharging 3 × 10^7^ fA at the same voltage?

One hypothesis is that the respiratory rate is dramatically increased under nanowire-expressing conditions. However, there is no indication or evidence to support this hypothesis. In fact, respiring on extracellular acceptors through ‘nanowires’ is *less efficient* for *G. sulfurreducens* growth than respiring on soluble species for the following reason:^98^ Organic carbon oxidation in the cytoplasm produces one proton per electron that is consumed during the reduction of a soluble terminal acceptor (*e*.*g*., fumarate) but remains trapped in the cytoplasm when electrons are discharged extracellularly. The build-up of protons in the cytoplasm reduces the electrochemical gradient across the inner-membrane that is required for ATP synthesis, and in fact, would entirely shut down respiration if a means of getting rid of some of the protons into the periplasm did not exist.^95, 99^

A second hypothesis is that the excessively conductive ‘nanowires’ are needed for communal life in biofilms. This may be the case, but biofilm growth on electrodes serving as infinite electron sinks is likely a laboratory artifact and irrelevant for the physiology of *G. sulfurreducens* in its natural habitat. If a natural electron sink, Fe(III) oxyhydroxide, occupies 50% of the space around a solitary *G. sulfurreducens* cell, bioenergetic calculations^45^ showed that the cell would need to reduce all available Fe(III) within a radius of 2 to 4 μm just to approach generating enough ATP for single-cell doubling. There is simply not enough oxidized mineral in a geographical area of the natural habitat for *G. sulfurreducens* to support a multi-cellular biofilm. It is also not clear what, if any, natural analog plays the role of switching on an electric field during anode growth to stimulate the expression of OmcZ filaments.^43^

A third hypothesis is that geometrical parameters for the CryoEM structure cannot be used in Eq. 10 with the reported conductivities because there is a substantial surface-induced structural change that is not captured by the solution-phase structure determination. However, filament height, helical parameters, and protein 2° structure content measured on surface-adsorbed cytochrome filaments are claimed to be consistent with the CryoEM structures.^33, 35^ The redox properties of OmcS in solution and solid-state were found to be similar,^29^ and the electrical conductivity of OmcS was determined at the same pH used to solve the CryoEM structure. ^35^

As already discussed elsewhere,^27^ the excessively large conductivities may reflect some abiological charge transport (instead of transfer) mechanism,^100^ or simply need to be corrected for the number of protein-electrode contacts. When the mobile electrode in conducting probe atomic force microscopy (CP-AFM) experiments, for example, spans ∼10 subunits of OmcS across it’s diameter, ascribing the current to a single protein-electrode contact does not seem appropriate.

Diagnosing why the experimental conductivities vastly exceed the requirements of cellular physiology further is beyond the scope of this work, but note that the inconsistency is between experimental findings, not theory and experiment. Indeed, by connecting spectroelectrochemical measurements to the CryoEM structure and applying the biologically relevant theory of redox conduction, the present work finds, in accord with bioenergetic calculations and experimental observations that tens-to-hundreds of the OmcS cytochrome filament are sufficient to discharge the respiratory flux of 10^6^ e^-^/s/cell from G. sulfurreducens

## 3. Conclusion

To summarize, a non-arbitrary comparison finds heme redox potentials computed from the CryoEM structure of OmcS and those reported 16-months later by spectroelectrochemistry to all agree within <0.04 eV. The consistent theory-experiment agreement enables an unambiguous assignment of the spectroelectrochemical redox potentials to specific hemes in the CryoEM structure, which is not presently possible by experimental techniques. The potentials are predicted to dynamically shift by ≤0.1 V due to redox cooperativities between the hemes, an effect that could also not be resolved experimentally but is consistent with prior literature. Because uncertainties in the potentials from both theory and experiment are within a factor of 2 of the maximal heme-heme interaction strength, the analysis generally confirms the finding of Portela *et al*. that the spectroelectrochemical titration curve can be effectively modeled as arising from six independent redox transitions, each dominated by a single heme. This independent heme approximation is justified by the close agreement with computations that assumed it and justifies the 1:1 mapping of spectroelectrochemical potentials to the hemes of the CryoEM structure. The mapping is a *prediction* that awaits experimental verification.

The structurally-assigned redox potentials define a reversible free energy ‘rollercoaster’ for electrons that pass through a unit cell of the homopolymer. When paired in the framework of non-adiabatic Marcus theory with computed electronic couplings and reorganization energies that do or do not include electronic polarization, a few pico-to hundreds of nanosecond heme-to-heme electron transfer rates are predicted. Slower rates are also anticipated for free energy landscapes based on different assignments of the spectroelectrochemical potentials to the hemes.

In general, however, any assignment of the spectroelectrochemical redox potentials to define a ΔG° landscape paired with a set λ^rxn^s that does or does not include the effect of electronic polarization gives rates that meet or exceed the typical millisecond timescale for enzymatic turnover.

A similar range of rates have been found by computational and experimental kinetic analyses of multi-heme systems that have the same heme packing geometries. The rates predicted for parallel-stacked hemes exceed experimental expectations, but those for the rate-limiting perpendicularly-stacked heme pairs agree more nearly with them.

The computed rates specify a diffusion constant that implies tens to hundreds of filaments are needed to discharge the 10^6^ e^-^/s metabolic flux of a *G. sulfurreducens* cell, which is consistent with the observed >20 or the ∼100 filaments/cell anticipated from bioenergetic calculations. The observation of >20 filaments/cell was made for what were once thought to be pili but are now argued to be OmcS ‘nanowires.’

Given that *G. sulfurreducens* does not likely use cytochrome ‘nanowires’ exclusively to discharge electrons and that the bacterium likely expresses many more than the theoretical *minimum* number of filaments for reasons unrelated to intrinsic ‘nanowire’ conductivity (e.g., the need to maximize productive contacts with oxidized minerals), the order-of-magnitude agreement is remarkable. The conclusion is that a bioenergetically manageable number of OmcS ‘nanowires’ have sufficient conductivity to discharge the entire cellular current if needed.

Vastly exceeding the needs of cellular respiration are the reported experimental conductivities of cytochrome ‘nanowires’ that can discharge 10^2^ – 10^5^ fA at -0.1 V *more* than generated by an entire cell. There is no indication or evidence that a *G. sulfurreducens* cell can dramatically oxidized more than ∼10^5^ molecules of acetate/s under ‘nanowire’-expressing growth. The cell would need to ‘burn’ at least 10-million molecules of acetate/s to make use of the reported conductivity for a *single* filament, let alone the combined conductivity of 100-such parallel ‘nanowires.’ The excessively large conductivities may have use in biofilm growth on the infinite electron sink provided by an electrode, but such electron acceptors do not exist in the natural habitat. A solitary *G. sulfurreducens* cell must reduce 25 to 50% of its biovolume in Fe oxide just to generate enough energy for single-cell doubling.^45^ There is unlikely enough oxidized mineral in one location of the natural habitat to support a multi-layer biofilm, meaning that even if the excessively large reported conductivities of ‘nanowires’ are used in that state, they have no bearing on the natural growth conditions.

The picture that emerges from prior bioenergetic^45^ calculations and the present study is: An isolated *G. sulfurreducens* cell must reach out and reduce minerals microns away just to survive. To do so, the cell expresses filaments that, if they have the structure resolved by CryoEM and are redox conductors as modeled here, can more than adequately discharge the cellular metabolic flux at a bioenergetically reasonable expense of synthesizing tens to hundreds of such filaments. This conclusion agrees with the fact that intracellular acetate oxidation and cytochrome reduction are the main rate-limiting processes for respiration compared to the extracellular transfer and interfacial discharge of electrons.^101^

The van der Waals packing of heme pairs in highly conserved geometries within binding sites that likely have a similar dielectric profile for accommodating the same redox cofactor imposes universal energetic and kinetic constraints on electron transfer through multi-heme cytochromes. This conclusion is supported by an analysis of all structurally characterized cytochrome ‘nanowires’ to date. Thus, a universal picture applicable to multi-heme chains in all kingdoms of life emerges from the foregoing computational analysis that connects cryogenic electron microscopy, spectroelectrochemistry, and cellular respiration.

## 4. Methods

### Simulation of Spectroelectrochemical Titration Curves

For a system like OmcS with non-to weakly interacting redox centers, the fraction of reduced species as a function of the solution potential (*i*.*e*., the spectroelectrochemical titration curve) can be approximated by a sum of independent Nernst equations (Eq. 11), as suggested by Portela *et al*.

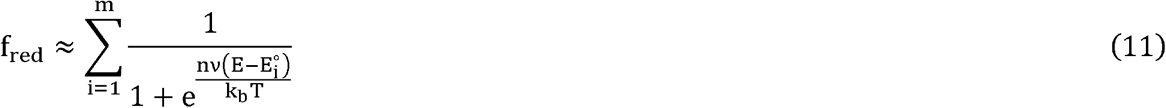

where 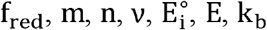, and T respectively signify the total reduced fraction, the number of redox-active centers (6 for OmcS), the Hill coefficient describing redox cooperativities in a *post hoc* fashion (assumed to be unity), the number of transferred electrons (each heme is a one-electron acceptor), the redox potential of the i^th^ heme (taken from either Dahl *et al*.^*35*^ or Guberman-Pfeffer^38^), the solution potential (−2 to +2 V), the Boltzmann constant, and absolute temperature (300 K). It is assumed here that all hemes experience the same voltage.

A Python program implementing Eq. 11 (CompareExpAndCalcRedoxTitrationAnalysis.py) was written to generate the data shown in Figure 2 and is available at the Zenodo repository accompanying this manuscript.^102^

### Computation of Redox Potentials

The QM/MM@MD methodology was extensively described in section S1.3.2 of the Supporting Information to Ref. 38; a brief description follows here for context. The six redox potentials compared in the present work to spectroelectrochemistry were presented in Table S11 of Ref. 38.

The redox potential of each heme was computed as

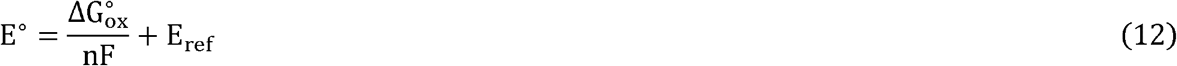

where n = 1 for the number of electrons accepted by each heme; F is the Faraday constant, and E_ref_ = 4.32 V to put the computed redox potentials on the same Standard Hydrogen Electrode (SHE) scale as the spectroelectrochemical potentials reported by Portela *et al*.^29^ The value used for E_ref_ is the absolute potential of SHE corrected for the integrated heat capacity and entropy of the electron.^103^ Very importantly, E_ref_ = 4.32 V was used for *all* computed redox potentials, meaning that the consistent level of agreement with experiment across the set of potentials, and the differences in computed redox potentials that govern electron transfer are independent of the particular value chosen for E_ref_.

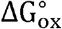 was computed by Eq. 13 under the approximation that the polarization of the environment (and therefore the free energy of solvation) is a linear function of the charge on the solute (*i*.*e*., the linear response approximation, LRA).

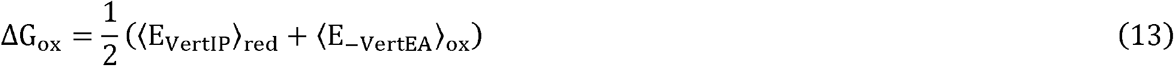

In Eq. 3, ⟨E_VertlP_⟩_red_ is the vertical ionization potential thermally averaged over configurations with the given heme in the reduced state, whereas ⟨E_-VertEA_⟩_ox_ is the negative of the vertical electron affinity computed over thermally averaged configurations with the given heme in the oxidized state. The vertical energy in either ensemble is given by the difference in energy between the oxidized and reduced electronic states at the same nuclear geometry, E_ox_(r_x_) − E_red_(r_x_), where x specifies the oxidized or reduced ensemble for the heme being considered in two oxidation states. Note that the free energy contribution from bulk solvation^103^ was omitted because it was found^38^ to be similar to thermal energy at 300 K.

To evaluate ⟨E_VertlP_⟩_red_ and ⟨E_−VertEA_⟩_ox_, classical molecular dynamics simulations were performed^38^ in which either all hemes were oxidized, or one heme was reduced while all other hemes remained oxidized. The all-heme-oxidized trajectory was propagated for 252 ns, whereas the single-heme-reduced trajectories were propagated for 301, 281, 266, 295, 242, and 294 ns, respectively, for Hemes #1 through #6. The AMBER FF99SB forcefield^104^ was used for the protein, whereas parameters for the heme cofactor were adopted from Crespo *et al*.^105^ and Henriques *et al*.^106^ The TIP3P water model^107^ and the monovalent ion parameters of Joung and Cheatham^108^ were used to model the solution state.

With 1.9 µs of simulations in hand, configurations for each of the six hemes in the fully oxidized ensemble, and the reduced heme in each of the single-heme-reduced ensembles were sampled and submitted to electronically-embedded QM/MM energy evaluations for the given heme in both the reduced and oxidized electronic states at the same nuclear geometry (*i*.*e*., two QM/MM single point calculations for each sampled configuration for each heme). The number of configurations sampled in the (all-heme-oxidized, single-heme-reduced) ensembles for each heme were: Heme #1 (100, 107); Heme #2 (154, 150); Heme #3 (131, 130); Heme #4 (128, 120); Heme #5 (139, 168); Heme #6 (112, 87). A total of (1526 frames) x (2 single − point calculations/frame) - 3052 calculations were performed on a 93-atom QM region (*i*.*e*., the cofactor shown in Figure 1 but with hydrogens present), embedded in the electrostatic environment of the protein-water matrix, using the B3LYP approximate density functional^109-111^ and a mixed, triple-ζ basis set: The LANL2TZ^112,113^ effective core potential and valance basis sets for Fe, and the 6-311G(d) basis set^114-116^ for H, C, N and S atoms. All QM calculations were performed using Gaussian 16 Rev. A.03.^117^

The distributions for both ⟨E_VertlP⟩red_ and ⟨E_−VertEA_⟩_oX_ calculated in this way were found to be symmetric, Gaussian-shaped, and of equal width, which are the conditions required by the LRA approximation.

### Computation of Heme Electronic Polarizability

The details are given in the *Assessment of Heme Electronic Polarizability* section of the Supporting Information for brevity here.

### Computation of Reorganization Energies

Reorganization energies were computed from Coulombic vertical energy gaps directly obtained from the fixed-point charge molecular dynamics simulations according to Eqs. 1 to 3 and 5.^34^ These reorganization energies were corrected for electronic polarization in the environment by applying recommended scaling factors of 0.56 or 0.80, or by adding to it the total (donor +acceptor) electronic polarization energy in the reactant and product states. The *Assessment of Heme Electronic Polarizability* and *Influence of Electronic polarization on Reorganization energy* sections of the Supporting Information discuss in detail how reorganization energies corrected for heme polarizability were obtained. These sections provide an annotated input script that was used to compute the Coulombic (fixed charge) contribution to the vertical energy gaps with CPPTRAJ of the AmberTools package,^118^ a configuration file for the TUPÃ python program^119^ to quantify the electric field exerted on the Fe centers, and the atomic coordinates of the quantum mechanically optimized heme active site models in the reduced and oxidized states used to compute the dipolar polarizability tensor in vacuum. A python script named PolarizabilityEffectOnReorganizationEnergy.py, and all the data needed to check its results are provided in the Zenodo repository associated with this manuscript.^102^

### Computation of Electronic Couplings

Previously reported^34^ electronic couplings used in the present analysis were computed with the diabatic state-based absolutely localized molecular orbital multi-state DFT 2 approach, as implemented in Q-Chem version 5.3.1,^120^ with the Perdew–Burke–Ernzerhof (PBE) approximate density functional^121^ and the def2-SVP basis set.^122^

### Computation of Redox Cooperativities

The BioDC Python workflow (Version 2.0)^47^ for computation of redox potentials, cooperativities, and conductivities was used for the second of these purposes here. The program is available on GitHub (https://github.com/Mag14011/BioDC), and all files related to this particular example case of OmcS are available at the Zenodo repository accompanying the manuscript.^102^

Experimental reports of redox cooperativities typically take the state in which all redox centers are reduced as the reference.^57^ The same all-heme-reduced state was taken as the reference for the redox cooperativity calculations. From this reference state, the difference in oxidation energy for heme i when heme j is oxidized versus reduced (ΔE) was computed for all possible pairings of hemes where i≠j; that is,

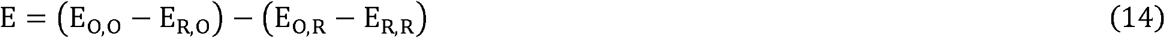

In Eq. 14, each subscript specifies the oxidation state of the i^th^ heme, followed by the j^th^ heme, where O = oxidized and R = reduced. ΔE indicates how much the oxidation of heme i is disfavored by the oxidation of heme j. Note that because the nuclear coordinates and atomic partial charges are the same in all energy terms of Eq. 14 except for hemes i and j, all Coulombic interactions other than between hemes i and j cancel. Furthermore, because the hemes are rather rigidly held in place by the protein matrix, with a standard deviation in the minimum edge-to-edge distance of 0.2 Å (Table S1), it is a reasonable approximation to compute the redox cooperativity for a single structure. This calculation was performed for the CryoEM structure PDB 6EF8. As a sanity check, the calculation was repeated for the alternate CryoEM structure PDB 6NEF. Interaction energies for both CryoEM structures agreed within 0.017 eV (less than thermal energy at 298 K). It is unlikely that thermal fluctuations will substantially change the reported interaction energies unless there is a large-scale conformational change in the filament.

BioDC provides an interface for computing the energy terms for the cooperativity calculation with the Poisson-Boltzmann Solvation Area model^123^ of the AmberTools package in which the protein is immersed in an implicit aqueous solvent.^118^ To that end, BioDC estimates the internal (*i*.*e*., protein) static dielectric constant based on the solvent accessibility^37^ of every adjacent heme pair and takes the average of these values as the static dielectric constant for all possible pairings of hemes in the redox cooperativity calculations.

## ASSOCIATED CONTENT

### Supporting Information

#### The Supporting Information is available free of charge at

Summary of metrical descriptors for heme packing in OmcS; assessment of heme electronic polarizability; protocol for assessing the influence of electronic polarization on reorganization energy; and electron transfer energetics for all structurally characterized cytochrome ‘nanowires.’

## AUTHOR INFORMATION

### Author Contributions

The research described herein was initiated and conducted solely by the corresponding author.

### Notes

The author declares no competing financial interest.

## ACKNOWLEDGMENT

All calculations were performed using the High Performance and Research Computing Services at Baylor University and personal computing resources.

## ABBREVIATIONS

AFM: atomic force microscopy
CryoEM: cryogenic electron microscopy
DFT: density functional theory
DHSE: Debye-Hückel shielded electrostatics
Mtr: metal-reducing
MD: molecular dynamics
Omc: outer-membrane cytochrome
QM/MM: quantum mechanics/molecular mechanics
redox: reduction-oxidation
SHE: standard hydrogen electrode.

