## Supporting Information for "Redox Conduction Through Cytochrome ‘Nanowires’ Can Sustain Cellular Respiration"

### Summary of Metrical Descriptors of Heme Packing in OmcS

Table S1. Metrical Parameters for the heme centers in OmcS from classical molecular dynamics simulations reported in Ref. ^1^. All the quantities are consistent with the highly conserved heme stacking geometries identified through a survey of >800 cytochromes in Ref. ^2^. Note that a single prime mark is used to indicate the heme in the next subunit of the filament.

| Heme Pair | Packing  Designation | Edge-to-Edge  Distance (Å) | Fe-to-Fe  Distance (Å) | Heme  Rotation (°) | Heme  Plane Tilt (°) | His-His Rotation (°) |
| --- | --- | --- | --- | --- | --- | --- |
| 1$\leftrightarrow$2 | Perpendicular | 6.0 ± 0.2 | 12.6 ± 0.2 | 130.7 ± 3.0 | 69.1 ± 2.6 | 12.0 ± 9.1, 28.3 ± 11.2 |
| 2$\leftrightarrow$3 | Parallel | 3.9 ± 0.2 | 9.4 ± 0.2 | 176.7 ± 1.7 | 11.0 ± 2.0 | 28.3 ± 11.2, 63.5 ± 10.1 |
| 3$\leftrightarrow$4 | Perpendicular | 6.0 ± 0.2 | 11.3 ± 0.2 | 143.4 ± 2.2 | 75.2 ± 2.3 | 63.5 ± 10.1, 42.5 ± 11.2 |
| 4$\leftrightarrow$5 | Parallel | 3.9 ± 0.2 | 9.5 ± 0.2 | 176.7 ± 1.8 | 8.0 ± 2.9 | 42.4 ± 11.2, 23,9 ± 12.0 |
| 5$\leftrightarrow$6 | Perpendicular | 5.9 ± 0.2 | 11.3 ± 0.2 | 142.2 ± 2.3 | 82.2 ± 2.7 | 23.9 ± 12.0, 66.4 ± 10.9 |
| 6$\leftrightarrow$1' | Parallel | 4.0 ± 0.2 | 9.7 ± 0.2 | 177.8 ± 1.6 | 7.7 ± 2.6 | 66.4 ± 10.9, 13.6 ± 8.2 |

### Correlation of Reaction Free Energies and Electronic Couplings in OmcS

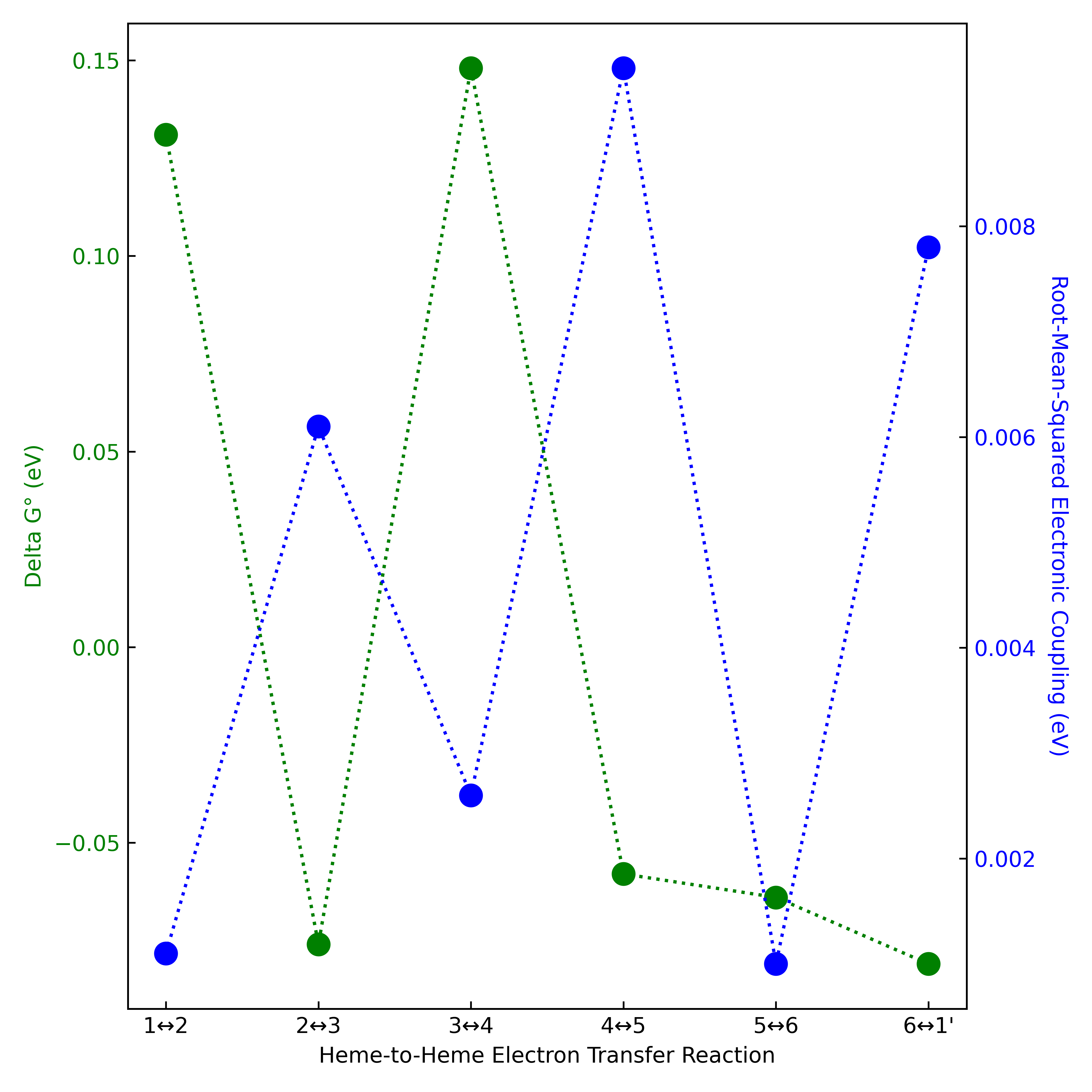

Figure S1. The root-mean-squared electronic coupling for adjacent hemes in the OmcS filament is typically largest for endergonic electron transfer reactions and smallest for exogenic reactions. This pattern is opposite to what was predicted for the deca-heme MtrF protein.^3^

### Comparison of Oxidized and Reduced Models of the Bis-His ligated *c*-type heme cofactor

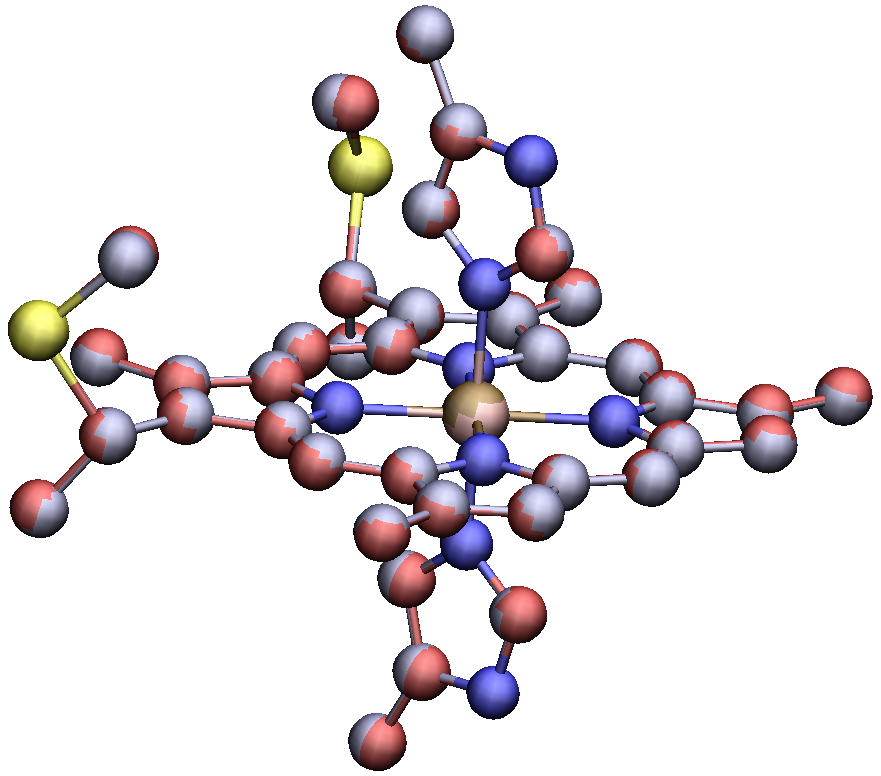

Figure S2. The BLYP/6-31G(d) optimized structures of the oxidized and reduced *c*-type bis-histidine-ligated heme cofactor of OmcS superimpose with a root-mean-squared coordinate deviation of ~0.04 Å, suggesting a minimal change in electronic structure consistent with low (0.05 to 0.08 eV) inner-sphere reorganization energy.^4, 5^ The reduced and oxidized cofactors are shown with red and blue-colored carbon atoms, respectively.

### Assessment of Heme Electronic Polarizability

The change in electronic polarizability between reduced and oxidized electronic states was computed (1) in vacuum for models of His-Met and His-His-ligated *c*-type hemes; (2) configurations of the His-Met-ligated *c*-type heme of cytochrome *c* sampled in the protein-water matrix during molecular dynamics; and (3) configurations for each of the six His-His-ligated c-type hemes of OmcS sampled in the protein-water matrix during molecular dynamics.

#### Calculations on Electronic Polarizability at Vacuum-Optimized Geometries

*Model hemes in Vacuum calculations.* Deachapunya *et al.*^6^ showed that the electronic polarizability of tetraphenylporphyrin-iron(III) chloride (FeTPPCl) can be quantitatively reproduced (Calc. 113.4 Å^3^ versus Exp. 102±11 Å^3^) by optimizing the geometry in vacuum with the BLYP functional and the 6-31G(d) basis set, followed by a polarizability calculation with the same functional but with diffusion functions added to the basis set (6-31+G(d)). Therefore, the models of His-Met- and His-His-ligated *c*-type heme cofactors shown in Figure S3 were optimized with the BLYP/6-31G(d) model chemistry. Each of the heme models was optimized in the reduced and oxidized electronic states. Harmonic frequency analyses confirmed that a local minimum on the ground state potential energy surface had been found in every case (*i.e.*, no negative frequencies). The atomic coordinates of the optimized geometries are provided in Tables S2 to S5. Subsequent calculation of the electronic polarizability was performed with a variety of approximate functional and basis set combinations, as shown in Table S6.

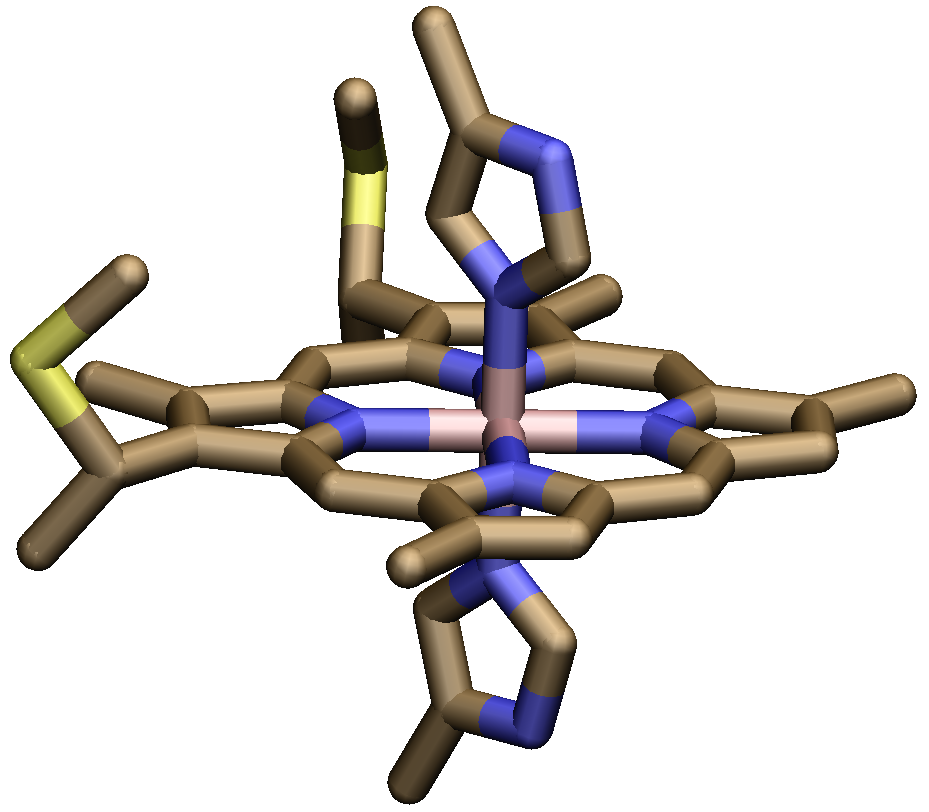

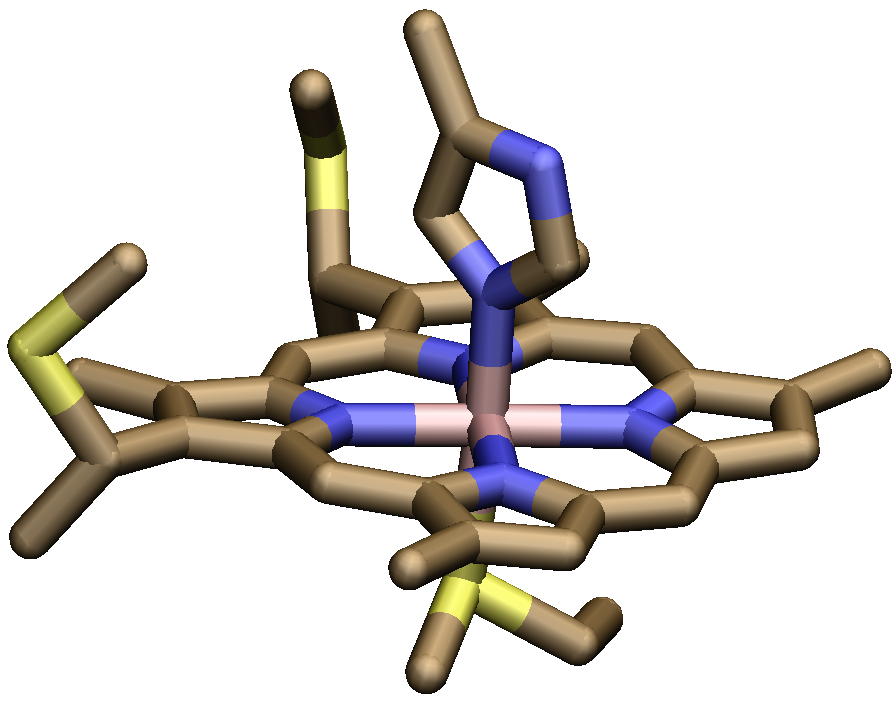

Figure S3. Model compounds for the His-His and His-Met ligated heme cofactors of OmcS and cytochrome *c*, respectively.

Table S2. Optimized structure of the reduced His-Met ligated *c*-type heme model at the BLYP/6-31G(d) level of theory. No negative frequencies were found. Self-consistent field energy = ‑4223.64340258 Hartrees. Coordinates are in Angstroms.

| Atom. | X (Å) | Y (Å) | Z (Å) |
| --- | --- | --- | --- |
| Fe | -0.139340 | 0.999937 | -0.106291 |
| N | -1.517070 | 2.414142 | 0.268039 |
| C | -1.308754 | 3.782076 | 0.443237 |
| C | -2.560625 | 4.454000 | 0.731386 |
| C | -3.549138 | 3.499872 | 0.759355 |
| C | -5.021784 | 3.687818 | 1.024040 |
| H | -5.251069 | 4.747036 | 1.224423 |
| H | -5.642044 | 3.370045 | 0.165546 |
| H | -5.364611 | 3.101803 | 1.896757 |
| C | -2.885016 | 2.229919 | 0.466965 |
| C | -3.544638 | 1.003194 | 0.365499 |
| H | -4.621900 | 1.002845 | 0.544820 |
| C | -2.954482 | -0.213202 | 0.008419 |
| N | -1.597840 | -0.387763 | -0.250879 |
| C | -3.694696 | -1.454658 | -0.185008 |
| C | -5.184239 | -1.602537 | 0.018249 |
| H | -5.459179 | -2.656046 | 0.173480 |
| H | -5.528093 | -1.038637 | 0.902840 |
| H | -5.761146 | -1.222814 | -0.846718 |
| C | -2.770318 | -2.403943 | -0.587726 |
| C | -2.991355 | -3.846218 | -0.981320 |
| H | -2.026778 | -4.304739 | -1.246585 |
| C | -3.940410 | -4.028717 | -2.191309 |
| H | -3.540242 | -3.475906 | -3.057665 |
| H | -4.024722 | -5.093477 | -2.465900 |
| H | -4.950234 | -3.644414 | -1.980972 |
| C | -1.466443 | -1.727672 | -0.601875 |
| C | -0.244243 | -2.346257 | -0.880928 |
| H | -0.264471 | -3.403663 | -1.147910 |
| C | 1.018621 | -1.748984 | -0.809948 |
| N | 1.248431 | -0.409405 | -0.507834 |
| C | 2.271564 | -2.468888 | -1.004880 |
| C | 2.374118 | -3.936928 | -1.345885 |
| H | 2.211391 | -4.124582 | -2.424445 |
| H | 3.362029 | -4.337738 | -1.075903 |
| H | 1.620117 | -4.532411 | -0.802402 |
| C | 3.288552 | -1.548186 | -0.809021 |
| C | 4.784143 | -1.740542 | -0.910409 |
| H | 5.290198 | -0.787071 | -0.695639 |
| C | 5.269681 | -2.221833 | -2.299756 |
| H | 4.946788 | -1.502721 | -3.071320 |
| H | 6.369876 | -2.291966 | -2.327419 |
| H | 4.853729 | -3.206709 | -2.561209 |
| C | 2.634605 | -0.272552 | -0.497042 |
| C | 3.295525 | 0.936813 | -0.256666 |
| H | 4.387064 | 0.932091 | -0.273057 |
| C | 2.684725 | 2.173046 | -0.032413 |
| N | 1.307927 | 2.385337 | 0.020712 |
| C | 3.415656 | 3.430052 | 0.126936 |
| C | 4.915215 | 3.590515 | 0.118801 |
| H | 5.364538 | 3.256816 | -0.834955 |
| H | 5.196046 | 4.646148 | 0.265052 |
| H | 5.400439 | 3.003274 | 0.920326 |
| C | 2.457379 | 4.404374 | 0.272705 |
| C | 1.161479 | 3.757310 | 0.220861 |
| C | -0.061413 | 4.411903 | 0.396415 |
| H | -0.036816 | 5.494834 | 0.548780 |
| C | 4.887100 | -2.192024 | 1.938104 |
| H | 3.788424 | -2.129773 | 1.942590 |
| H | 5.315495 | -1.188588 | 2.091748 |
| S | 5.496621 | -2.953571 | 0.370554 |
| C | 0.304210 | -1.435741 | 4.950645 |
| H | 0.281089 | -2.459805 | 4.548383 |
| H | -0.534041 | -1.340142 | 5.665018 |
| C | 0.205263 | -0.446722 | 3.825158 |
| N | 0.215378 | 0.937580 | 4.021139 |
| H | 0.288394 | 1.414332 | 4.917067 |
| C | 0.107540 | 1.555959 | 2.802010 |
| H | 0.091031 | 2.631398 | 2.662857 |
| N | 0.027867 | 0.639898 | 1.830059 |
| C | 0.088353 | -0.605871 | 2.457515 |
| H | 0.043705 | -1.524359 | 1.882166 |
| C | -2.421543 | -4.634555 | 1.738490 |
| H | -2.381599 | -3.565445 | 1.997062 |
| H | -2.747801 | -5.209804 | 2.618424 |
| S | -3.672612 | -4.942409 | 0.416655 |
| H | 2.610572 | 5.475152 | 0.416161 |
| H | -2.663260 | 5.526815 | 0.902215 |
| H | 1.243350 | -1.324351 | 5.523104 |
| H | 5.218427 | -2.850694 | 2.755472 |
| H | -1.420958 | -4.982598 | 1.435282 |
| S | -0.312138 | 1.233805 | -2.522218 |
| C | -1.539019 | 2.526851 | -3.097989 |
| C | 1.228594 | 2.023878 | -3.168651 |
| C | -2.931852 | 1.918891 | -3.322689 |
| H | -1.131955 | 2.927904 | -4.040748 |
| H | -1.564741 | 3.335750 | -2.353142 |
| H | 1.365556 | 3.028273 | -2.744048 |
| H | 2.069703 | 1.386649 | -2.866859 |
| H | 1.171817 | 2.069983 | -4.266961 |
| H | -3.333751 | 1.481041 | -2.396806 |
| H | -2.905157 | 1.133915 | -4.096019 |
| H | -3.629301 | 2.707620 | -3.655510 |

Table S3. Optimized structure of the oxidized His-Met ligated *c*-type heme model at the BLYP/6-31G(d) level of theory. No negative frequencies were found. Self-consistent field energy = ‑4223.47246036 Hartrees. Coordinates are in Angstroms.

| Atom. | X (Å) | Y (Å) | Z (Å) |
| --- | --- | --- | --- |
| Fe | -0.153552 | 0.994352 | -0.081875 |
| N | -1.539393 | 2.409397 | 0.266667 |
| C | -1.354247 | 3.789312 | 0.381193 |
| C | -2.606964 | 4.440891 | 0.687656 |
| C | -3.572398 | 3.466225 | 0.795021 |
| C | -5.035887 | 3.633921 | 1.111633 |
| H | -5.277524 | 4.692887 | 1.290259 |
| H | -5.677826 | 3.279589 | 0.285306 |
| H | -5.328912 | 3.066286 | 2.012747 |
| C | -2.892609 | 2.200516 | 0.521878 |
| C | -3.531716 | 0.961261 | 0.470062 |
| H | -4.599826 | 0.943299 | 0.691534 |
| C | -2.934424 | -0.244902 | 0.092555 |
| N | -1.587454 | -0.401892 | -0.219229 |
| C | -3.663941 | -1.490768 | -0.095748 |
| C | -5.141128 | -1.663205 | 0.158800 |
| H | -5.385444 | -2.718147 | 0.346799 |
| H | -5.470714 | -1.085163 | 1.038099 |
| H | -5.746193 | -1.319773 | -0.700761 |
| C | -2.744391 | -2.422973 | -0.560997 |
| C | -2.975545 | -3.859260 | -0.982683 |
| H | -2.020591 | -4.307903 | -1.296492 |
| C | -3.966909 | -4.003024 | -2.165364 |
| H | -3.608709 | -3.412741 | -3.025087 |
| H | -4.044343 | -5.057389 | -2.474084 |
| H | -4.974538 | -3.645854 | -1.903549 |
| C | -1.454853 | -1.738036 | -0.605166 |
| C | -0.236042 | -2.340189 | -0.924267 |
| H | -0.253065 | -3.389994 | -1.215387 |
| C | 1.019861 | -1.733370 | -0.843745 |
| N | 1.234974 | -0.394651 | -0.511775 |
| C | 2.278423 | -2.433600 | -1.040289 |
| C | 2.403673 | -3.888796 | -1.422109 |
| H | 2.322737 | -4.032583 | -2.515330 |
| H | 3.370003 | -4.298074 | -1.095133 |
| H | 1.614443 | -4.502057 | -0.956715 |
| C | 3.285881 | -1.506130 | -0.805719 |
| C | 4.786482 | -1.692324 | -0.885846 |
| H | 5.288745 | -0.745050 | -0.636634 |
| C | 5.288534 | -2.133531 | -2.283860 |
| H | 4.963494 | -1.404009 | -3.044389 |
| H | 6.388594 | -2.184388 | -2.296753 |
| H | 4.894718 | -3.120082 | -2.571203 |
| C | 2.618745 | -0.243474 | -0.481512 |
| C | 3.268096 | 0.970325 | -0.236139 |
| H | 4.358602 | 0.972697 | -0.237413 |
| C | 2.644336 | 2.205007 | -0.046511 |
| N | 1.266841 | 2.411328 | -0.013993 |
| C | 3.361164 | 3.472539 | 0.075210 |
| C | 4.857814 | 3.647991 | 0.080815 |
| H | 5.317436 | 3.291130 | -0.858326 |
| H | 5.125381 | 4.709158 | 0.199144 |
| H | 5.334399 | 3.088688 | 0.905579 |
| C | 2.390959 | 4.445663 | 0.166079 |
| C | 1.104993 | 3.791081 | 0.128349 |
| C | -0.122827 | 4.440386 | 0.281377 |
| H | -0.112539 | 5.527821 | 0.386846 |
| C | 5.023297 | -2.093204 | 1.958704 |
| H | 3.936888 | -1.944870 | 2.059025 |
| H | 5.544799 | -1.126955 | 2.048830 |
| S | 5.446720 | -2.939034 | 0.373142 |
| C | 0.403355 | -1.370471 | 5.000932 |
| H | 0.418232 | -2.400982 | 4.616632 |
| H | -0.438350 | -1.289942 | 5.711209 |
| C | 0.267363 | -0.402898 | 3.861120 |
| N | 0.233466 | 0.986097 | 4.037784 |
| H | 0.296456 | 1.474258 | 4.930265 |
| C | 0.099156 | 1.588623 | 2.822973 |
| H | 0.046728 | 2.661493 | 2.673044 |
| N | 0.044171 | 0.655453 | 1.862420 |
| C | 0.148432 | -0.583801 | 2.497375 |
| H | 0.129484 | -1.509817 | 1.933452 |
| C | -2.198874 | -4.872861 | 1.605191 |
| H | -2.035749 | -3.841062 | 1.953929 |
| H | -2.482032 | -5.500066 | 2.463747 |
| S | -3.605602 | -4.966424 | 0.414220 |
| H | 2.534643 | 5.521658 | 0.266238 |
| H | -2.728254 | 5.516173 | 0.820047 |
| H | 1.337560 | -1.210500 | 5.567531 |
| H | 5.370792 | -2.758670 | 2.763161 |
| H | -1.272039 | -5.274249 | 1.164938 |
| S | -0.367482 | 1.180412 | -2.570089 |
| C | -1.610490 | 2.466245 | -3.130849 |
| C | 1.172990 | 1.971101 | -3.213717 |
| C | -3.010325 | 1.860200 | -3.307387 |
| H | -1.221623 | 2.840924 | -4.091224 |
| H | -1.605921 | 3.291326 | -2.404145 |
| H | 1.295368 | 2.984654 | -2.808245 |
| H | 2.018772 | 1.344383 | -2.904081 |
| H | 1.113955 | 1.996402 | -4.311794 |
| H | -3.402626 | 1.457121 | -2.361673 |
| H | -3.007009 | 1.054174 | -4.058059 |
| H | -3.702350 | 2.646856 | -3.652734 |

Table S4. Optimized structure of the reduced His-His ligated *c*-type heme model at the BLYP/6-31G(d) level of theory. No negative frequencies were found. Self-consistent field energy = ‑3971.87419173 Hartrees. Coordinates are in Angstroms.

| Atom. | X (Å) | Y (Å) | Z (Å) |
| --- | --- | --- | --- |
| Fe | -0.069883 | -1.033084 | -0.006921 |
| N | -1.313557 | -2.582653 | -0.353997 |
| C | -0.976581 | -3.911491 | -0.604482 |
| C | -2.170115 | -4.711646 | -0.800746 |
| C | -3.254422 | -3.878138 | -0.671785 |
| C | -4.717543 | -4.224495 | -0.787164 |
| H | -4.849941 | -5.296670 | -1.006060 |
| H | -5.271168 | -4.004210 | 0.144394 |
| H | -5.216457 | -3.655977 | -1.593777 |
| C | -2.705432 | -2.550445 | -0.393187 |
| C | -3.481431 | -1.407532 | -0.196415 |
| H | -4.565097 | -1.530229 | -0.253472 |
| C | -2.998381 | -0.123412 | 0.071239 |
| N | -1.652929 | 0.211080 | 0.184635 |
| C | -3.853093 | 1.041286 | 0.271496 |
| C | -5.362236 | 1.017219 | 0.208632 |
| H | -5.767010 | 2.023200 | 0.024698 |
| H | -5.721433 | 0.364614 | -0.605990 |
| H | -5.810632 | 0.633982 | 1.145397 |
| C | -3.008682 | 2.110282 | 0.521017 |
| C | -3.352487 | 3.546743 | 0.839388 |
| H | -2.424986 | 4.122751 | 0.978147 |
| C | -4.205866 | 3.723745 | 2.119388 |
| H | -3.673547 | 3.283197 | 2.979230 |
| H | -4.382433 | 4.792067 | 2.329106 |
| H | -5.181778 | 3.221980 | 2.033128 |
| C | -1.641679 | 1.575778 | 0.454279 |
| C | -0.474796 | 2.326321 | 0.636914 |
| H | -0.586825 | 3.390720 | 0.848694 |
| C | 0.836423 | 1.838765 | 0.579932 |
| N | 1.181903 | 0.512372 | 0.339098 |
| C | 2.020817 | 2.668380 | 0.773267 |
| C | 1.994056 | 4.155087 | 1.040864 |
| H | 1.787502 | 4.384720 | 2.104062 |
| H | 2.953734 | 4.621624 | 0.773827 |
| H | 1.209632 | 4.658534 | 0.449212 |
| C | 3.112409 | 1.823269 | 0.659398 |
| C | 4.582575 | 2.138903 | 0.806101 |
| H | 5.170021 | 1.219863 | 0.660461 |
| C | 4.970610 | 2.724253 | 2.185958 |
| H | 4.674397 | 2.018290 | 2.980058 |
| H | 6.059716 | 2.885497 | 2.251333 |
| H | 4.466967 | 3.683395 | 2.380784 |
| C | 2.571447 | 0.487114 | 0.381595 |
| C | 3.339485 | -0.665072 | 0.186814 |
| H | 4.425054 | -0.567793 | 0.248788 |
| C | 2.849281 | -1.944112 | -0.083578 |
| N | 1.501905 | -2.275524 | -0.198216 |
| C | 3.698014 | -3.119220 | -0.285428 |
| C | 5.205038 | -3.143118 | -0.235125 |
| H | 5.593487 | -2.819567 | 0.748408 |
| H | 5.586657 | -4.159851 | -0.423745 |
| H | 5.657112 | -2.473297 | -0.990057 |
| C | 2.839228 | -4.164700 | -0.524017 |
| C | 1.487781 | -3.642440 | -0.469123 |
| C | 0.331028 | -4.406828 | -0.659367 |
| H | 0.458905 | -5.473664 | -0.865454 |
| C | 4.749930 | 2.473550 | -2.056783 |
| H | 3.659188 | 2.332087 | -2.089658 |
| H | 5.252564 | 1.497835 | -2.152587 |
| S | 5.252770 | 3.343097 | -0.507492 |
| C | -0.017358 | 1.439281 | -5.098340 |
| H | -0.117877 | 2.457636 | -4.693311 |
| H | -0.866766 | 1.269620 | -5.785520 |
| C | 0.009244 | 0.446534 | -3.972089 |
| N | 0.136878 | -0.931509 | -4.170766 |
| H | 0.222304 | -1.401517 | -5.069084 |
| C | 0.125687 | -1.553806 | -2.948321 |
| H | 0.210107 | -2.626503 | -2.812063 |
| N | -0.002514 | -0.649115 | -1.971878 |
| C | -0.075217 | 0.595029 | -2.600785 |
| H | -0.181492 | 1.507908 | -2.024676 |
| C | -3.113270 | 4.182743 | -1.967390 |
| H | -2.971810 | 3.106932 | -2.151223 |
| H | -3.579614 | 4.645044 | -2.850834 |
| S | -4.271343 | 4.458710 | -0.556698 |
| C | -0.410856 | -0.678071 | 5.630116 |
| H | -1.293029 | -1.103145 | 6.143567 |
| H | -0.522217 | 0.416863 | 5.633113 |
| C | -0.284045 | -1.169785 | 4.217139 |
| N | -0.126319 | -2.519656 | 3.888116 |
| H | -0.081713 | -3.294916 | 4.545470 |
| C | -0.041789 | -2.631397 | 2.523692 |
| H | 0.084231 | -3.571575 | 1.997846 |
| N | -0.136778 | -1.424225 | 1.955954 |
| C | -0.287266 | -0.512470 | 3.001948 |
| H | -0.386815 | 0.550126 | 2.808369 |
| H | 3.094354 | -5.207270 | -0.721272 |
| H | -2.171689 | -5.782432 | -1.011500 |
| H | 0.908519 | 1.411798 | -5.702033 |
| H | 0.478385 | -0.923148 | 6.239807 |
| H | 5.059879 | 3.118962 | -2.892921 |
| H | -2.137418 | 4.661729 | -1.786626 |

Table S5. Optimized structure of the oxidized His-His ligated *c*-type heme model at the BLYP/6-31G(d) level of theory. No negative frequencies were found. Self-consistent field energy = ‑3971.71045563 Hartrees. Coordinates are in Angstroms.

| Atom. | X (Å) | Y (Å) | Z (Å) |
| --- | --- | --- | --- |
| Fe | -0.067204 | -1.029903 | -0.007915 |
| N | -1.294859 | -2.603492 | -0.351103 |
| C | -0.952174 | -3.932922 | -0.594821 |
| C | -2.141792 | -4.732230 | -0.785182 |
| C | -3.229858 | -3.900125 | -0.660036 |
| C | -4.690611 | -4.251807 | -0.773309 |
| H | -4.818322 | -5.324864 | -0.983583 |
| H | -5.241053 | -4.025328 | 0.157313 |
| H | -5.186517 | -3.690336 | -1.585175 |
| C | -2.684731 | -2.569569 | -0.388504 |
| C | -3.464716 | -1.429365 | -0.197724 |
| H | -4.546594 | -1.557469 | -0.254953 |
| C | -2.989634 | -0.141493 | 0.063273 |
| N | -1.650230 | 0.209988 | 0.175990 |
| C | -3.853127 | 1.016613 | 0.258608 |
| C | -5.360664 | 0.978824 | 0.202964 |
| H | -5.771249 | 1.976104 | -0.007952 |
| H | -5.720179 | 0.300569 | -0.588706 |
| H | -5.794829 | 0.622979 | 1.155809 |
| C | -3.017317 | 2.097661 | 0.503256 |
| C | -3.382011 | 3.534901 | 0.812875 |
| H | -2.462762 | 4.127615 | 0.937970 |
| C | -4.221128 | 3.697870 | 2.105608 |
| H | -3.678182 | 3.260012 | 2.959910 |
| H | -4.400314 | 4.764152 | 2.315912 |
| H | -5.196119 | 3.192968 | 2.029218 |
| C | -1.650937 | 1.578629 | 0.440946 |
| C | -0.491873 | 2.338547 | 0.623804 |
| H | -0.611699 | 3.401867 | 0.830417 |
| C | 0.819268 | 1.854843 | 0.571873 |
| N | 1.168525 | 0.527062 | 0.333011 |
| C | 2.000342 | 2.681358 | 0.768026 |
| C | 1.974942 | 4.165821 | 1.042772 |
| H | 1.818630 | 4.383192 | 2.115743 |
| H | 2.919455 | 4.636656 | 0.735227 |
| H | 1.163478 | 4.668088 | 0.490167 |
| C | 3.094447 | 1.834094 | 0.656670 |
| C | 4.565604 | 2.158226 | 0.808331 |
| H | 5.161392 | 1.245294 | 0.656973 |
| C | 4.937074 | 2.731156 | 2.199312 |
| H | 4.628916 | 2.023826 | 2.987328 |
| H | 6.025460 | 2.883323 | 2.272971 |
| H | 4.440868 | 3.693957 | 2.394354 |
| C | 2.557012 | 0.499269 | 0.378358 |
| C | 3.330246 | -0.649416 | 0.187211 |
| H | 4.413819 | -0.544733 | 0.250196 |
| C | 2.849517 | -1.931937 | -0.079297 |
| N | 1.508303 | -2.280385 | -0.195529 |
| C | 3.707613 | -3.100674 | -0.274407 |
| C | 5.213486 | -3.113373 | -0.221273 |
| H | 5.593238 | -2.781988 | 0.761789 |
| H | 5.601455 | -4.127402 | -0.403433 |
| H | 5.657620 | -2.444651 | -0.980252 |
| C | 2.856567 | -4.155656 | -0.509996 |
| C | 1.503980 | -3.649347 | -0.460260 |
| C | 0.355540 | -4.424847 | -0.648135 |
| H | 0.489730 | -5.490583 | -0.848750 |
| C | 4.921197 | 2.416000 | -2.042597 |
| H | 3.854701 | 2.183250 | -2.185295 |
| H | 5.513282 | 1.487018 | -2.050921 |
| S | 5.200574 | 3.377127 | -0.491125 |
| C | -0.012112 | 1.418248 | -5.116173 |
| H | -0.096467 | 2.442070 | -4.722891 |
| H | -0.868895 | 1.247362 | -5.791793 |
| C | 0.010724 | 0.436265 | -3.980776 |
| N | 0.123460 | -0.946657 | -4.168476 |
| H | 0.198984 | -1.422117 | -5.066601 |
| C | 0.113303 | -1.563463 | -2.952781 |
| H | 0.187783 | -2.636290 | -2.813259 |
| N | -0.000692 | -0.648315 | -1.980725 |
| C | -0.064783 | 0.595978 | -2.611402 |
| H | -0.158880 | 1.513749 | -2.042221 |
| C | -3.098269 | 4.304676 | -1.951776 |
| H | -2.856272 | 3.260911 | -2.205940 |
| H | -3.572100 | 4.784627 | -2.821169 |
| S | -4.330054 | 4.393440 | -0.580133 |
| C | -0.420680 | -0.650464 | 5.630675 |
| H | -1.301875 | -1.082345 | 6.137320 |
| H | -0.535766 | 0.443497 | 5.636261 |
| C | -0.289747 | -1.141583 | 4.218131 |
| N | -0.126570 | -2.493151 | 3.890143 |
| H | -0.080789 | -3.265945 | 4.552760 |
| C | -0.038954 | -2.615354 | 2.535418 |
| H | 0.091252 | -3.558183 | 2.015739 |
| N | -0.137203 | -1.407164 | 1.963944 |
| C | -0.292988 | -0.487117 | 3.002719 |
| H | -0.396051 | 0.574109 | 2.807222 |
| H | 3.121250 | -5.195780 | -0.701686 |
| H | -2.141288 | -5.803061 | -0.990159 |
| H | 0.908960 | 1.367930 | -5.723429 |
| H | 0.469142 | -0.896451 | 6.236819 |
| H | 5.262064 | 3.059777 | -2.867288 |
| H | -2.177963 | 4.856393 | -1.701121 |

Table S6. Iso- and anisotropic polarizabilities $\left( \alpha\right)$, and the anisotropy parameter $\left( \kappa=\frac{\alpha_{\mathrm{aniso}}}{3*\alpha_{\mathrm{iso}}} \right)$ computed in vacuum for His-Met and His-His ligated *c*-type heme models with various approximate density functionals and basis sets. All calculations were performed using Gaussian 16, Revision A.03. A common vacuum optimized geometrical structure in a given redox state was used for all polarizability calculations.

| Model Chemistry | $\alpha_{iso,red}$ | $\alpha_{iso,ox}$ | $\Delta\alpha_{\mathrm{iso}}$ | $\alpha_{aniso,red}$ | $\alpha_{aniso,ox}$ | $\Delta\alpha_{\mathrm{aniso}}$ | $\kappa_{\mathrm{red}}$ | $\kappa_{\mathrm{ox}}$ |
| --- | --- | --- | --- | --- | --- | --- | --- | --- |
| His-His Ligated Heme | | | | | | | | |
| CAM-B3LYP/BS1 | 86.55 | 86.94 | 0.39 | 39.80 | 39.31 | -0.49 | 0.15 | 0.15 |
| B3LYP/BS1 | 88.84 | 89.62 | 0.78 | 40.30 | 40.28 | -0.02 | 0.15 | 0.15 |
| BLYP/BS1 | 91.92 | 98.46 | 6.54 | 39.37 | 49.25 | 9.88 | 0.14 | 0.17 |
| PBE/BS1 | 92.12 | 98.09 | 5.97 | 39.10 | 48.26 | 9.16 | 0.14 | 0.16 |
| BLYP/6-31G(d) | 92.25 | 96.24 | 3.99 | 38.29 | 46.46 | 8.17 | 0.14 | 0.16 |
| BLYP/6-31+G(d) | 102.60 | 107.53 | 4.93 | 39.59 | 50.48 | 10.89 | 0.13 | 0.16 |
| BLYP/6-311+G(d) | 102.99 | 108.45 | 5.46 | 39.59 | 51.08 | 11.49 | 0.13 | 0.16 |
| BLYP/6-311+G(d,p) | 103.67 | 109.07 | 5.40 | 39.60 | 51.15 | 11.55 | 0.13 | 0.16 |
| BLYP/6-311+G(2d,p) | 104.67 | 109.89 | 5.22 | 39.69 | 51.12 | 11.43 | 0.13 | 0.16 |
| His-Met Ligated Heme | | | | | | | | |
| CAM-B3LYP/BS1 | 85.76 | 87.03 | 1.27 | 43.30 | 40.97 | -2.33 | 0.17 | 0.16 |
| B3LYP/BS1 | 87.99 | 89.64 | 1.65 | 44.01 | 42.74 | -1.27 | 0.17 | 0.16 |
| BLYP/BS1 | 90.99 | 99.76 | 8.77 | 43.29 | 54.62 | 11.33 | 0.16 | 0.18 |
| PBE/BS1 | 91.26 | 99.23 | 7.97 | 42.91 | 53.24 | 10.33 | 0.16 | 0.18 |
| BLYP/6-31G(d) | 91.16 | 96.76 | 5.60 | 42.51 | 50.89 | 8.38 | 0.16 | 0.18 |
| BLYP/6-31+G(d) | 101.38 | 108.62 | 7.24 | 43.59 | 56.02 | 12.43 | 0.14 | 0.17 |
| BLYP/6-311+G(d) | 101.85 | 109.68 | 7.83 | 43.64 | 56.76 | 13.12 | 0.14 | 0.17 |
| BLYP/6-311+G(d,p) | 102.51 | 110.24 | 7.73 | 43.74 | 56.79 | 13.05 | 0.14 | 0.17 |
| BLYP/6-311+G(2d,p) | 103.50 | 110.95 | 7.45 | 43.87 | 56.64 | 12.77 | 0.14 | 0.17 |

BS1 = Fe = LANL2DZ; H, C, N, S = 6-31G(d)

#### Calculations of Electronic Polarizability on Molecular Dynamics-Generated Configurations

*His-His-Ligated c-type Hemes of OmcS.* A set of 250 snapshots were randomly selected with a minimum time increment of >20 ps from molecular dynamics trajectories in which each of the six hemes were in the reduced state and all other hemes were in the oxidized state. At each frame, the electronic polarizability was computed in the oxidized and reduced electronic states. The polarizability was computed with the BLYP/6-31G(d) model chemistry. The results are summarized in Tables S7 and S9; the polarizability tensor computed at each frame is provided in PerFramePolarizabilities.xlsx.

*His-Met-Ligated c-type Heme of Cytochrome c.* The polarizability calculations for a His-Met-ligated heme were repeated, but now at geometries sampled during classical molecular dynamics within the cytochrome *c* active site. Separate molecular dynamics trajectories were propagated with the charge distribution on the heme corresponding to the reduced and oxidized states. From each of these ensembles, 200 snapshots were selected randomly so long as the minimum time increment between the snapshots was >20 ps. The polarizability was computed for the heme at each configuration in the oxidized electronic state with and without the protein-water environment, as well as in the reduced electronic state with and without the protein-water environment. The results are summarized in Tables S7 and S8; the polarizability tensor computed at each frame is provided in PerFramePolarizabilities.xlsx. The polarizability calculations were performed with the BLYP functional, the LANL2DZ effective core and valance basis sets for Fe, and the 6-31G(d) basis set for H, C, N, and S atoms. The molecular dynamics simulations for cytochrome *c* used to generate the sampled configurations were prepared and performed as described in the following paragraphs.

Table S7. Iso- and anisotropic polarizabilities $\left( \alpha\right)$, and the anisotropy parameter $\left( \kappa=\frac{\alpha_{\mathrm{aniso}}}{3*\alpha_{\mathrm{iso}}} \right)$ computed in vacuum for the His-His and His-Met ligated hemes of cytochrome *c* and OmcS, both in vacuum and within the protein-water matrix. The calculations were performed over 200 (cytochrome *c*) or 250 (OmcS) configurations from either the reduced or oxidized ensemble (Ensb.) from classical molecular dynamics for the heme being analyzed. The polarizability was computed using the BLYP function with either the BS1 basis set (LANL2DZ for Fe and 6-31G(d) for H, C, N, and S) (cytochrome *c*) or the 6-31G(d) basis set for all atoms (OmcS). The latter basis set underestimates the oxidized-reduced polarizability difference predicted by the former basis set by 10-30% (Table S7). The table gives a summary view of ‘raw data’ provided in Tables S13 – S32. All polarizability calculations were performed with Gaussian 16, Revision A.03 and analyzed with MultiWFN.

| Structure | $\alpha_{iso,red}$ | $\alpha_{iso,ox}$ | $\Delta\alpha_{\mathrm{iso}}$ | $\alpha_{aniso,red}$ | $\alpha_{aniso,ox}$ | $\Delta\alpha_{\mathrm{aniso}}$ | $\kappa_{\mathrm{red}}$ | $\kappa_{\mathrm{ox}}$ |
| --- | --- | --- | --- | --- | --- | --- | --- | --- |
| Cyt. *c* (Red. Ensb., vac.) | 88.8±0.5 | 96.4±2.5 | 7.6±2.6 | 41.4±1.3 | 49.9±4.0 | 8.5±4.0 | 0.16 | 0.17 |
| Cyt. *c* (Ox. Ensb. Vac.) | 88.7±0.6 | 95.9±2.2 | 7.3±2.3 | 40.9±1.1 | 48.9±3.2 | 8.8±3.4 | 0.15 | 0.17 |
| Cyt. *c* (Red. Ensb., Env.) | 88.5±0.6 | 94.6±2.4 | 6.1±2.5 | 41.6±1.1 | 47.8±3.8 | 6.2±4.0 | 0.16 | 0.17 |
| Cyt. *c* (Ox. Ensb. Env.) | 88.8±0.6 | 93.1±2.0 | 4.3±2.1 | 41.3±1.2 | 44.7±3.0 | 3.4±3.2 | 0.16 | 0.16 |
| OmcS Heme #1 (Red. Ensb. Env.) | 90.3±0.7 | 91.7±0.8 | 1.4±1.8 | 38.1±1.2 | 40.5±1.3 | 2.4±0.8 | 0.14 | 0.15 |
| OmcS Heme #2 (Red. Ensb. Env.) | 90.3±0.6 | 92.1±1.0 | 1.8±2.0 | 39.1±1.2 | 43.0±1.6 | 4.0±1.0 | 0.14 | 0.16 |
| OmcS Heme #3 (Red. Ensb. Env.) | 90.0±0.6 | 91.3±0.8 | 1.3±1.8 | 37.9±1.2 | 39.2±1.4 | 1.3±0.8 | 0.14 | 0.14 |
| OmcS Heme #4 (Red. Ensb. Env.) | 90.4±0.6 | 92.3±0.9 | 1.9±2.0 | 39.2±1.2 | 42.0±1.6 | 2.8±0.9 | 0.14 | 0.15 |
| OmcS Heme #5 (Red. Ensb. Env.) | 90.3±0.6 | 91.6±0.8 | 1.3±2.4 | 38.5±1.6 | 41.5±1.8 | 3.0±0.8 | 0.14 | 0.15 |
| OmcS Heme #6 (Red. Ensb. Env.) | 90.3±0.7 | 91.9±0.9 | 1.6±2.1 | 38.1±1.4 | 41.0±1.6 | 2.9±0.9 | 0.14 | 0.15 |

Table S8. Polarizability $\left( \alpha\right)$ tensors in upper-right triangle form for the reduced and oxidized state for configurations of the His-Met ligated heme of cytochrome *c*, considered both with and without the surrounding protein-water matrix. All configurations were generated by molecular dynamics with the protein-water matrix present. The BLYP functional, LANL2DZ basis set for Fe, and 6-31G(d) basis set for H, C, N, and S were used for the polarizability calculations, which were performed with Gaussian 16, Revision A.03 and analyzed with MultiWFN.

| Vacuum | | | | | | | | |
| --- | --- | --- | --- | --- | --- | --- | --- | --- |
| Reduced Ensemble | | | | | | | | |
| $\boldsymbol{\alpha}_{\mathbf{ox}}$ | | | $\boldsymbol{\alpha}_{\mathbf{red}}$ | | | $\boldsymbol{\Delta\alpha}$ | | |
| 100.5±11.7 | 1.7±14.4 | 2.3±10.8 | 92.0±9.6 | 1.0±11.8 | 2.4±9.2 | 8.5±15.2 | 0.7±18.6 | -0.1±14.2 |
|  | 91.0±15.6 | -1.1±14.0 |  | 84.5±12.9 | -1.1±11.4 |  | 6.5±20.2 | 0.0±18.1 |
|  |  | 97.7±14.6 |  |  | 89.8±11.8 |  |  | 7.9±18.7 |
| Oxidized Ensemble | | | | | | | | |
| $\boldsymbol{\alpha}_{\mathbf{ox}}$ | | | $\boldsymbol{\alpha}_{\mathbf{red}}$ | | | $\boldsymbol{\Delta\alpha}$ | | |
| 92.1±14.5 | -4.8±13.3 | 1.5±13.9 | 87.7±12.2 | -3.9±10.5 | 1.1±11.7 | 7.3±19.0 | -0.9±16.9 | 0.4±18.2 |
|  | 97.5±12.8 | -2.9±11.3 |  | 89.7±11.1 | -2.4±9.3 |  | 7.8±17.0 | -0.5±14.6 |
|  |  | 98.2±12.9 |  |  | 91.6±10.8 |  |  | 6.7±16.8 |
| In Protein-Water Environment | | | | | | | | |
| Reduced Ensemble | | | | | | | | |
| $\boldsymbol{\alpha}_{\mathbf{ox}}$ | | | $\boldsymbol{\alpha}_{\mathbf{red}}$ | | | $\boldsymbol{\Delta\alpha}$ | | |
| 98.3±10.9 | 1.6±13.8 | 2.1±10.6 | 91.8±9.7 | 1.0±11.8 | 2.4±9.3 | 6.6±14.5 | 0.7±18.2 | -0.3±14.1 |
|  | 89.6±15.1 | -1.2±13.2 |  | 84.2±12.9 | -1.1±11.4 |  | 5.3±19.9 | -0.1±17.5 |
|  |  | 95.8±14.1 |  |  | 89.6±11.8 |  |  | 6.2±18.4 |
| Oxidized Ensemble | | | | | | | | |
| $\boldsymbol{\alpha}_{\mathbf{ox}}$ | | | $\boldsymbol{\alpha}_{\mathbf{red}}$ | | | $\boldsymbol{\Delta\alpha}$ | | |
| 89.5±13.2 | -4.6±12.4 | 1.4±12.2 | 84.8±12.3 | -3.9±105 | 1.1±11.8 | 4.6±18.1 | -0.6±16.2 | 0.3±1.8 |
|  | 95.1±12.6 | -3.0±10.2 |  | 89.7±11.2 | -2.4±9.4 |  | 5.4±16.8 | -0.6±13.8 |
|  |  | 94.7±11.4 |  |  | 91.8±10.9 |  |  | 2.9±15.8 |

Table S9. Polarizability $\left( \alpha\right)$ tensors in upper triangle form for the reduced and oxidized states over 250 configurations for each His-His ligated heme of OmcS, considered with the protein-water matrix present. All configurations were generated by molecular dynamics with the protein-water matrix present. The polarizability was computed using the BLYP/6-31G(d) model chemistry as implemented in Gaussian 16, Revision A.03. The results were analyzed with MultiWFN.

| Heme # | Oxidized | | | Reduced | | | Difference | | |
| --- | --- | --- | --- | --- | --- | --- | --- | --- | --- |
| 1 | -75.2 | -16.2 | -9.0 | -74.5 | -9.3 | -8.4 | 0.7 | -0.9 | -0.6 |
|  |  | 97.7 | -1.4 |  | 95.6 | -1.5 |  | 1.9 | 0.2 |
|  |  |  | 102.1 |  |  | 100.5 |  |  | 1.6 |
| 2 | 100.3 | -4.4 | -13.1 | 98.5 | -4.2 | -12.2 | 1.8 | -0.2 | -0.9 |
|  |  | 99.7 | -13.1 |  | 96.5 | -11.4 |  | 3.2 | -1.7 |
|  |  |  | 76.4 |  |  | 75.8 |  |  | 0.5 |
| 3 | 89.1 | 8.1 | 16.2 | 88.0 | 8.1 | 15.7 | 1.1 | 0.0 | 0.5 |
|  |  | 97.0 | -11.2 |  | 95.3 | -10.8 |  | 1.7 | -0.4 |
|  |  |  | 87.7 |  |  | 86.7 |  |  | 1.0 |
| 4 | 83.0 | -11.6 | 14.5 | 82.1 | -11.6 | 13.1 | 0.9 | 0.0 | 1.4 |
|  |  | 96.4 | 7.4 |  | 94.9 | 6.4 |  | 1.5 | 1.0 |
|  |  |  | 97.6 |  |  | 94.2 |  |  | 3.4 |
| 5 | 104.2 | 0.3 | 0.7 | 102.2 | 0.5 | 0.6 | 2.0 | -0.1 | 0.1 |
|  |  | 71.5 | 12.8 |  | 71.6 | 11.8 |  | -0.1 | 1.1 |
|  |  |  | 99.2 |  |  | 97.2 |  |  | 2.0 |
| 6 | 99.3 | 7.5 | 5.5 | 98.1 | 7.7 | 4.7 | 1.2 | -0.2 | 0.8 |
|  |  | 90.8 | -18.7 |  | 88.8 | -16.9 |  | 2.0 | -1.8 |
|  |  |  | 85.6 |  |  | 84.1 |  |  | 1.5 |

*Structure setup and parameterization of Cytochrome c for Molecular Dynamics.* Crystal structures of oxidized (PDB 1AKK)^7^ and reduced (PDB 1GIW)^8^ horse heart cytochrome *c* were prepared with the assistance of the Structure Preparation and Relaxation module of the BioDC program. In this process, the protein was centered in a water box with 20 Å-thick padding on all sides and a sufficient number of counterions for charge neutrality (7 or 8 Cl^-^ ions in the reduced and oxidized states, respectively). An additional step (not part of the BioDC module) was taken to acylate the N-terminus of the protein.

The parameters for the His-Met ligated *c*-type heme were in part taken from Crespo *et a*.^9^ and developed with the Metal Center Parameter Builder (MCPB) program^10^ of the AmberTools suite^11^ based on quantum chemical calculations using the B3LYP approximate density functional and a mixed basis set (LANL2TZ(f) for Fe and 6-31G(d) for all second-row elements). Gaussian 16 Rev. A.03 was used for the quantum chemical calculations.^12^ The AMBER FF99SB forcefield for proteins,^13^ TIP3P water model,^14^ and the monovalent ion parameters of Joung and Cheatham^15^ were used to describe the rest of the system. The developed parameters for the His-Met ligated *c*-type heme are provided with the BioDC program in the accompanying Zenodo repository for this article.^16^

*Molecular Dynamics of Cytochrome c.* In a procedure very similar to that used previously for OmcS,^17^ solvated (oxidized or reduced) cytochrome *c* was subjected to 10000 steps of steepest descent followed by 40000 steps of conjugate-gradient minimization, all with a 10 kcal/(mol Å^2^) restraint on the heavy atoms of the protein backbone and the FE, NA, NB, NC, ND, C3D, C2A, C3B, C2C, CA, CB atoms of the heme group (using PDB atom nomenclature).

The system was subsequently heated in the *NVT* ensemble from 0 to 300 K at a rate of 0.3 K/ps and held at the final temperature for 5.0 ns. A 1.0 kcal/(mol Å^2^) restraint on the protein backbone and aforementioned heme atoms was applied during the heating process and first 1.0 ns at the final temperature, but then reduced to 0.1 kcal/(mol Å^2^) for the remaining 4.0 ns. After this thermalization process, the density of each system was equilibrated under 1.0-bar pressure for 4.0 ns in the *NPT* ensemble. Production stage simulations were conducted in the *NVT* ensemble at 300 K for 100 ns.

All *NVT* and *NPT* simulations (including the production-stage) employed periodic boundary conditions, the Particle Mesh Ewald^18^ treatment of electrostatic interactions with a direct sum cut-off of 10.0 Å, the SHAKE algorithm^19, 20^ to rigidify bonds to hydrogen atoms, a Langevin thermostat with a collision frequency of 2 ps^-1^, and an integration timestep for the Langevin equation of motion of 2.0 fs. Pressure in *NPT* simulations was regulated with a Monte Carlo barostat having a relaxation time of 1.0 ps. PMEMD in its CPU and GPU^21^ implementations of the Amber22 package^22^ were used to perform the minimization and dynamical simulations, respectively.

### Influence of Electronic polarization on Reorganization energy

The expression for quantifying polarization energy is given by Eq. S1.

$$U_{\mathrm{pol}}=-\frac{1}{2}\left( \Delta\alpha_{\mathrm{xx}}E^{2^{x}}+2\Delta\alpha_{\mathrm{xy}}E_{x}E_{y}+2\Delta\alpha_{\mathrm{xz}}E_{x}E_{z}+\Delta\alpha_{\mathrm{yy}}E_{y}^{2}+2\Delta\alpha_{\mathrm{zy}}E_{z}E_{z}+\Delta\alpha_{\mathrm{zz}}E_{z}^{2} \right) (S1)$$

Here, the multiplication of the components of the difference polarizability tensor $\left( \boldsymbol{\Delta\alpha} \right)$ with the electric field vector $\left( \mathbf{E} \right)$ are explicitly written out.

The sum of Eq. S1 evaluated for the donor and the acceptor gives the contribution of polarization energy to the electrostatic vertical energy gap $\left( \Delta U \right)$ in a given (reactant or product) state. The total vertical energy gap is $\Delta U=\Delta U_{\mathrm{coul}}+\Delta U_{\mathrm{pol}}$, where $\Delta U_{\mathrm{coul}}$ is the change in Coulombic interaction energy for altering the distribution of fixed-point charges on the donor and acceptor upon oxidation and reduction, respectively. The average and variance of $\Delta U$, in turn, enters the various definitions of the outer-sphere reorganization energy $\left( \lambda_{\mathrm{out}} \right)$ given in Eqs. 1 through 3 of the main text.

Given this background, the following ingredients are needed to assess the influence of electronic polarization of the active site on $\lambda_{\mathrm{out}}$: (1) The electric field vector on (i) the donor in the reactant state, (ii) the acceptor in the reactant state, (iii) the donor in the product state, and (iv) the acceptor in the product state, and (2) the difference polarizability tensor expressed as (i) $\left( \alpha^{\mathrm{ox}}-\alpha^{\mathrm{red}} \right)$ for the donor and (ii) $\left( \alpha^{\mathrm{red}}-\alpha^{\mathrm{ox}} \right)$ for the acceptor.

The electric field vectors were measured during the course of production-state non-polarizable classical molecular dynamics simulations using the TUPÃ python program^23^ with the following configuration file. Seven simulations were analyzed in which one heme was reduced and all other hemes were oxidized in a trimer model of the OmcS filament. The hemes that were separately reduced comprised the six hemes in the central subunit of the modeled oligomer and the first heme of the next subunit. In preparation for the electric field analysis with TUPÃ, each of these simulations was centered on the reduced (*i.e.*, electron donating heme). The obtained electronic field vectors on the Fe centers of the hemes are summarized in Table S10 and Figures S4 to S9.

**TUPÃ Configuration file:**

```
[Environment Selection]
sele_environment    = resid 1:1321 and not resid 1280

[Probe Selection]
# Provide the probe selection for the MODE of you choice
mode                = ATOM
selatom             = resname HEH and resid 1280 and name FE

[Solvent]
include_solvent     = True
solvent_cutoff      = 10
solvent_selection   = resname WAT Na+

[Time]
dt                  = 20
```

Note that “1280” is the residue ID of Heme #1 (residue name HEH) in the simulations.

Table S10. Average ± standard deviation for the components of the electric field vectors (in V/Å) on the Fe centers of the electron donating and accepting hemes in the reactant and product states for each electron transfer step through a unit cell of the OmcS filament. Note that there is some redundancy in the table because the simulation in which Heme #2 was reduced, for example, was considered both the produced state for the Heme #1 → 2 reaction and the reactant state for the Heme #2 → #3 reaction.

|  | Reactant State (V/A) | | | | | |
| --- | --- | --- | --- | --- | --- | --- |
|  | Donor | | | Acceptor | | |
|  | $E_{x}$ | $E_{y}$ | $E_{z}$ | $E_{x}$ | $E_{y}$ | $E_{z}$ |
| $1\to2$ | 0.267  ± 0.121 | -0.006  ± 0.155 | -0.112  ± 0.101 | 0.121  ± 0.117 | 0.000  ± 0.120 | 0.151  ± 0.118 |
| $2\to3$ | -0.037  ± 0.137 | -0.106  ± 0.124 | 0.105  ± 0.119 | -0.114  ± 0.081 | -0.118  ± 0.067 | -0.030  ± 0.049 |
| $3\to4$ | 0.050  ± 0.045 | -0.123  ± 0.044 | -0.093  ± 0.041 | 0.325  ± 0.090 | -0.062  ± 0.098 | -0.097  ± 0.098 |
| $4\to5$ | 0.117  ± 0.158 | -0.261  ± 0.093 | -0.184  ± 0.071 | 0.257  ± 0.118 | -0.189  ± 0.161 | -0.125  ± 0.093 |
| $5\to6$ | 0.261  ± 0.103 | -0.036  ± 0.112 | -0.225  ± 0.080 | 0.251  ± 0.117 | -0.090  ± 0.158 | 0.056  ± 0.097 |
| $6\to1^{'}$ | 0.177  ± 0.141 | 0.170  ± 0.131 | 0.085  ± 0.098 | -0.083  ± 0.149 | 0.158  ± 0.128 | 0.117  ± 0.096 |
|  | Product State (V/A) | | | | | |
|  | Donor | | | Acceptor | | |
|  | $E_{x}$ | $E_{y}$ | $E_{z}$ | $E_{x}$ | $E_{y}$ | $E_{z}$ |
| $1\leftarrow2$ | -0.037  ± 0.137 | -0.106  ± 0.124 | 0.105  ± 0.119 | 0.065  ± 0.125 | -0.108  ± 0.132 | -0.156  ± 0.105 |
| $2\leftarrow3$ | 0.050  ± 0.045 | -0.123  ± 0.044 | -0.093  ± 0.041 | 0.119  ± 0.112 | -0.060  ± 0.125 | 0.041  ± 0.117 |
| $3\leftarrow4$ | 0.117  ± 0.158 | -0.261  ± 0.093 | -0.184  ± 0.071 | -0.016  ± 0.045 | -0.077  ± 0.048 | -0.218  ± 0.037 |
| $4\leftarrow5$ | 0.261  ± 0.103 | -0.036  ± 0.112 | -0.225  ± 0.080 | 0.153  ± 0.114 | -0.096  ± 0.128 | -0.304  ± 0.085 |
| $5\leftarrow6$ | 0.177  ± 0.141 | 0.170  ± 0.131 | 0.085  ± 0.090 | 0.266  ± 0.112 | 0.189  ± 0.133 | -0.293  ± 0.072 |
| $6\leftarrow1^{'}$ | 0.125  ± 0.156 | 0.178  ± 0.116 | 0.135  ± 0.095 | 0.240  ± 0.120 | 0.064  ± 0.122 | -0.046  ± 0.092 |

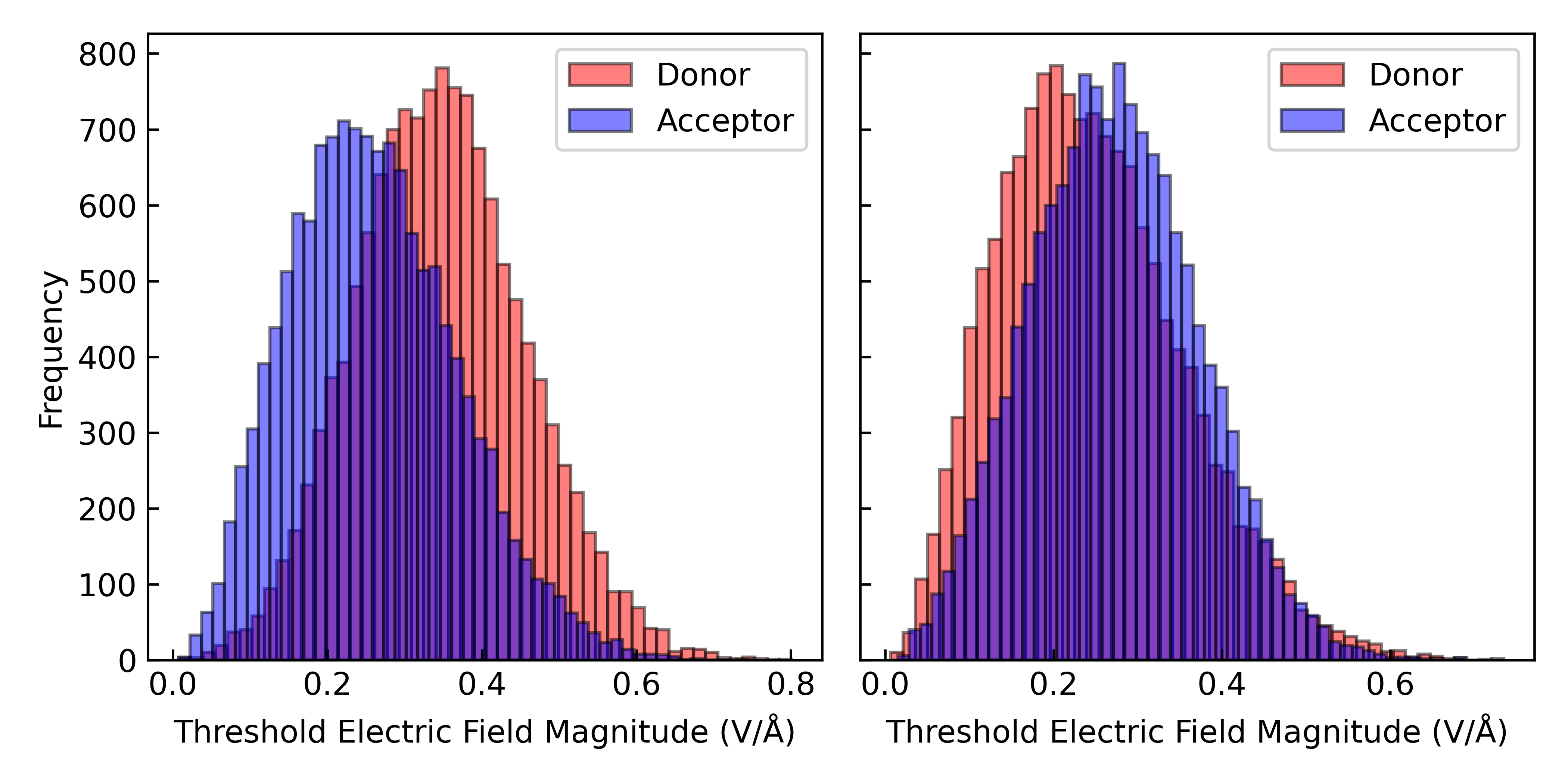

Figure S4. Distribution of electric field magnitudes on the Fe centers of the donor (Heme #1) and acceptor (Heme #2) in the reactant state (*left*) and the donor (Heme #2) and acceptor (Heme #1) in the product state (*right*).

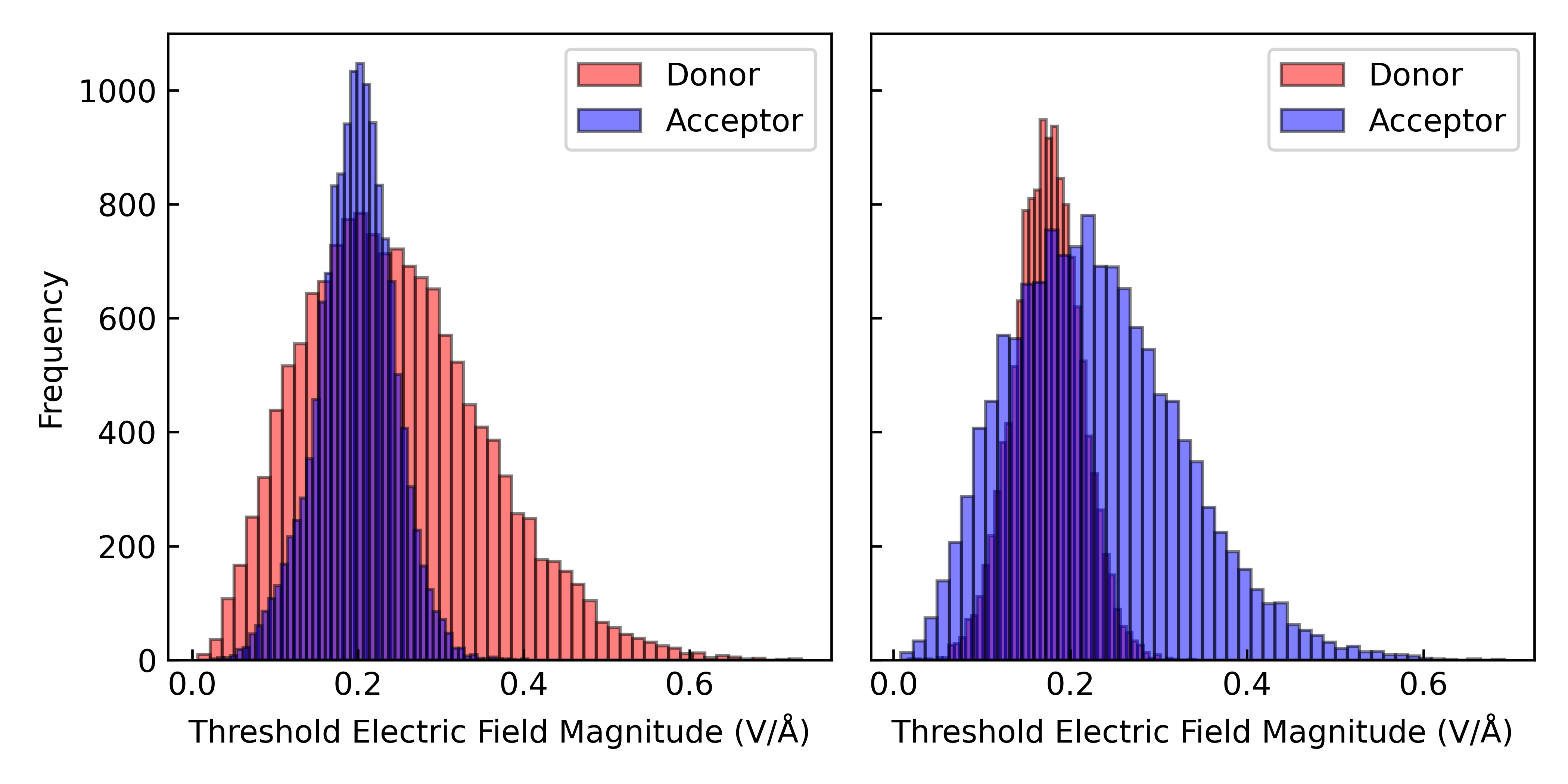

Figure S5. Distribution of electric field magnitudes on the Fe centers of the donor (Heme #2) and acceptor (Heme #3) in the reactant state (*left*) and the donor (Heme #3) and acceptor (Heme #2) in the product state (right).

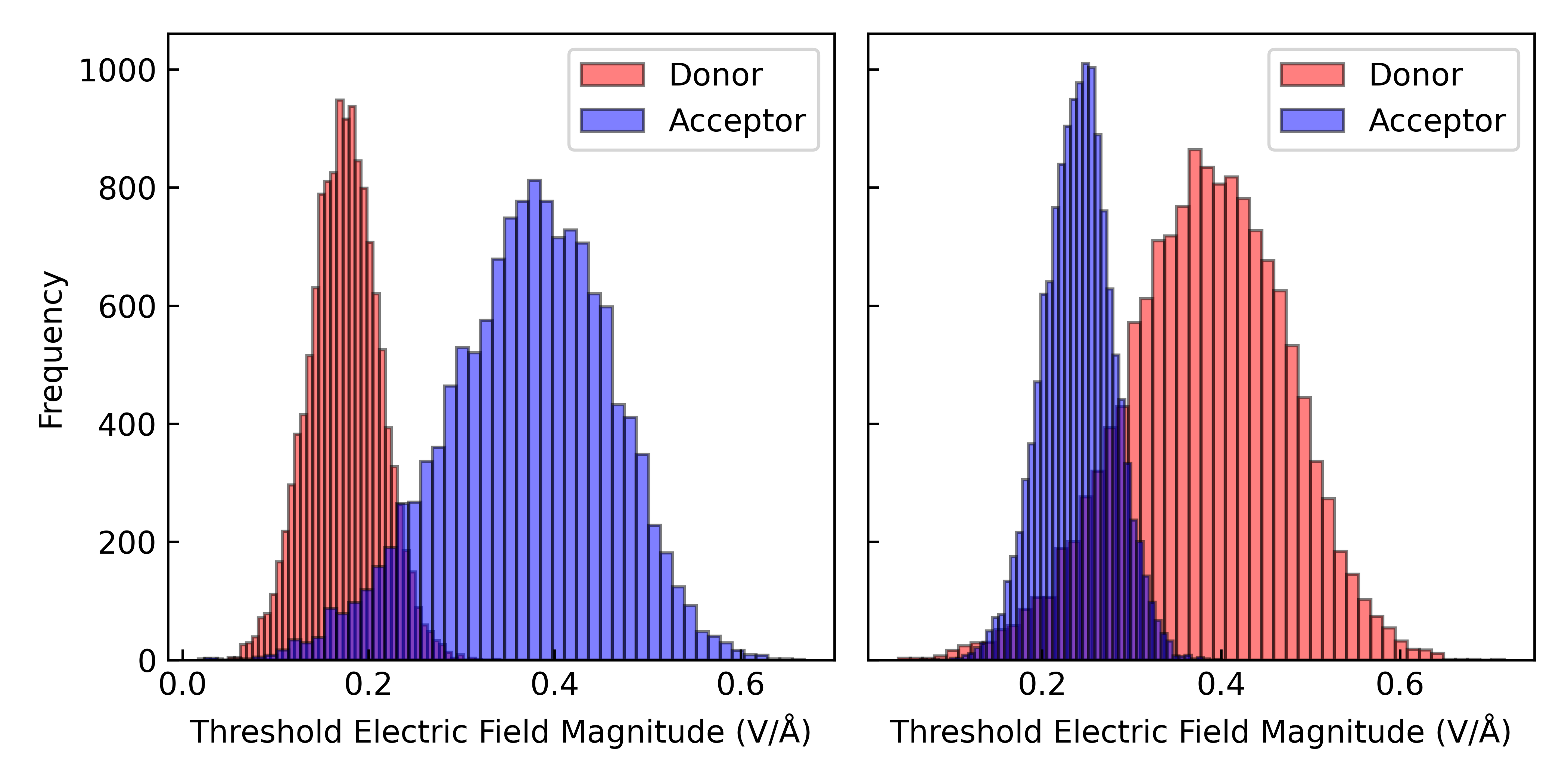

Figure S6. Distribution of electric field magnitudes on the Fe centers of the donor (Heme #3) and acceptor (Heme #4) in the reactant state (*left*) and the donor (Heme #4) and acceptor (Heme #3) in the product state (*right*).

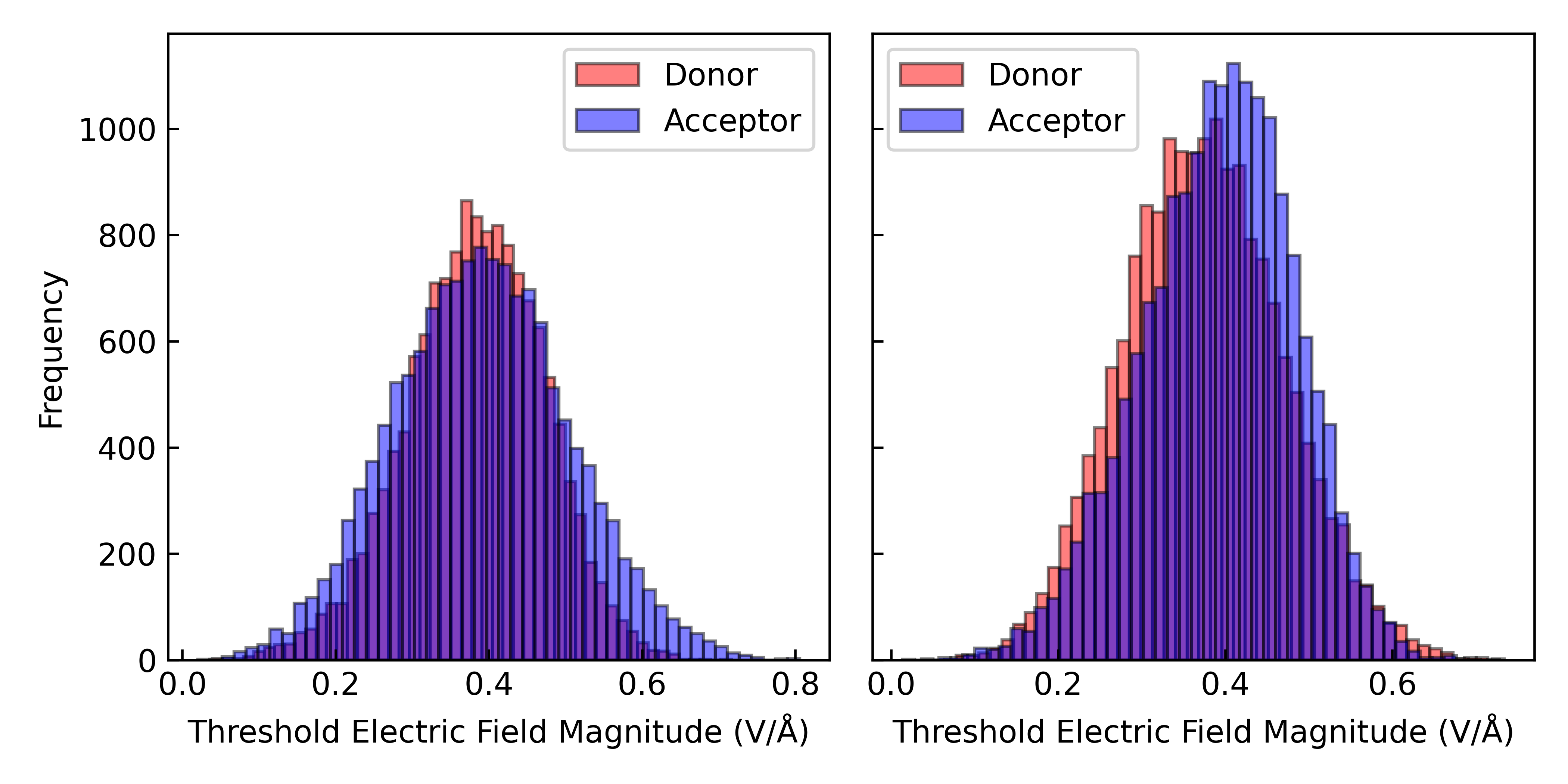

Figure S7. Distribution of electric field magnitudes on the Fe centers of the donor (Heme #4) and acceptor (Heme #5) in the reactant state (*left*) and the donor (Heme #5) and acceptor (Heme #4) in the product state (*right*).

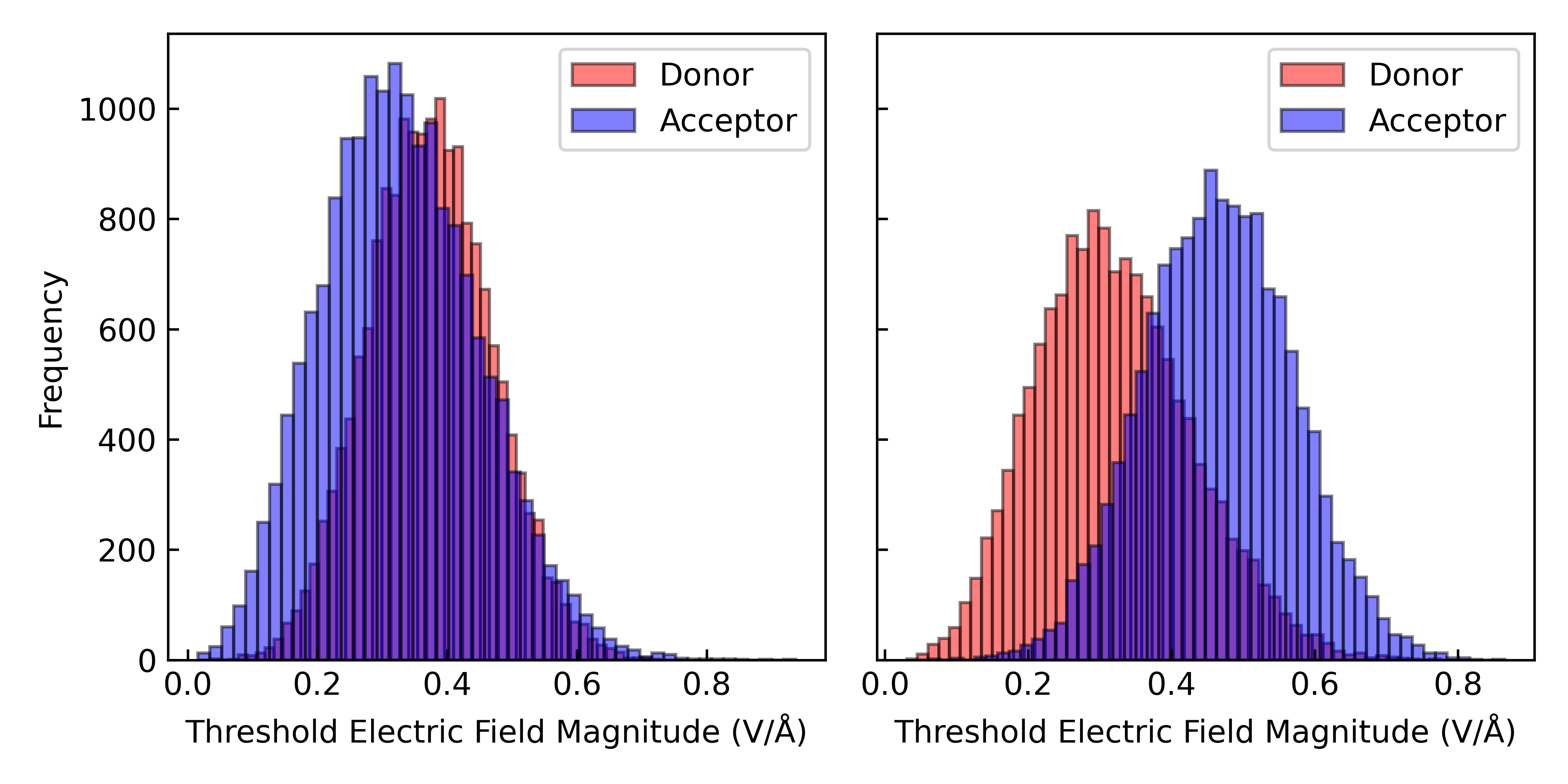

Figure S8. Distribution of electric field magnitudes on the Fe centers of the donor (Heme #5) and acceptor (Heme #6) in the reactant state (*left*) and the donor (Heme #6) and acceptor (Heme #5) in the product state (*right*).

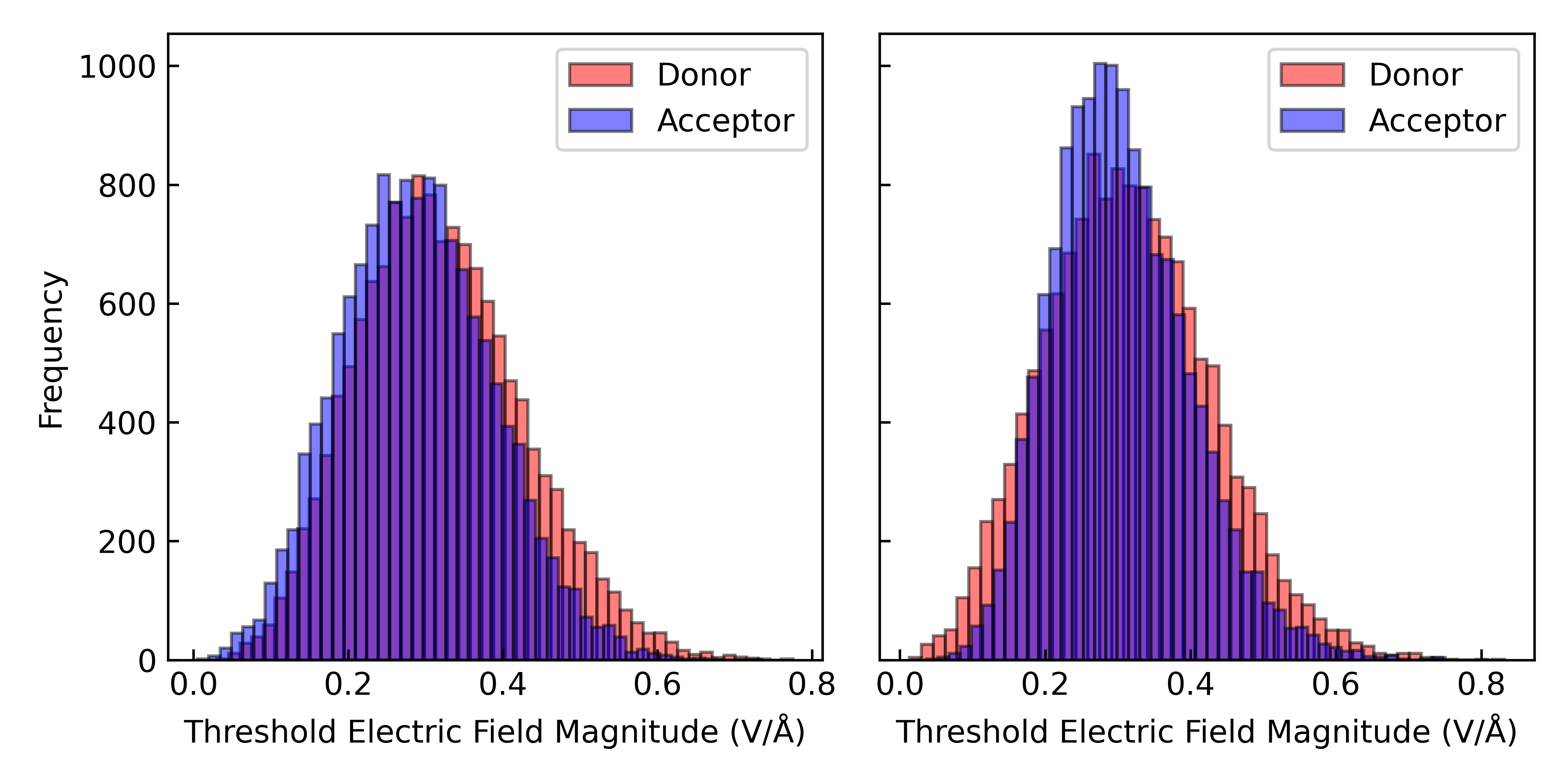

Figure S9. Distribution of electric field magnitudes on the Fe centers of the donor (Heme #6) and acceptor (Heme #1’) in the reactant state (*left*) and the donor (Heme #1’) and acceptor (Heme #6) in the product state (*right*).

Comparison of the polarizability tensor for the heme group computed in vacuum and embedded in a protein-water electrostatic environment (Tables S7 and S8 above) indicated little difference. The prior literature prescription^24, 25^ was therefore adopted of computing the polarizability tensor in vacuum in the standard orientation as defined by the Gaussian quantum chemical software package, and then rotating the tensor into the coordinate frame of the dynamical simulations for which the electric field analysis was performed.

The rotation matrix to be applied to the polarizability tensor at each frame of the dynamics was determined from a coordinate superimposition of the FE, NA, NB, NC, ND, CAB, and CAC atoms (PDB nomenclature) of the heme group in the quantum chemical calculation onto the same atoms of the donor or the acceptor (Figure S10). Figures S11 through S16 show that these coordinate superimpositions aligned the atoms with the donor and the acceptor in the reactant and product states with a root-mean-squared deviation of ~0.2 to 0.3 Å.

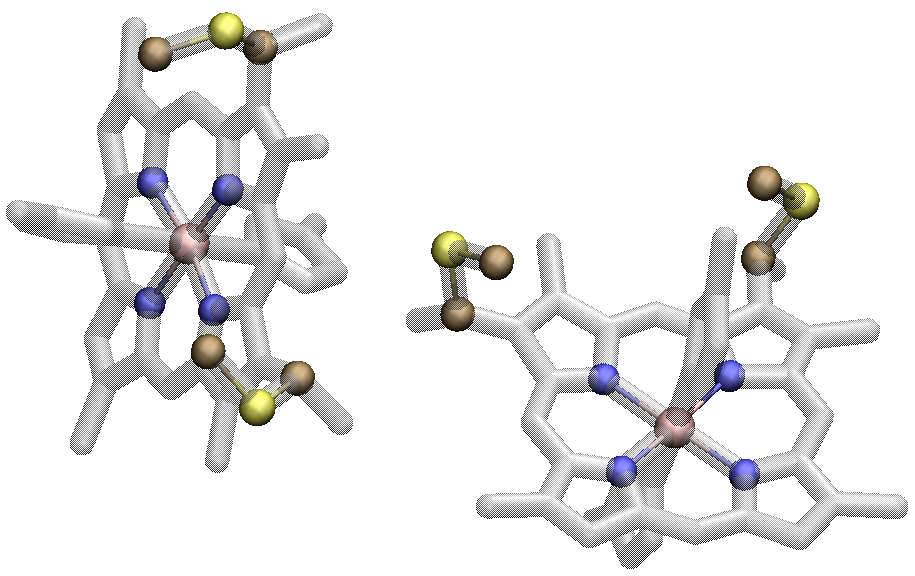

Figure S10 Depiction of the atoms (colored by element) used for coordinate superimposition of the quantum chemically optimized heme group in the reduced and oxidized states onto the donor and acceptor respectively at each frame of the dynamical simulations. The donor (Heme #1) and acceptor (Heme #2) in this case are shown from right to left.

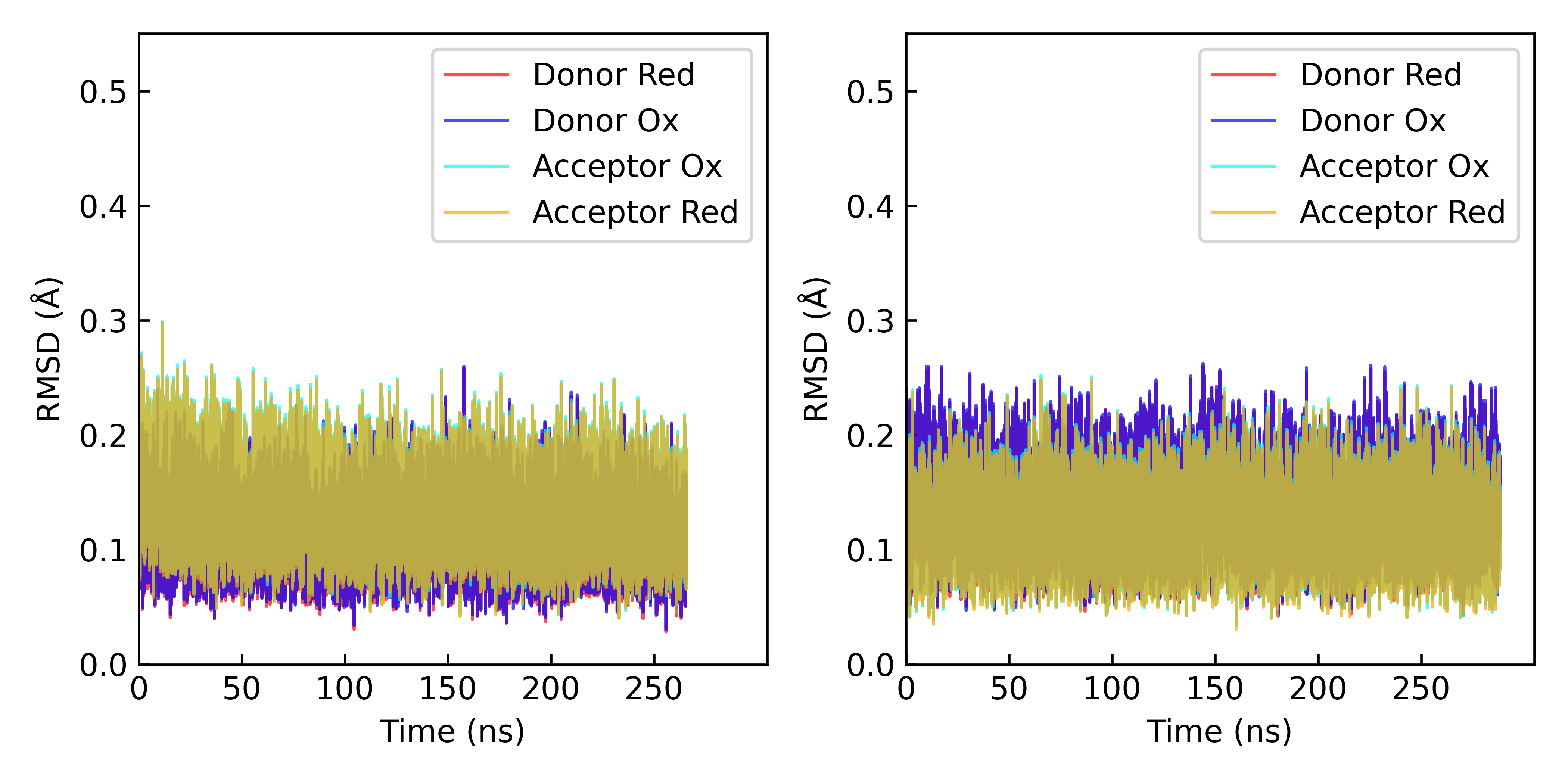

Figure S11. Root-mean-squared (RMSD) for superimposition of the FE, NA, NB, NC, ND , CAB, and CAC atoms of the quantum chemically optimized heme group in the reduced and oxidized states onto the donor and acceptor hemes in dynamical simulations of the reactant and product states. The reactant (*left*) and product (*right*) states pertain to the Heme #1 → #2 and #2 → #1 electron transfer reactions.

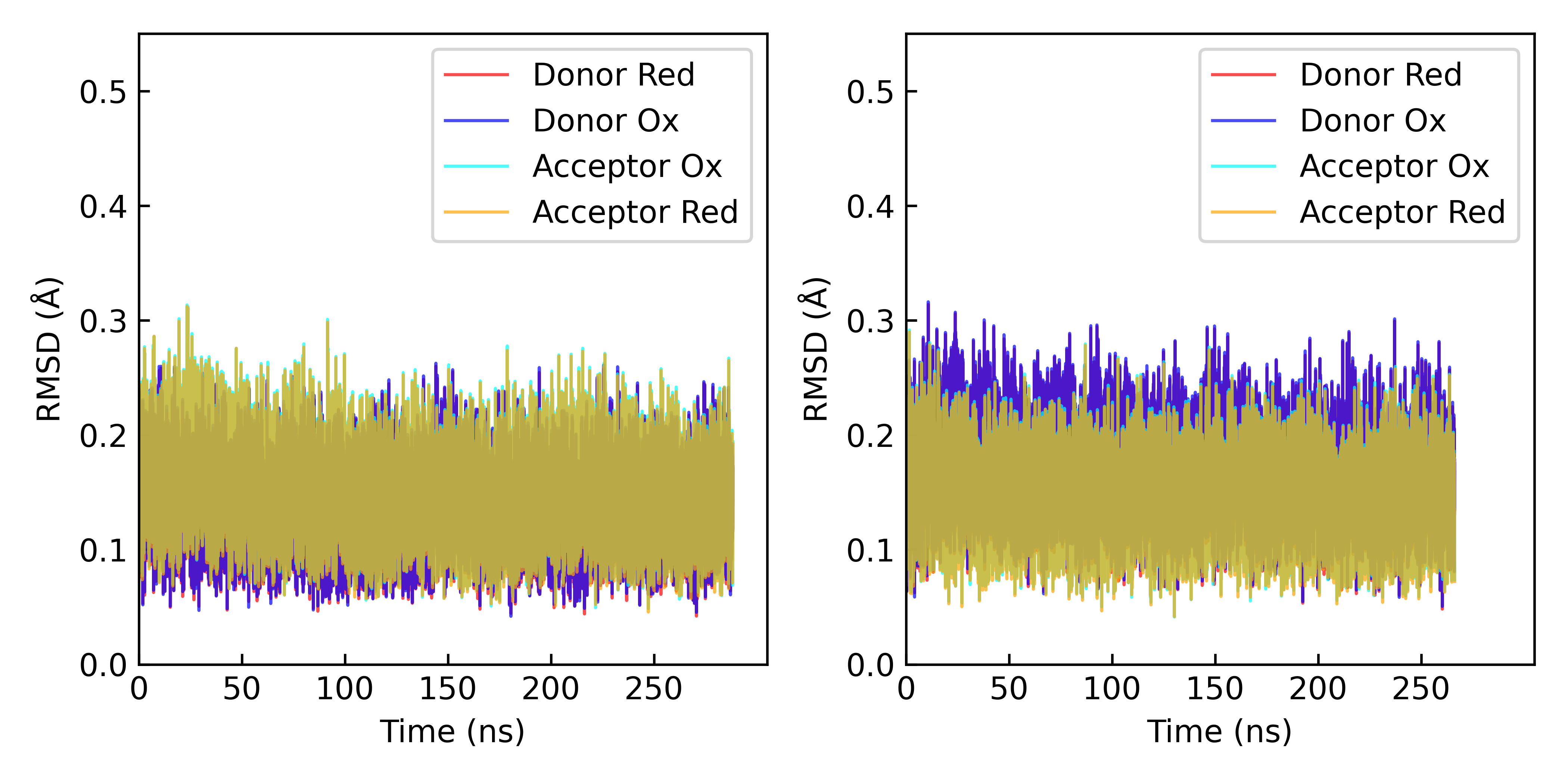

Figure S12. Root-mean-squared (RMSD) for superimposition of the FE, NA, NB, NC, ND , CAB, and CAC atoms of the quantum chemically optimized heme group in the reduced and oxidized states onto the donor and acceptor hemes in dynamical simulations of the reactant and product states. The reactant (*left*) and product (*right*) states pertain to the Heme #2 → #3 and #3 → #2 electron transfer reactions.

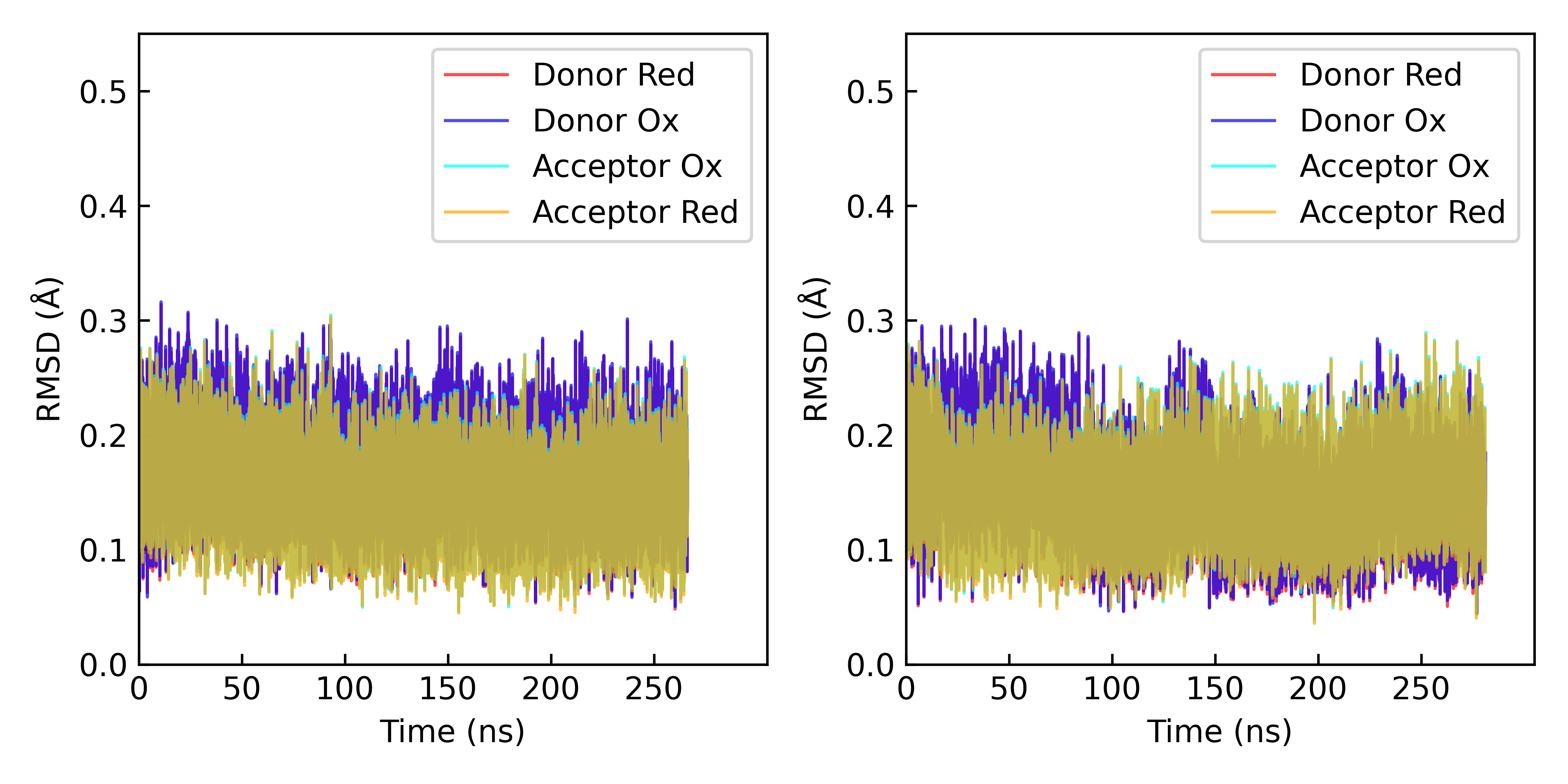

Figure S13. Root-mean-squared (RMSD) for superimposition of the FE, NA, NB, NC, ND , CAB, and CAC atoms of the quantum chemically optimized heme group in the reduced and oxidized states onto the donor and acceptor hemes in dynamical simulations of the reactant and product states. The reactant (*left*) and product (*right*) states pertain to the Heme #3 → #4 and #4 → #3 electron transfer reactions.

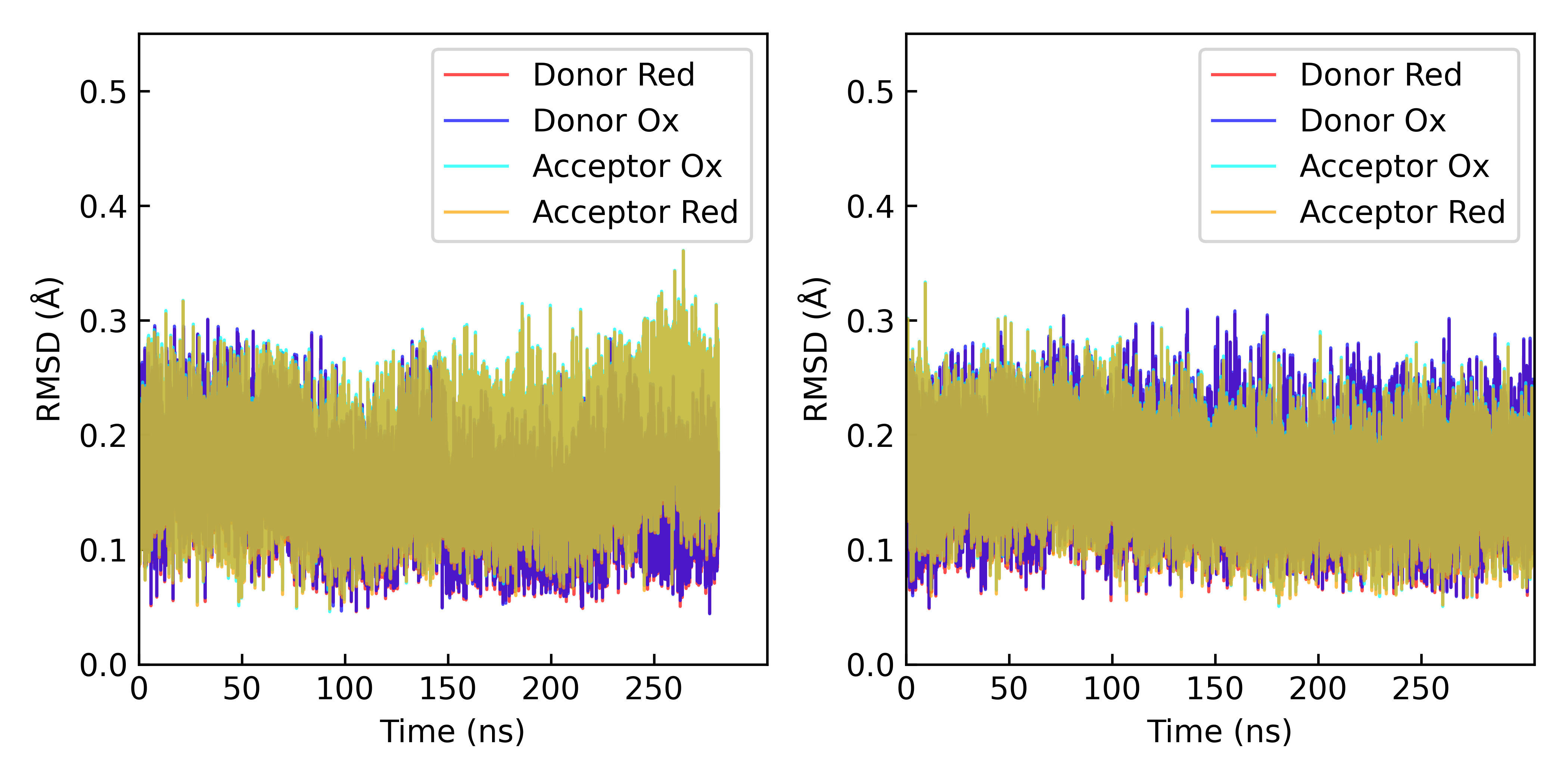

Figure S14. Root-mean-squared (RMSD) for superimposition of the FE, NA, NB, NC, ND , CAB, and CAC atoms of the quantum chemically optimized heme group in the reduced and oxidized states onto the donor and acceptor hemes in dynamical simulations of the reactant and product states. The reactant (*left*) and product (*right*) states pertain to the Heme #4 → #5 and #5 → #4 electron transfer reactions.

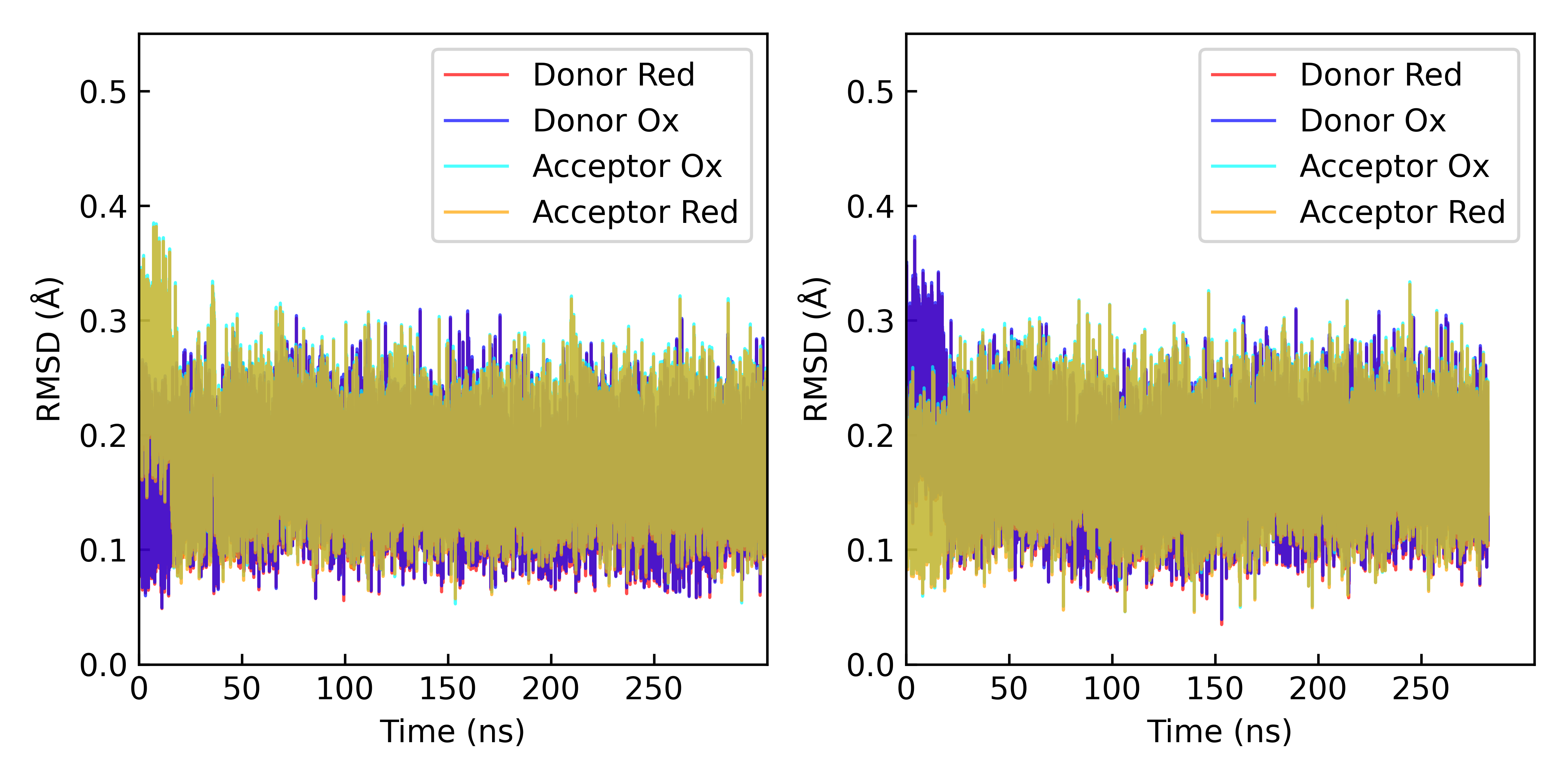

Figure S15. Root-mean-squared (RMSD) for superimposition of the FE, NA, NB, NC, ND , CAB, and CAC atoms of the quantum chemically optimized heme group in the reduced and oxidized states onto the donor and acceptor hemes in dynamical simulations of the reactant and product states. The reactant (*left*) and product (*right*) states pertain to the Heme #5 → #6 and #6 → #5 electron transfer reactions.

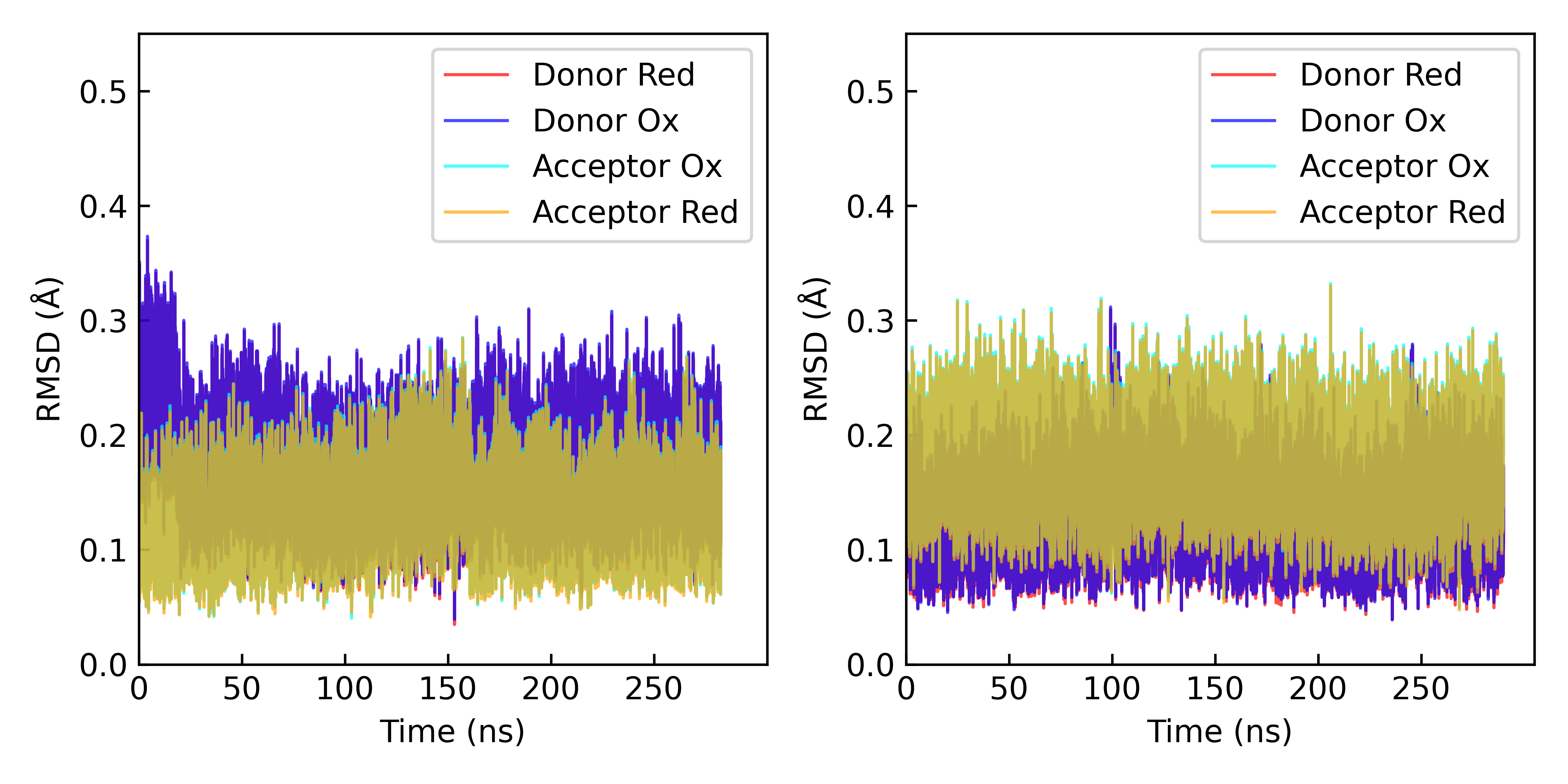

Figure S16. Root-mean-squared (RMSD) for superimposition of the FE, NA, NB, NC, ND , CAB, and CAC atoms of the quantum chemically optimized heme group in the reduced and oxidized states onto the donor and acceptor hemes in dynamical simulations of the reactant and product states. The reactant (*left*) and product (*right*) states pertain to the Heme #6 → #1’ and #1’ → #6 electron transfer reactions.

The rotation matrix at each frame of the dynamics was applied to the polarizability tensor in the reduced and oxidized states according to $\boldsymbol{\alpha}^{\mathbf{'}}\boldsymbol{=R\alpha}\mathbf{R}^{\mathbf{T}}$, where **R** is the rotation matrix and $\mathbf{R}^{\mathbf{T}}$ is its transpose. Subtraction of these rotated tensors gave the difference polarizability tensor needed in Eq. S1.

As a check on the implementation, $\boldsymbol{\alpha}$ was computed for acetaldehyde and then the molecule was rotated in 10° increments up to 90° from the original orientation. The same $\boldsymbol{\alpha}^{\mathbf{'}}$ was obtained by either (1) re-computing $\boldsymbol{\alpha}$ for the rotated molecule, or (2) rotating the originally computed $\boldsymbol{\alpha}$ by the rotation matrix determined from a coordinate superimposition.

With the appropriately rotated difference polarizability tensors and the electric field vectors in the same coordinate frame as the molecular dynamics, Eq. S1 was evaluated for the donor and the acceptor in the reactant state and the donor and acceptor in the product state. The obtained polarization energies were added to $\Delta U_{\mathrm{coul}}$, which was computed with the following input file to the CPPTRAJ program of AmberTools.^26^ The input file is annotated here for readability.

All energetic quantities are presented in Table S11. The time courses for the energies are visualized in Figures S17 to S22. The resulting reorganization energies are summarized in Table 3 of the main text. The dependence of the reorganization energy on the threshold magnitude of the field used to filter configurations form molecular dynamics is shown in Figures S23 to S27 for the electron transfer steps not shown in the main text.

```
#===============================================================
# Define the Boltzmann constant in eV/K and absolute temperature
kb= 0.00008617333262145
T= 300
#===============================================================

##Forward Reaction##############################################
# Load the reactant state topology and reactant state trajectory
parm   r1280/omcs_r1280.prmtop
trajin r1280/prod_1280centered.nc 1 last 1 parmindex 0

# Compute the electrostatic energy of the entire system in the
# reactant state
esander I1280 out I1280.dat cut 10.0 igb 0 ntb 1 ntf 2 ntc 2
run

# Compute the electrostatic self-energy of the reduced donor
strip !(:HEH&:1280) outprefix DAsubsysA1280
esander DADAsubsysA1280 out DAsubsysA1280.dat cut 10.0 igb 0 ntb 1 ntf 2 ntc 2
run

unstrip

# Compute the electrostatic self-energy of the oxidized acceptor
strip !(:HEH&:1274) outprefix DAsubsysA1274
esander DADAsubsysA1274 out DAsubsysA1274.dat cut 10.0 igb 0 ntb 1 ntf 2 ntc 2
run

unstrip

# Remove the self-energies from the total electrostatic energy
# to get a pair interaction energy
sysA = (I1280[elec] + I1280[elec14])
subAA = (DADAsubsysA1280[elec] + DADAsubsysA1280[elec14])
subAB = (DADAsubsysA1274[elec] + DADAsubsysA1274[elec14])
PairIntA = (sysA - subAA - subAB)
writedata If.dat sysA subAAb subAB PairIntA
run
#---------------------------------------------------------------

clear trajin

#---------------------------------------------------------------
# Load the reactant state trajectory with the product state
# topology
parm   r1274/omcs_r1274.prmtop
trajin r1280/prod_1280centered.nc 1 last 1 parmindex 1

# Compute the electrostatic energy of the entire system in the
# reactant state but with the product state topology
esander F1274 out F1274.dat cut 10.0 igb 0 ntb 1 ntf 2 ntc 2
run

# Compute the electrostatic self-energy of the donor in the
# reactant ensemble but with the product-state (oxidized) charge
# distribution
strip !(:HEH&:1280) outprefix DAsubsysB1280
esander DADAsubsysB1280 out DAsubsysB1280.dat cut 10.0 igb 0 ntb 1 ntf 2 ntc 2
run

unstrip

# Compute the electrostatic self-energy of the acceptor in the
# reactant state but with the product-state (reduced) charge
# distribution
strip !(:HEH&:1274) outprefix DAsubsysB1274
esander DADAsubsysB1274 out DAsubsysB1274.dat cut 10.0 igb 0 ntb 1 ntf 2 ntc 2
run

unstrip

# Remove the self-energies from the total electrostatic energy
# to get a pair interaction energy
sysB = (F1274[elec] + F1274[elec14])
subBA = (DADAsubsysB1280[elec] + DADAsubsysB1280[elec14])
subBB = (DADAsubsysB1274[elec] + DADAsubsysB1274[elec14])
PairIntB = (sysB - subBA - subBB)
writedata Ib.dat sysB subBA subBB PairIntB
run

# Compute the vertical energy gap for the forward reaction as
# the difference between the pair interaction energies; that is,
# (the energy of the oxidized donor and reduced acceptor) – (the
# energy of the reduced donor and oxidized acceptor), both
# interacting with the environment in the reactant ensemble.
VEGf = (PairIntB - PairIntA) * 0.0434
writedata VEGf.dat PairIntA PairIntB VEGf
#===============================================================

clear trajin

##Reverse Reaction##############################################
## Note that for consistency with the above input, the term
## “product state” is used. ####################################
#===============================================================
# Load the product state trajectory with the product state
# topology
trajin r1274/prod_1274centered.nc 1 last 1 parmindex 1

# Compute the electrostatic energy of the system in the product
# state
esander I1274 out I1274.dat cut 10.0 igb 0 ntb 1 ntf 2 ntc 2
run

# Compute the electrostatic self-energy of the oxidized acceptor
# in the product state
strip !(:HEH&:1280) outprefix DAsubsysC1280
esander DADAsubsysC1280 out DAsubsysC1280.dat cut 10.0 igb 0 ntb 1 ntf 2 ntc 2
run

unstrip

# Compute the electrostatic self-energy of the reduced donor in
# the product state
strip !(:HEH&:1274) outprefix DAsubsysC1274
esander DADAsubsysC1274 out DAsubsysC1274.dat cut 10.0 igb 0 ntb 1 ntf 2 ntc 2
run

unstrip

# Remove the self-energies from the electrostatic energy of the
# product state
sysC = (I1274[elec] + I1274[elec14])
subCA = (DADAsubsysC1280[elec] + DADAsubsysC1280[elec14])
subCB = (DADAsubsysC1274[elec] + DADAsubsysC1274[elec14])
PairIntC = (sysC - subCA - subCB)
writedata Ff.dat sysC subCA subCB PairIntC
run
#---------------------------------------------------------------

clear trajin

#---------------------------------------------------------------
# Load the product state trajectory with the reactant state
# topology
trajin r1274/prod_1274centered.nc 1 last 1 parmindex 0

# Compute the total electrostatic energy of the product state
# ensemble with the donor-acceptor pair having the reactant
# state charge distributions
esander F1280 out F1280.dat cut 10.0 igb 0 ntb 1 ntf 2 ntc 2
run

# Compute the electrostatic self-energy of the reduced acceptor
# in the product state ensemble
strip !(:HEH&:1280) outprefix DAsubsysD1280
esander DADAsubsysD1280 out DAsubsysD1280.dat cut 10.0 igb 0 ntb 1 ntf 2 ntc 2
run

unstrip

# Compute the electrostatic self-energy of the oxidized donor in
# the product state ensemble
strip !(:HEH&:1274) outprefix DAsubsysD1274
esander DADAsubsysD1274 out DAsubsysD1274.dat cut 10.0 igb 0 ntb 1 ntf 2 ntc 2
run

unstrip

# Remove the donor-acceptor self-energies from the electrostatic
# energy of the product state with the donor-acceptor pair
# having the reactant state charge distribution
sysD = (F1280[elec] + F1280[elec14])
subDA = (DADAsubsysD1280[elec] + DADAsubsysD1280[elec14])
subDB = (DADAsubsysD1274[elec] + DADAsubsysD1274[elec14])
PairIntD = (sysD - subDA - subDB)
writedata Fb.dat sysD subDA subDB PairIntD
run

# Compute the vertical energy gap for the backward reaction
VEGb = (PairIntC - PairIntD) * 0.0434
writedata VEGb.dat PairIntC PairIntD VEGb
#===============================================================

# Compute the statistical moments of the vertical energy gaps in
# the forward and backward directions and the associated
# reorganization energies
avgVEGf   = avg(VEGf)
avgVEGb   = avg(VEGb)
lambdast  = (avgVEGf - avgVEGb)/2

varVEGf   = stdev(VEGf)^2
varVEGb   = stdev(VEGb)^2
lambdavarf = varVEGf/(2*kb*T)
lambdavarb = varVEGb/(2*kb*T)

xg = (lambdavarf + lambdavarb)/(2*lambdast)

printdata avgVEGf avgVEGb varVEGf varVEGb
printdata lambdast lambdavarf lambdavarb xg
writedata Reorg1280,1274.dat lambdast lambdavarf lambdavarb xg
run

quit
```

Table 11. Coulombic, polarization contributions and total electrostatic vertical energy gaps for each forward $\left( \to\right)$ and backward $\left( \leftarrow\right)$ electron transfer step in OmcS. The average ± variance is shown for each quantity, where the statistics are taken over n frames. The length of the analyzed production-stage simulations is given by n frames multiplied by the saving frequency of 0.02 ns/frame.

|  | n | $\Delta U_{\mathrm{Coul}}$ | ${\Delta U}_{\mathrm{pol}}^{D}$ | ${\Delta U}_{\mathrm{pol}}^{A}$ | ${\Delta U}_{\mathrm{pol}}^{D+A}$ | $\Delta U$ |
| --- | --- | --- | --- | --- | --- | --- |
| $1\to2$ | 13296 | 0.698±0.105 | -0.333±0.031 | 0.086±0.139 | -0.247±0.047 | 0.452±0.197 |
| $1\leftarrow2$ | 14421 | -0.927±0.050 | -0.228±0.054 | 0.279±0.058 | 0.051±0.109 | -0.876±0.166 |
| $2\to3$ | 14421 | 0.504±0.026 | -0.228±0.054 | 0.158±0.080 | -0.069±0.060 | 0.435±0.088 |
| $2\leftarrow3$ | 13314 | -0.436±0.027 | -0.160±0.005 | 0.163±0.044 | 0.003±0.045 | -0.434±0.061 |
| $3\to4$ | 13314 | 0.722±0.029 | -0.160±0.005 | 0.538±0.066 | 0.378±0.072 | 1.100±0.110 |
| $3\leftarrow4$ | 14065 | -0.410±0.027 | -0.473±0.211 | 0.092±0.012 | -0.381±0.156 | -0.791±0.183 |
| $4\to5$ | 14065 | 0.583±0.034 | -0.473±0.211 | 0.394±0.077 | -0.080±0.358 | 0.503±0.457 |
| $4\leftarrow5$ | 17479 | -0.741±0.031 | -0.403±0.040 | 0.457±0.098 | 0.054±0.135 | -0.687±0.153 |
| $5\to6$ | 17479 | 0.653±0.040 | -0.403±0.040 | 0.246±0.055 | -0.156±0.098 | 0.497±0.137 |
| $5\leftarrow6$ | 14127 | -0.638±0.038 | -0.184±0.048 | 0.722±0.109 | 0.538±0.164 | -0.101±0.171 |
| $6\to1$’ | 14127 | 0.645±0.036 | -0.184±0.048 | 0.294±0.073 | 0.110±0.127 | 0.755±0.130 |
| $6\leftarrow1^{'}$ | 14501 | -0.495±0.033 | -0.342±0.104 | 0.406±0.083 | 0.064±0.195 | -0.430±0.238 |

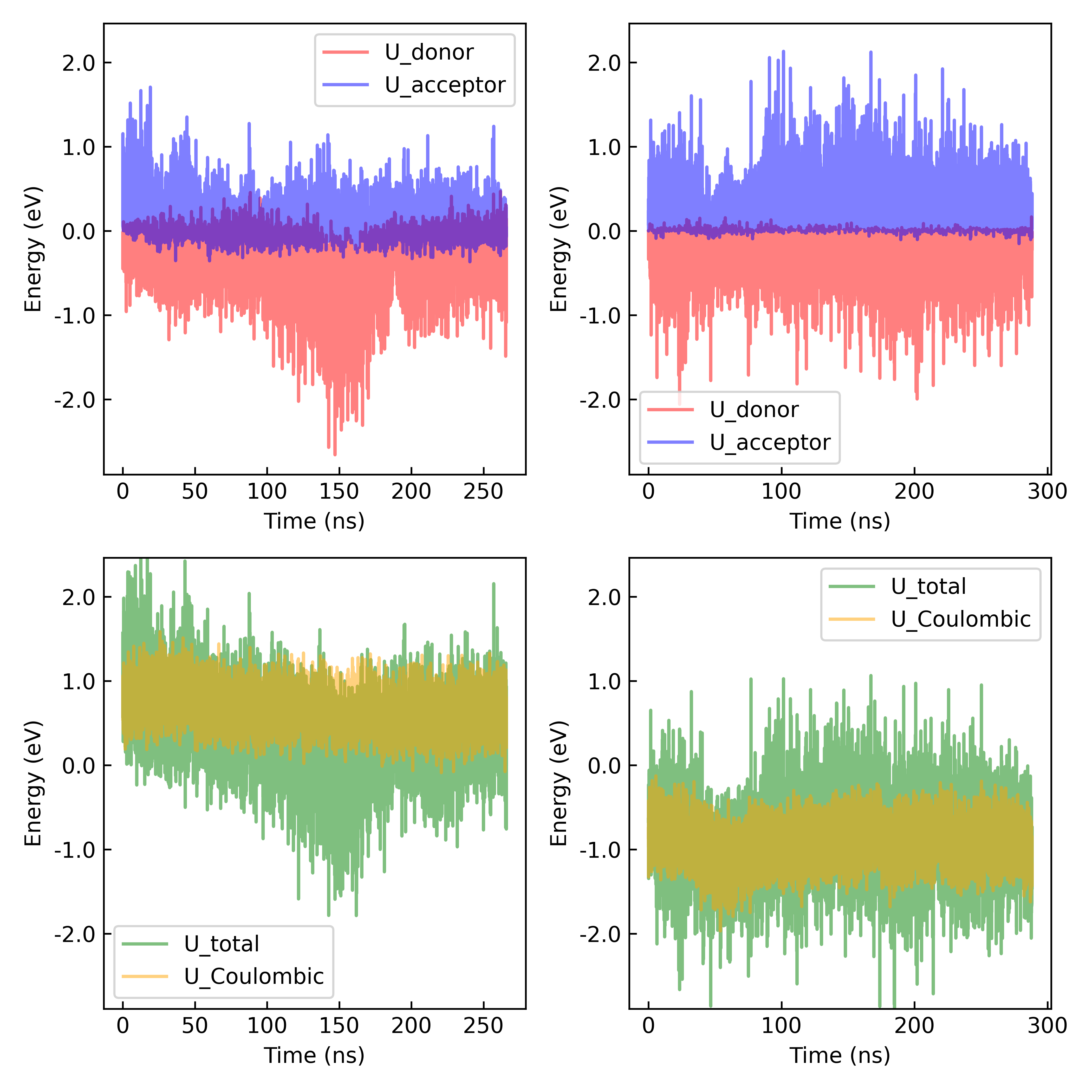

Figure S17. Time series for the polarization energies of the donor and acceptor, as well as the Coulombic energy and the total (Coulomb + donor and acceptor polarization energies) vertical energy gap computed in the reactant (*left column*) and product (*right column*) states for the reaction Heme #1^red^ + Heme #2^ox^ → Heme #1^ox^ + Heme #2^red^.

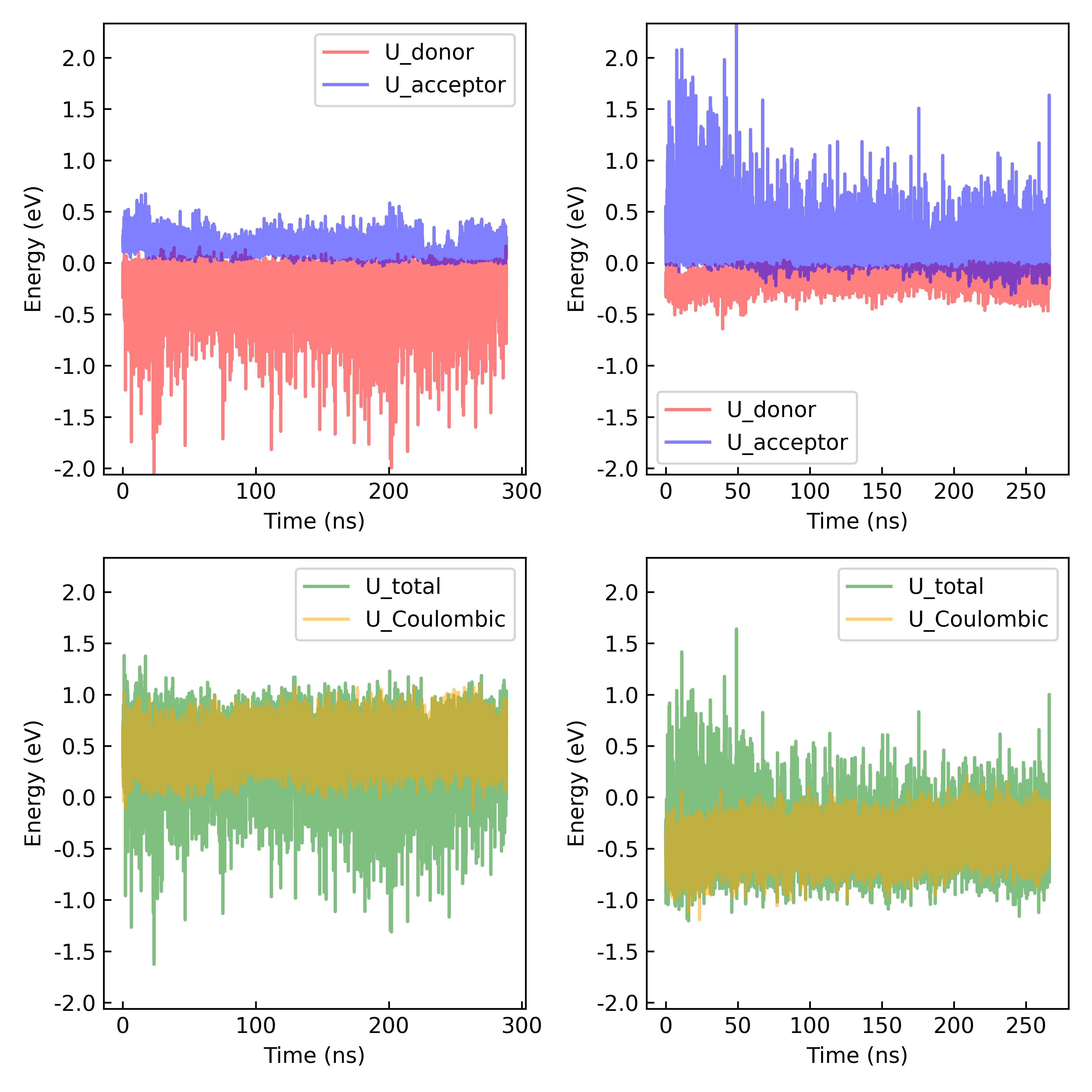

Figure S18. Time series for the polarization energies of the donor and acceptor, as well as the Coulombic energy and the total (Coulomb + donor and acceptor polarization energies) vertical energy gap computed in the reactant (*left column*) and product (*right column*) states for the reaction Heme #2^red^ + Heme #3^ox^ → Heme #2^ox^ + Heme #3^red^.

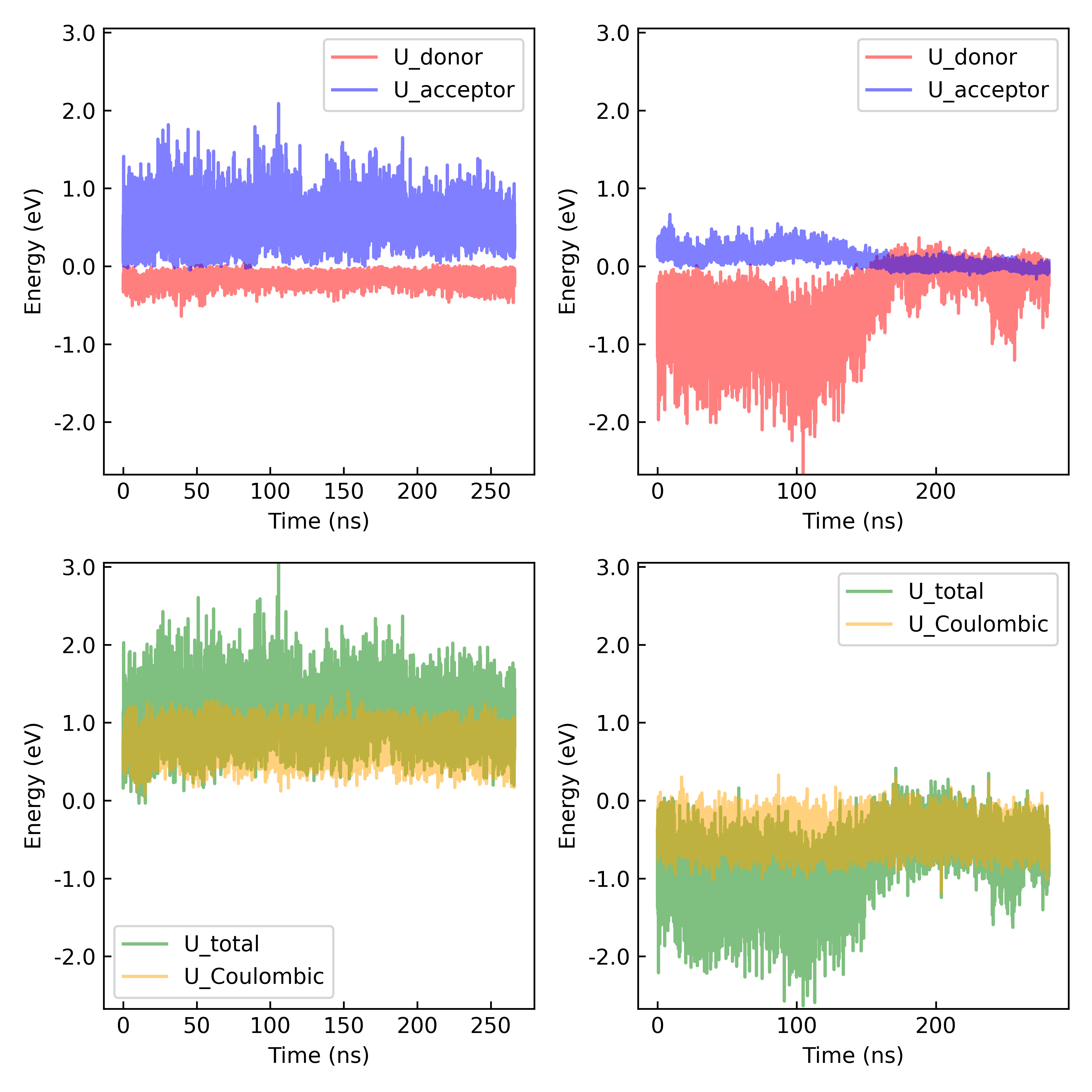

Figure S19. Time series for the polarization energies of the donor and acceptor, as well as the Coulombic energy and the total (Coulomb + donor and acceptor polarization energies) vertical energy gap computed in the reactant (*left column*) and product (*right column*) states for the reaction Heme #3^red^ + Heme #4^ox^ → Heme #3^ox^ + Heme #4^red^.

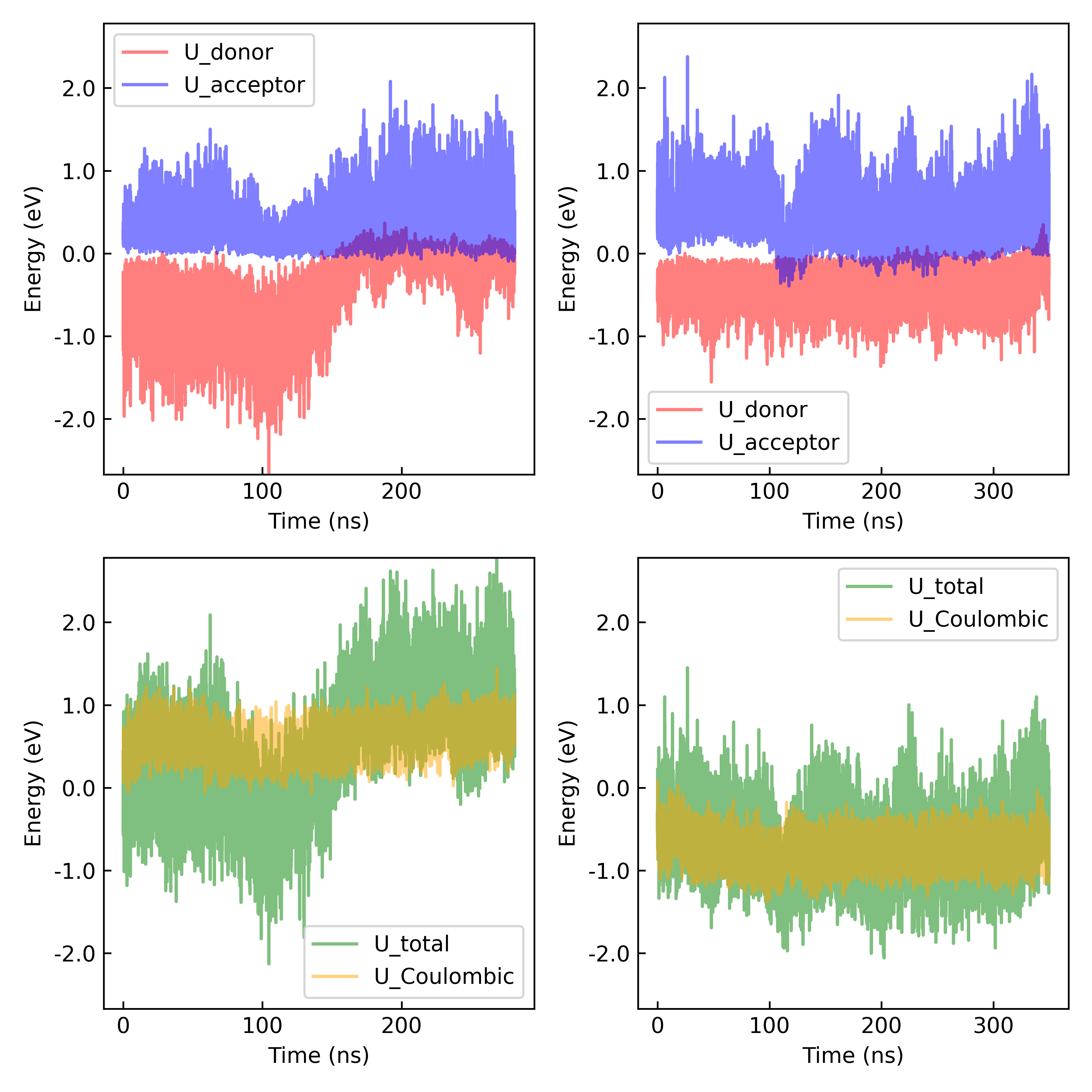

Figure S20. Time series for the polarization energies of the donor and acceptor, as well as the Coulombic energy and the total (Coulomb + donor and acceptor polarization energies) vertical energy gap computed in the reactant (*left column*) and product (*right column*) states for the reaction Heme #4^red^ + Heme #5^ox^ → Heme #4^ox^ + Heme #5^red^.

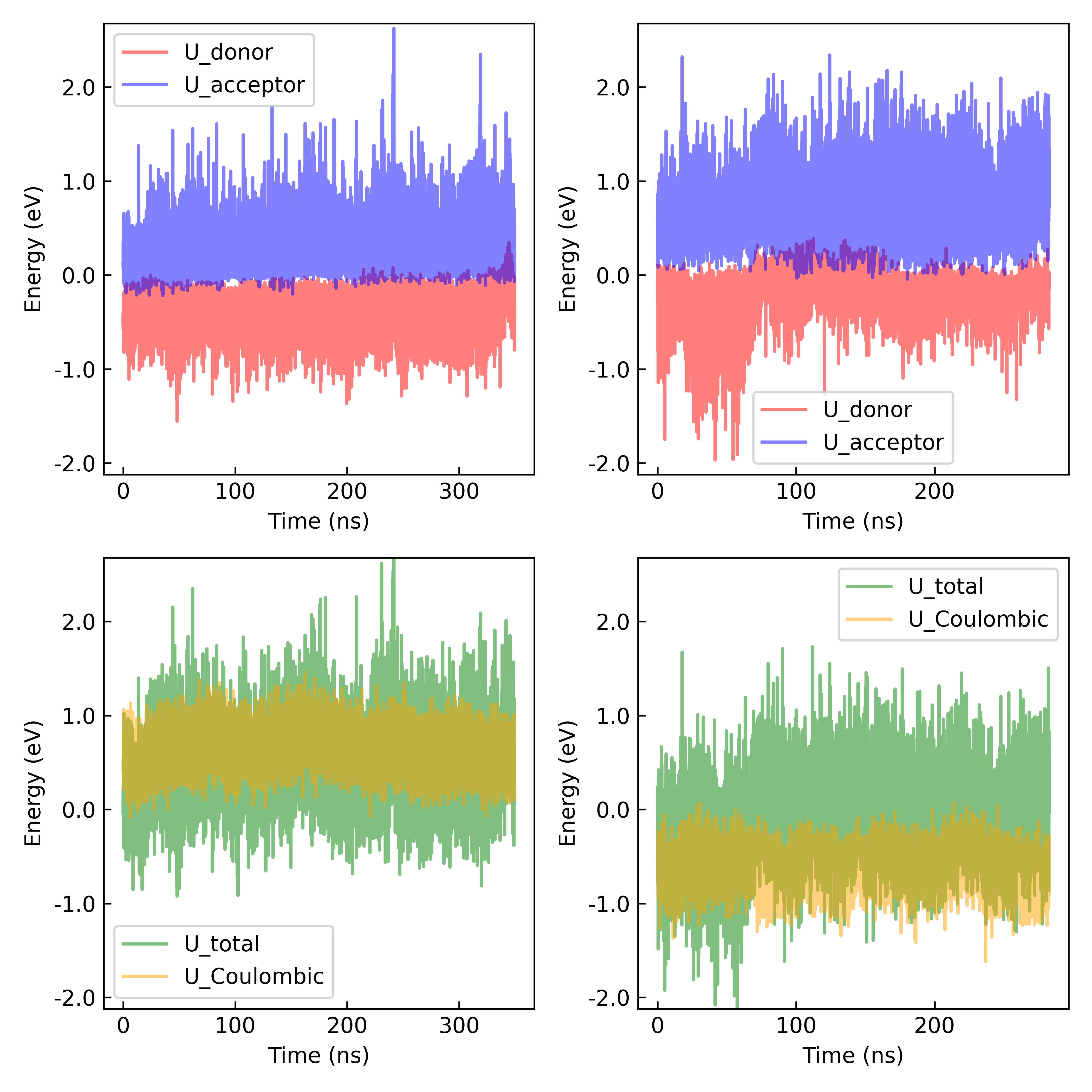

Figure S21. Time series for the polarization energies of the donor and acceptor, as well as the Coulombic energy and the total (Coulomb + donor and acceptor polarization energies) vertical energy gap computed in the reactant (*left column*) and product (*right column*) states for the reaction Heme #5^red^ + Heme #6^ox^ → Heme #5^ox^ + Heme #6^red^.

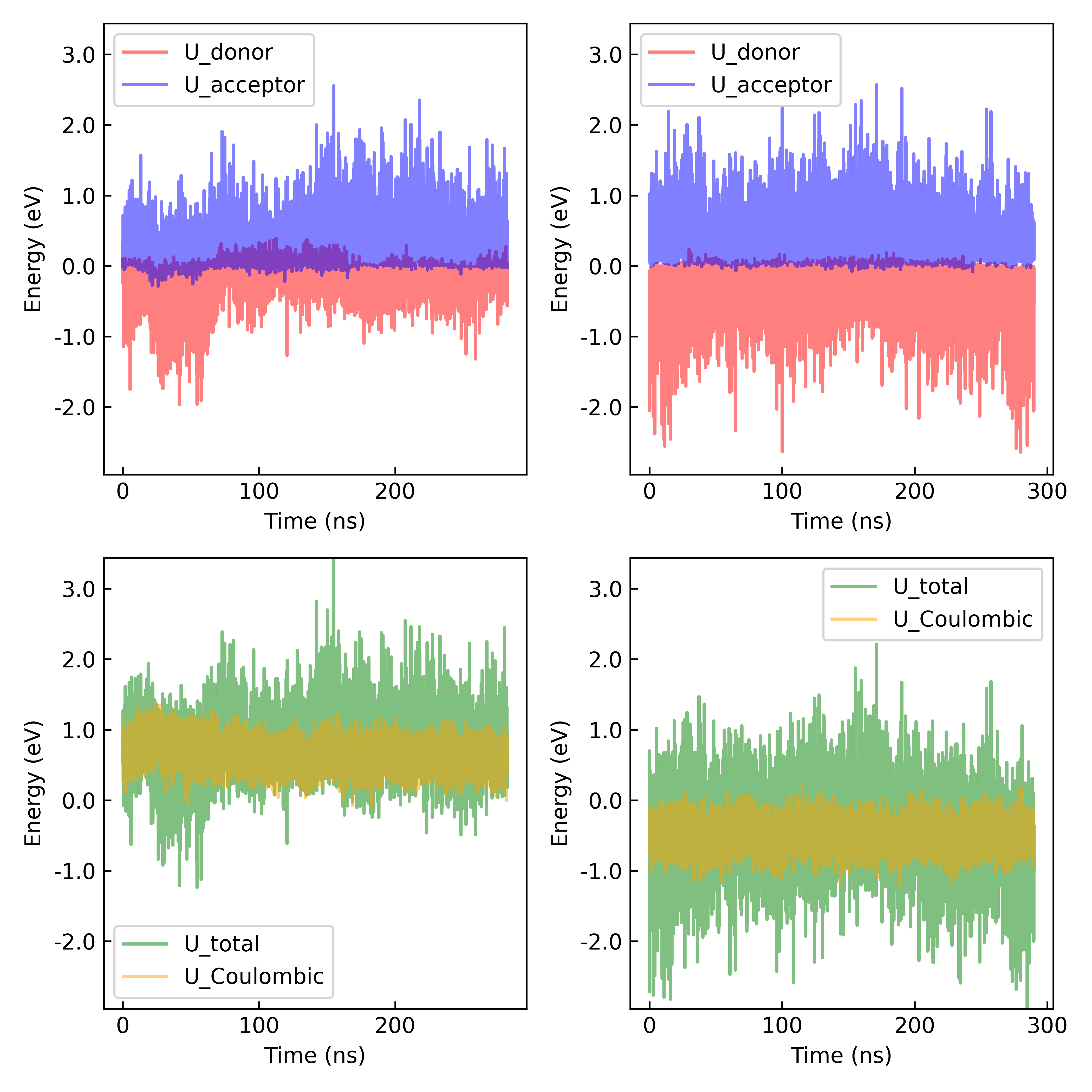

Figure S22. Time series for the polarization energies of the donor and acceptor, as well as the Coulombic energy and the total (Coulomb + donor and acceptor polarization energies) vertical energy gap computed in the reactant (*left column*) and product (*right column*) states for the reaction Heme #6^red^ + Heme #1’^ox^ → Heme #6^ox^ + Heme #1’^red^.

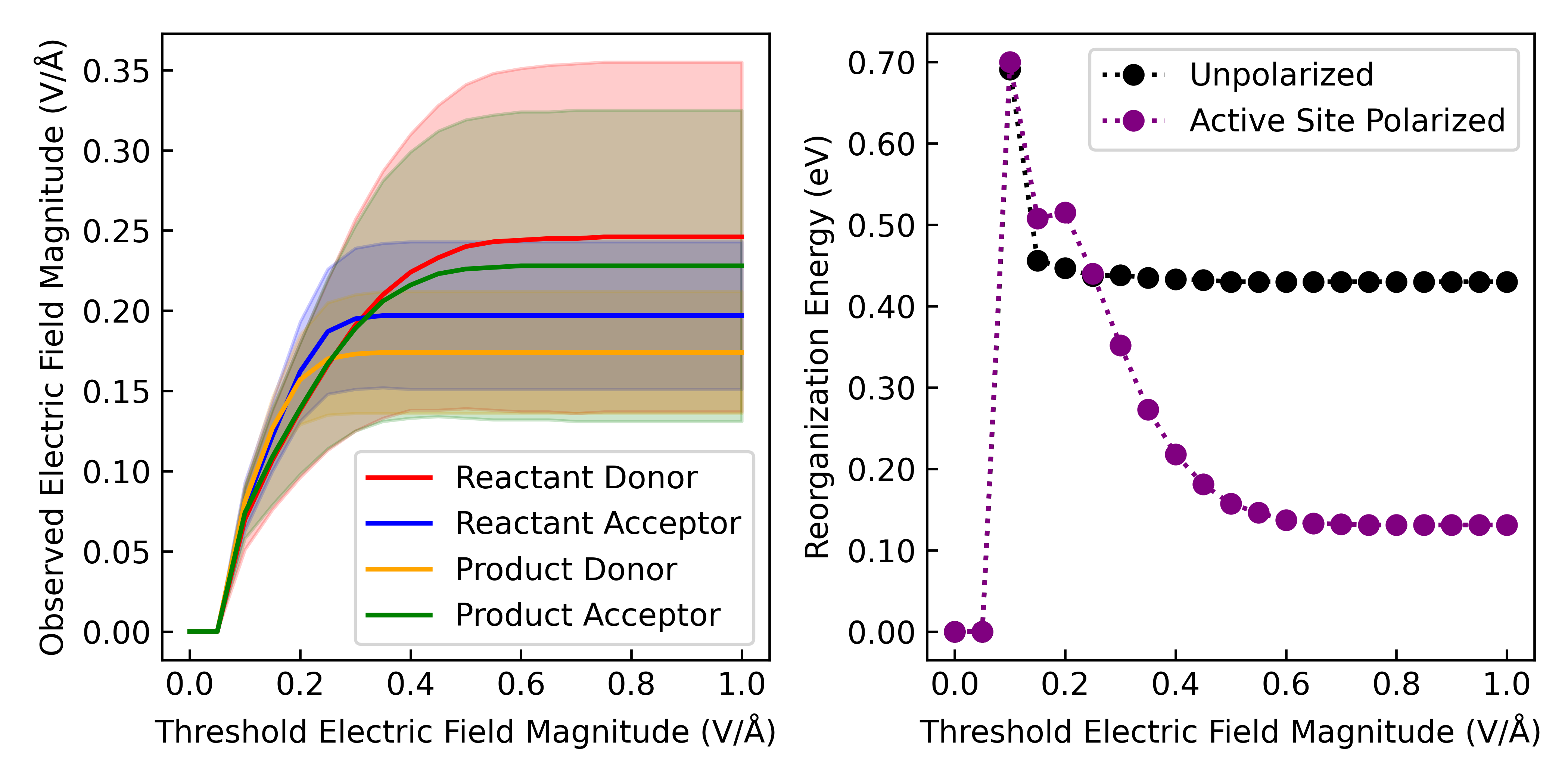

Figure S23. A wide distribution of electric field magnitudes exerted on the Fe centers of the heme groups in OmcS induces a significance decrease in the outer-sphere reorganization energy. The average magnitude and standard deviation of the field on the donor and acceptor in both states (*left*) and the reorganization energy (*right*) are plotted as a function of the threshold field magnitude used to filter the configurations from molecular dynamics. The data is shown for electron transfer between Hemes #2 and #3.

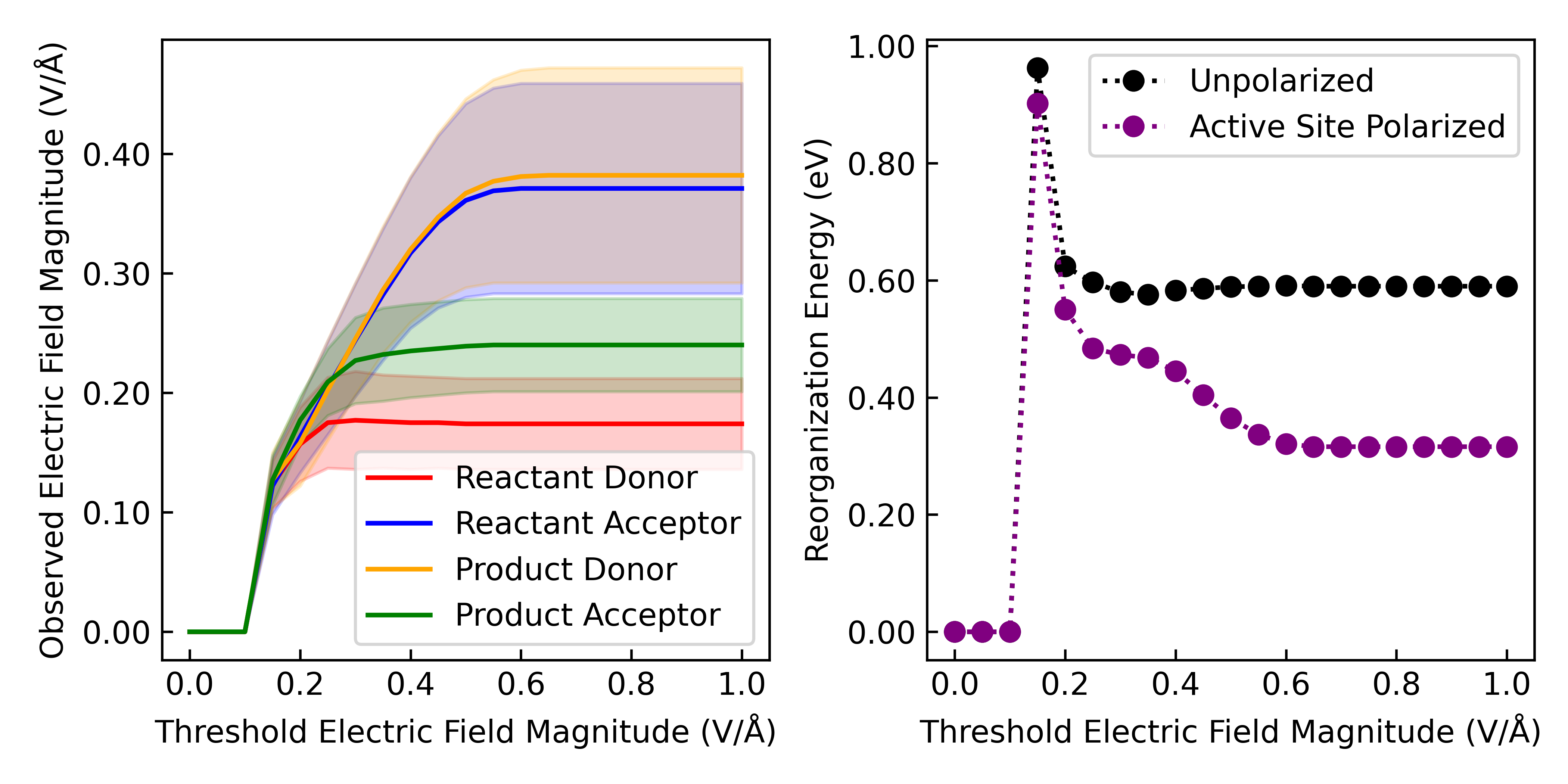

Figure S24. A wide distribution of electric field magnitudes exerted on the Fe centers of the heme groups in OmcS induces a significance decrease in the outer-sphere reorganization energy. The average magnitude and standard deviation of the field on the donor and acceptor in both states (*left*) and the reorganization energy (*right*) are plotted as a function of the threshold field magnitude used to filter the configurations from molecular dynamics. The data is shown for electron transfer between Hemes #3 and #4.

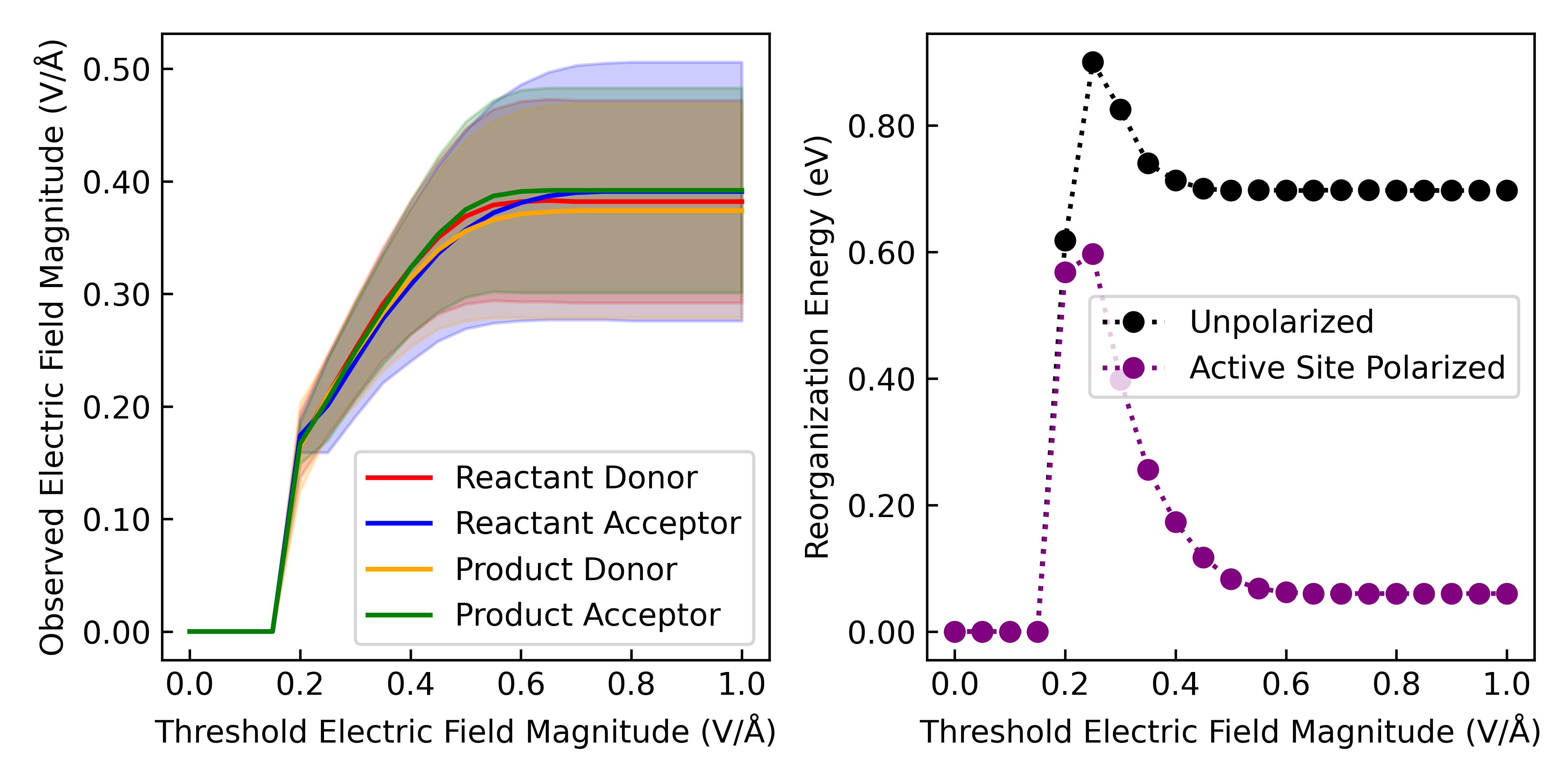

Figure S25. A wide distribution of electric field magnitudes exerted on the Fe centers of the heme groups in OmcS induces a significance decrease in the outer-sphere reorganization energy. The average magnitude and standard deviation of the field on the donor and acceptor in both states (*left*) and the reorganization energy (*right*) are plotted as a function of the threshold field magnitude used to filter the configurations from molecular dynamics. The data is shown for electron transfer between Hemes #4 and #5.

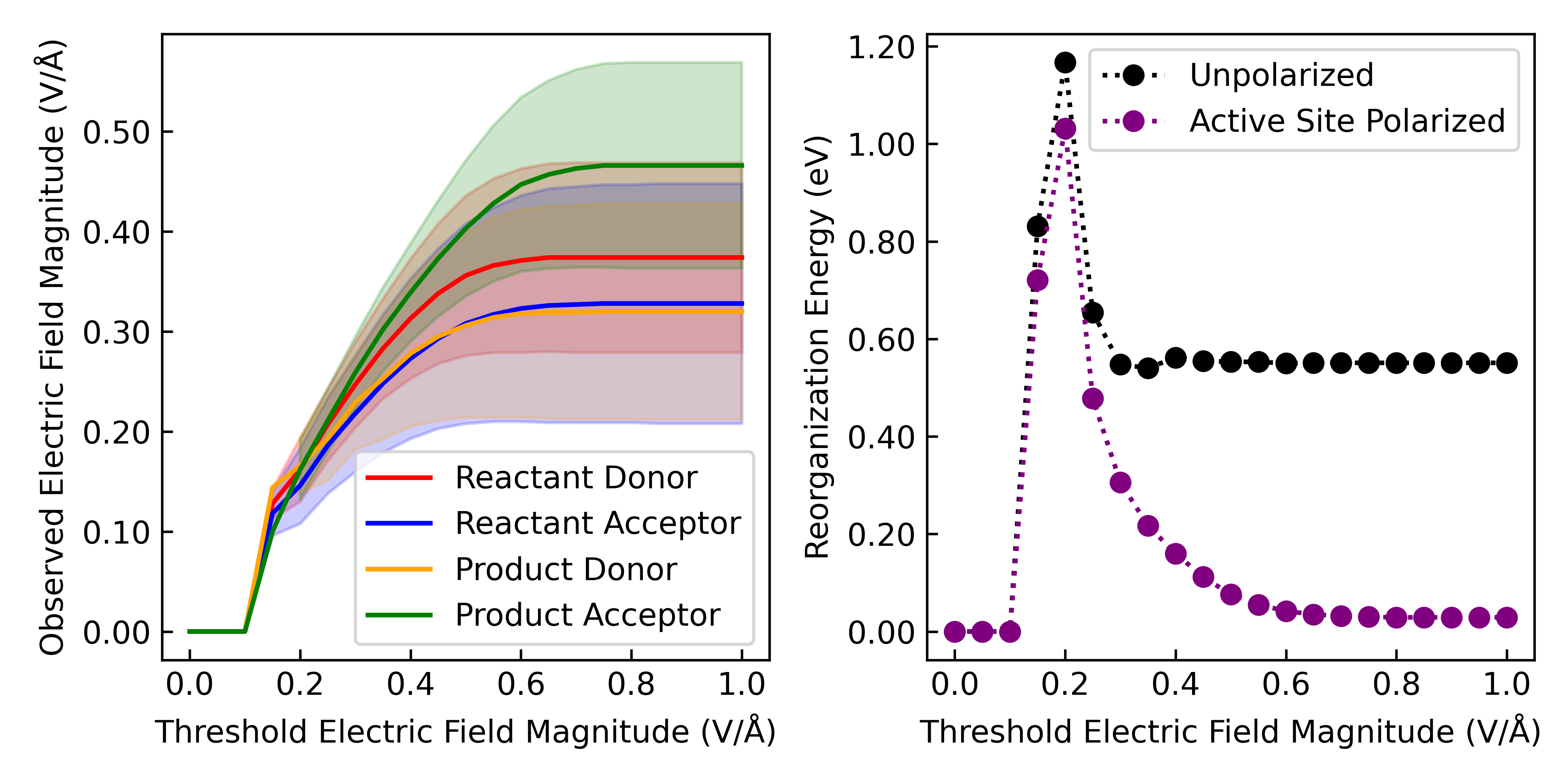

Figure S26. A wide distribution of electric field magnitudes exerted on the Fe centers of the heme groups in OmcS induces a significance decrease in the outer-sphere reorganization energy. The average magnitude and standard deviation of the field on the donor and acceptor in both states (*left*) and the reorganization energy (*right*) are plotted as a function of the threshold field magnitude used to filter the configurations from molecular dynamics. The data is shown for electron transfer between Hemes #5 and #6.

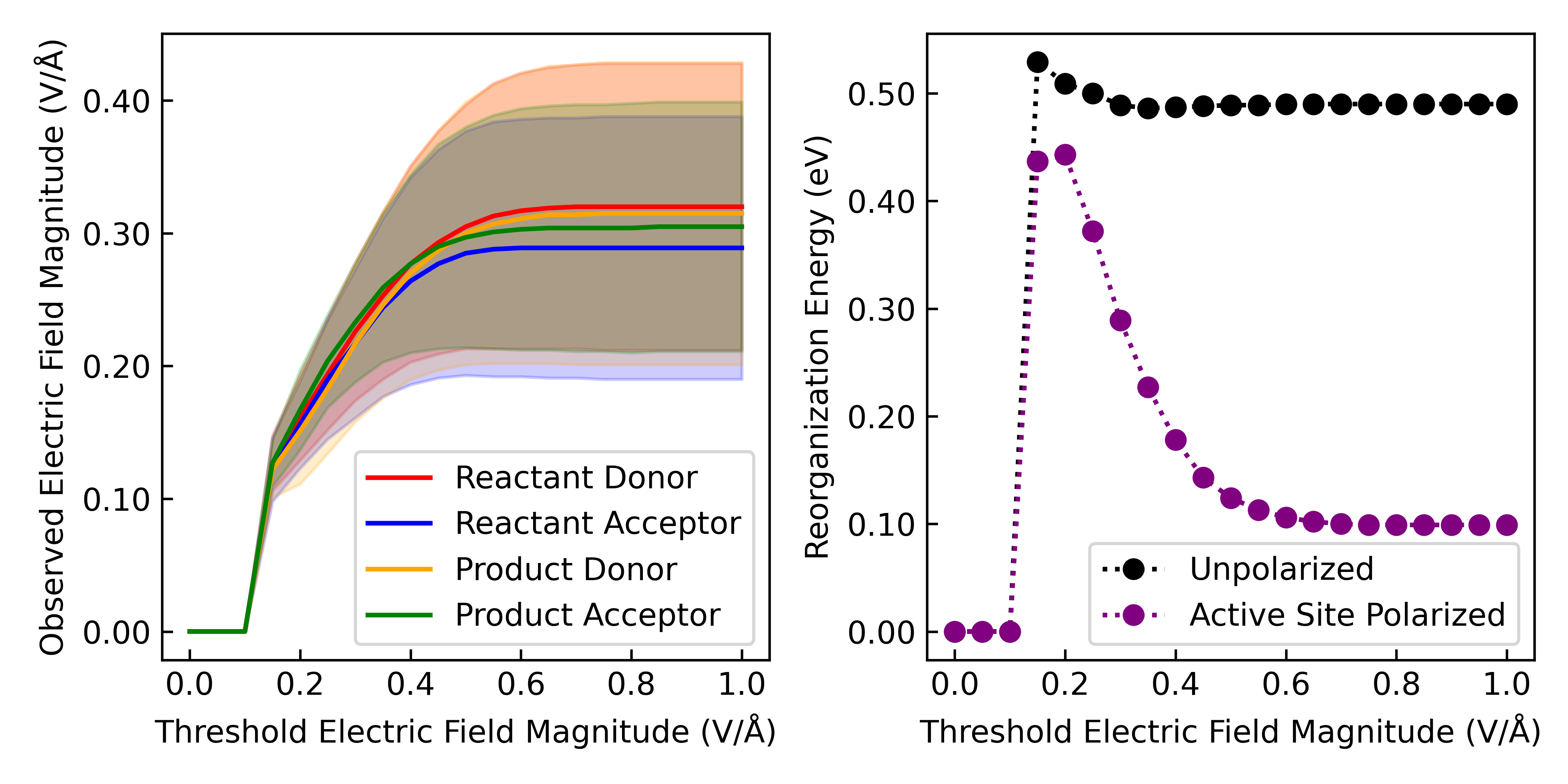

Figure S27. A wide distribution of electric field magnitudes exerted on the Fe centers of the heme groups in OmcS induces a significance decrease in the outer-sphere reorganization energy. The average magnitude and standard deviation of the field on the donor and acceptor in both states (*left*) and the reorganization energy (*right*) are plotted as a function of the threshold field magnitude used to filter the configurations from molecular dynamics. The data is shown for electron transfer between Hemes #6 and #1’.

### Electron Transfer Energetics in All Structurally Characterized Cytochrome Nanowires

Table S12. Activation energies and electronic couplings computed for all structurally characterized cytochrome ‘nanowires’ as of July 2024 using quantum mechanics/molecular mechanics at molecular dynamics-generated configurations (QM/MM@MD) and BioDC protocols. Importantly: (1) the electronic coupling $\sqrt{\left\langle H^{2} \right\rangle}$ in the BioDC protocol is a parameter that assigns the pre-defined values 2.0 or 8.0 meV for perpendicular and parallel heme pairs. (2) The reorganization energies that enter $E_{a}$ for the QM/MM@MD approach come from non-polarizable molecular dynamics and were therefore scaled down by 0.56. The reorganization energies that enter $E_{a}$ for the BioDC approach were computed from the minimized structure using the parameterized form of the Marcus continuum expression based on solvent accessibility developed by Blumberger and co-workers that is intended to reproduce $\lambda$ from polarizable molecular dynamics; these $\lambda$s were not scaled, even though they are consistently larger than the scaled ones used for QM/MM@MD.

|  | QM/MM@MD | | BioDC | |
| --- | --- | --- | --- | --- |
| Heme Index | $E_{a}$ (eV) | $\langle H\rangle$ (meV) | $E_{a}$ (eV) | $\langle H\rangle$ (meV) |
| A3MW92 (PDB 8E5F) | | | | |
| 1$\leftrightarrow$2 |  |  | 0.166 | 2.000 |
| 2$\leftrightarrow$3 |  |  | 0.222 | 8.000 |
| 3$\leftrightarrow$4 |  |  | 0.133 | 2.000 |
| 4$\leftrightarrow$1’ |  |  | 0.202 | 8.000 |
| F2KMU8 (PDB 8E5G) | | | | |
| 1$\leftrightarrow$2 |  |  | 0.207 | 2.00 |
| 2$\leftrightarrow$3 |  |  | 0.112 | 8.00 |
| 3$\leftrightarrow$4 |  |  | 0.191 | 2.00 |
| 4$\leftrightarrow$1’ |  |  | 0.146 | 8.00 |
| OmcE (PDB | | | | |
| 1$\leftrightarrow$2 | 0.105 | 1.472 | 0.143 | 2.00 |
| 2$\leftrightarrow$3 | 0.063 | 13.001 | 0.280 | 8.00 |
| 3$\leftrightarrow$4 | 0.065 | 1.775 | 0.189 | 2.00 |
| 4$\leftrightarrow$1’ | 0.145 | 4.411 | 0.154 | 8.00 |
| OmcS (PDB 6EF8) | | | | |
| 1$\leftrightarrow$2 | 0.220 | 1.13 | 0.274 | 2.00 |
| 2$\leftrightarrow$3 | 0.032 | 6.09 | 0.206 | 8.00 |
| 3$\leftrightarrow$4 | 0.182 | 2.64 | 0.066 | 2.00 |
| 4$\leftrightarrow$5 | 0.088 | 9.52 | 0.106 | 8.00 |
| 5$\leftrightarrow$6 | 0.065 | 1.04 | 0.241 | 2.00 |
| 6$\leftrightarrow$1’ | 0.072 | 7.90 | 0.132 | 8.00 |
| OmcZ (7LQ5) | | | | |
| 1$\leftrightarrow$2 | 0.137 | 2.04 | 0.334 | 2.00 |
| 2$\leftrightarrow$3 | 0.120 | 3.19 | 0.166 | 8.00 |
| 3$\leftrightarrow$4 | 0.220 | 7.15 | 0.042 | 8.00 |
| 4$\leftrightarrow$5 | 0.088 | 4.06 | 0.077 | 2.00 |
| 5$\leftrightarrow$6 | 0.084 | 5.21 | 0.167 | 8.00 |
| 6$\leftrightarrow$7 | 0.010 | 2.32 | 0.183 | 2.00 |
| 7$\leftrightarrow$1’ | 0.139 | 4.71 | 0.224 | 8.00 |
